# Multi-cohort analysis of 37,739 oral microbiomes reveals ecologically influential health-associated microbial sub-communities across major oral subsites

**DOI:** 10.64898/2026.08.16.745142

**Authors:** Omprakash Shete, Alisha Ansari, Mithun Verma, Abhishek P, Esha Chauhan, Sourav Goswami, Tarini Shankar Ghosh

**Affiliations:** Department of Computational Biology, Indraprastha Institute of Information Technology (IIIT-Delhi), Okhla Phase III, New Delhi, India

**Keywords:** subsite-resolved oral microbiome, health-associated core taxa, microbial modules, potential diagnostic signature, sHACK/HAC

## Abstract

The oral cavity contains multiple microbial sub-niches, but which taxa consistently play an ecologically important, health-associated role within each niche, and how conserved they are across populations, remains poorly understood, partly due to the lack of a standardised identification framework.

We developed a multi-cohort framework integrating 37,739 oral microbiome profiles (16S rRNA and shotgun sequencing) from 142 cohorts (41 countries) ranking 542 taxa across four oral habitats, supragingival, subgingival, tongue–tonsil, and buccal–palate-mucosa, via a new Health-Associated-Core (HAC) score capturing consistent prevalence, ecological influence, and health-association. For saliva, with available longitudinal sampling, we extended this into a salivary-Health-Associated-Core-Keystone (sHACK) score additionally capturing stability-association, ranking 499 taxa.

Using two complementary approaches for identifying ecological modules, high-sHACK salivary taxa concentrated within a single, connected sub-community of 28 members, consistently linked to prevalence, ecological influence, stability, and health. This sub-community’s abundance alone outperformed conventional dysbiosis indices in distinguishing healthy from diseased individuals and tracked stability in an independent cohort of 4,621 microbiomes. Comparable sub-communities emerged across three other subsites, with compositional differences mirroring physicochemical variation between sites.

Machine learning linked taxa-specific-genome-encoded functions to their corresponding subsite-specific HAC/sHACK scores, offering a unified framework for prioritizing oral microbes diagnostically and therapeutically.

## Introduction

The human oral cavity harbours one of the most diverse microbial ecosystems in the body, comprising hundreds of bacterial species distributed across distinct anatomical niches, including saliva, dental plaque, the tongue, and mucosal surfaces^1,2^. These communities interact extensively with host tissues and contribute to immune regulation, metabolism, and colonization resistance^2,3^. Disruptions in oral microbial community structure have been linked to numerous local and systemic diseases, including periodontitis, dental caries, oral cancer, cardiovascular disease, inflammatory bowel disease, and metabolic disorders^2,4–7^. These associations highlight the potential of the oral microbiome as a target for diagnostic and therapeutic interventions. Realising this potential, however, requires a robust understanding of the microbial features that define healthy and stable oral ecosystems.

Most oral microbiome studies have focused on describing community composition within individual cohorts or identifying taxa associated with specific diseases^8–13^. As a result, it remains unclear which oral microbes consistently contribute to ecological organization, community stability, and host health across populations. In ecological systems, keystone species exert disproportionate effects on community structure and resilience. Recent large-scale meta-analyses of the gut microbiome have demonstrated that integrating ecological influence, longitudinal stability, and disease associations can systematically identify microbial keystones at population scale^14^. Comparable frameworks have not yet been applied to the oral microbiome, particularly in ways that account for the substantial heterogeneity arising from sequencing technologies, study designs, and disease contexts.

A further challenge is the pronounced ecological heterogeneity of the oral cavity. Oral habitats differ markedly in oxygen availability, nutrient sources, host immune interactions, and epithelial turnover, resulting in distinct microbial communities across supragingival plaque, subgingival plaque, the tongue, saliva, and mucosal surfaces^11,15–21^. Ecological processes such as microbial succession, metabolic cross-feeding, and spatial biofilm organization reinforce this niche specialization^15,22–24^. Although oral habitat-specific community structure is well established, it remains unknown whether common health-associated keystone taxa and signatures of microbiome stability conserved across population-groups. How do these keystone signatures vary across oral subsites? Consequently, identifying globally relevant health-associated keystone taxa remains a major challenge in oral microbiome research.

To address this gap, we performed a large-scale, subsite-resolved meta-analysis of 37,739 publicly available oral microbiome profiles from 142 cohorts to identify health-associated and ecologically important taxa across five major oral habitats. Building on a previously developed keystone-identification framework^14^, we quantified the consistency of taxa associations with prevalence in healthy microbiomes, ecological influence, and host health, and integrated these properties into subsite-specific Health-Associated Core (HAC) scores. For saliva-associated microbiomes, where longitudinal data were available, we further incorporated microbial associations with temporal stability to derive a salivary Health-Associated Core-Keystone (sHACK) score.

Beyond generating oral taxa rankings, we identified conserved ecological modules, or sub-communities, within each habitat using two complementary strategies: one grouping taxa by their shared association patterns with the rest of the community, and the other based on direct pairwise co-occurrence across cohorts. We observed specific ecological modules within each oral sub-niche that were selectively enriched for the high-HAC/high-sHACK taxa. For saliva-associated microbiomes, high-sHACK salivary taxa clustered within a single, interconnected sub-community of 28 members. The abundance of this sub-community alone surpassed conventional dysbiosis indices in discriminating healthy from diseased individuals, and reliably captured stability in an independent cohort of 4,621 microbiomes. Similar sub-communities were observed in three other habitats, with compositional variation reflecting underlying physicochemical differences between sites. Finally, using machine-learning, we identified the genomic-functional-hallmarks of the high HAC/sHACK taxa across subsites. By jointly evaluating ecological influence, health-association, and persistence, this framework prioritizes taxa that underpin microbiome resilience across oral habitats.

## Results

### Creation of a global subsite-resolved oral microbiome repository and identification of consensus oral-associated taxa

Identifying microbial determinants of health and microbiome resilience across different oral sub-ecosystems first required building a large repository of global oral microbiome datasets. Using extensive literature and keyword-based searches on the European Nucleotide Archive (ENA) database^25^, we curated and processed oral microbiome datasets from 142 study-cohorts, encompassing 37,739 oral microbiomes from individuals aged ≥12 years across 41 countries. This dataset comprised 31,988 16S rRNA amplicon (16S) profiles and 5,751 whole-genome shotgun sequencing (WGS) profiles (**Table S1**). WGS and 16S data were processed using MetaPhlAn3 and SPINGO, respectively, to ensure consistent taxonomic profiling^26,27^.

The oral microbiomes were sampled from various subsites within the oral environment. Given the strong variation in micro-environments across oral subsites, and to enable a subsite-resolved analysis, all samples were harmonised into five major oral subsite categories: saliva-sputum-oral wash (hereafter salivary), supragingival, subgingival, tongue-tonsil-related sites, and buccal-palate-other surface sites (See **Methods**; **Table S1** for detailed subsite-wise cohort information). Categorization was performed by manually mapping author-reported sampling terms to standardised oral habitat categories (**Table S2**). Amongst these categories, the salivary category was largest, encompassing 22,949 microbiomes from 124 study-cohorts. Of these, 105 cohorts (18,800 microbiomes, 28 countries) were investigated in the discovery phase, and 19 cohorts (9 countries) encompassing 4,149 microbiomes in the validation phase, to validate patterns learnt in discovery. The discovery sub-group included 15,244 and 3,556 microbiomes from non-diseased (control) and diseased individuals, respectively (**Table S1A**), including 2,692 longitudinally sampled microbiomes. The validation sub-cohort included 2,885 and 1,217 microbiomes from control and diseased individuals, respectively, of which 1,525 were longitudinal (**Table S1F**). Supragingival formed the second largest group, with 10,381 samples (**Table S1B**), followed by subgingival (n = 1,974) (**Table S1C**), tongue-tonsil-related sites (n = 1,693) (**Table S1D**), and buccal-palate-other surface sites (n = 742) (**Table S1E**).

The geographical and subsite-wise distribution of the curated repository is summarised in **Figure 1A**. Although samples were collected from 41 countries, the largest representation came from the United States, followed by China, indicating that a substantial fraction of currently available oral microbiome data originates from industrialised-urban populations. This curated, subsite-resolved data resource provided the basis for all downstream analyses within each subsite (subject-specific metadata in **Table S3**).

**Figure 1.**
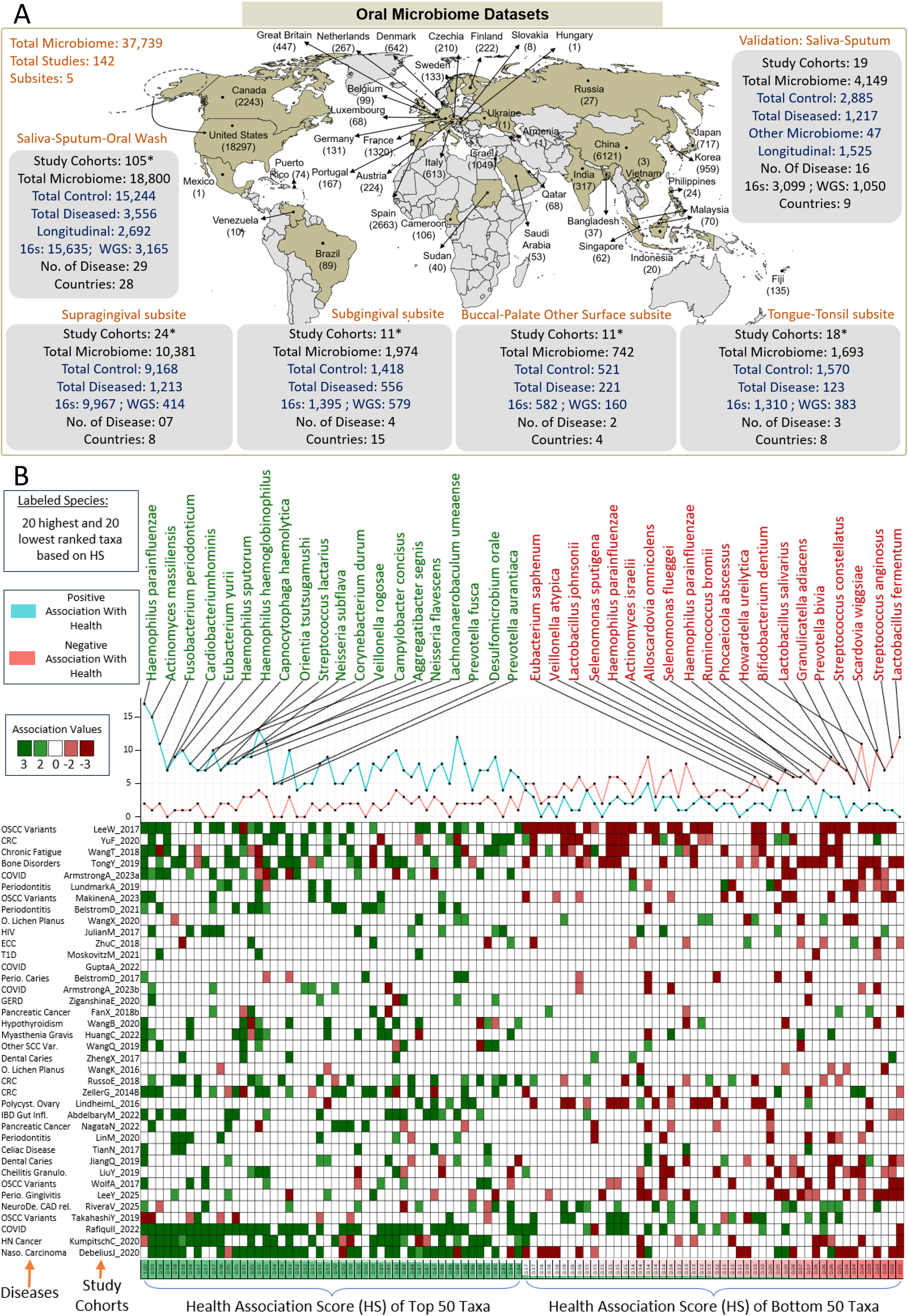
Global oral microbiome repository and salivary health-association patterns. **(A) Global distribution and composition of the curated oral microbiome repository**, comprising 37,739 oral microbiomes from 142 study-cohorts across 41 countries and five oral subsites: saliva-sputum-oral wash (abbreviated as salivary), supragingival, subgingival, tongue-tonsil, and buccal-palate-other-surface subsites. Subsite-wise boxes summarize study-cohort numbers, sample counts, disease-control status, sequencing strategy, disease categories, and country representation. Country-wise microbiome counts are shown on the map. For the salivary subsite, summary of the validation cohort is shown separately. Asterisks indicate study-cohort counts after splitting datasets by sequencing strategy, subsite, or exposure status. **(B) Study-cohort-level health-association patterns of salivary taxa**. Shown using a heatmap. Columns represent matched case-control salivary study-cohorts, and rows represent the top 50 and bottom 50 taxa ranked by Health-Association Score (HS). Each cell indicates whether a taxon was control-associated, disease-associated, or non-significant within a given cohort. Disease categories corresponding to each study-cohort are annotated on the left side of the heatmap. The upper line plot summarizes recurrent control- and disease-associated signals across cohorts, while labelled taxa indicate the top 20 and bottom 20 taxa by HS.

To prevent downstream investigations from being influenced by random taxonomic noise arising from sparsely detected taxa, we identified consensus sets of subsite-associated taxa for each oral subsite, by jointly optimizing the detection rate of taxa-groups across microbiomes from that subsite and their cumulative representation across all contributing study-cohorts (**Text S1**). This approach yielded 499, 301, 196, 266, and 166 consensus taxa for the salivary, supragingival, subgingival, tongue-tonsil, and buccal-palate-other-surface subsites, respectively (**Figures S1-S5**; **Table S4**), used for all downstream analyses.

We began by focusing on the salivary subsite, which contained the largest number of microbiomes, including longitudinally sampled cohorts, enabling assessment of taxa based on the consistency of their association with three key properties relevant for identifying microbes important for health and microbiome resilience: health-association, ecological influence, and association with longitudinal microbiome stability.

### Salivary taxa could be ranked in a specific order based on the consistency of their association with health across the study-cohorts

We first investigated the consistency of health-association across salivary taxa. Of the 105 discovery-phase cohorts, 38 included salivary microbiomes (N = 5,466) from matched diseased (N = 2,840) and control (N = 2,626) sub-groups, spanning 26 disease categories, grouped into broad classes (**Table S5A-B**).

For each cohort, we compared taxon abundances between diseased and matched-control sub-groups (Mann-Whitney test), treating significant depletion in disease as a positive health-association. *Haemophilus parainfluenzae* was most consistently health-associated, depleted in disease across 17 cohorts, followed by *Actinomyces massiliensis* (15) and *Fusobacterium periodonticum* (11), each enriched in disease in fewer than two cohorts (**Figure 1B**). *Cardiobacterium hominis*, *Eubacterium yurii*, *Haemophilus sputorum*, *Streptococcus lactarius*, *Corynebacterium durum*, *Veillonella rogosae*, *Neisseria subflava*, and multiple *Prevotella*, *Kingella*, and *Lachnoanaerobaculum* taxa were similarly depleted in disease (p ≤ 0.05) in over 10% of cohorts, with enrichment in no more than two. Conversely*, Lactobacillus fermentum*, *Streptococcus anginosus*, *Scardovia wiggsiae*, *Streptococcus constellatus*, *Granulicatella adjacens*, and *Bifidobacterium dentium* showed the opposite, more consistent enrichment in disease (**Figure 1B**). Salivary taxa thus varied markedly in health-association consistency across cohorts and diseases, suggesting a specific ranking order.

In our earlier gut-microbiome study identifying HACKs^14^, we developed a scheme scoring taxa by consistency of health-association across cohorts, classifying each taxon per cohort as control-, disease-associated, or non-significant, then converting this recurrence into a normalised score ranking all considered taxa. We applied this to compute Health-Association scores (HS) for 499 salivary taxa (**Text S2**; **Figure S6**; **Table S6A**), which captured the patterns above (**Figure 1B**).

To assess reproducibility, we recomputed stratified HS within cohort subsets by sequencing type (16S, WGS) and exposure status (**Table S6B**). Overall HS correlated significantly with HS computed specifically within 16S, WGS, and exposure-related cohorts (16S vs. overall: R = 0.88, p = 1.4e-153; WGS vs. overall: R = 0.38, p = 2.7e-9; exposure vs. overall: R = 0.34, p = 3.9e-13) (**Figure S7**). Most consistently health-associated taxa noted above, including *H. parainfluenzae*, *F. periodonticum*, *A. massiliensis*, *C. haemolytica, C. durum*, *V. rogosae*, *N. subflava*, and specific *Prevotella*, *Oribacterium*, and *Lachnoanaerobaculum* taxa, remained in the top 30% across all stratified analyses (**Figure S7**), indicating overall HS reliably captures health-association patterns even within distinct cohort subcategories.

### Integrating health-, core-, and stability-association identifies candidate keystone taxa of the salivary microbiome

Core-Association reflects how consistently a taxon is a core member of the salivary microbiome across cohorts, defined jointly by high prevalence in non-diseased individuals and strong association with community-wide compositional variability (ranked R² contribution to beta-diversity). Applying this to 15,127 control microbiomes from 99 cohorts (**Table S7**, **Text S2**), we identified core-associated taxa per cohort using an optimised prevalence threshold of 0.85 and R² rank threshold of 0.75 for the salivary subsite (**Text S2**; **Figure S8A-B**), within our previously developed 3R framework^14^. We quantified this with a Core-Association score (CS) for each taxon, defined as the proportion of control cohorts where a taxon was core-associated, rank-scaled between 0 and 1 (**Figure S8C**).

The ranking was dominated by *Prevotella*, *Veillonella*, *Actinomyces*, *Streptococcus*, *Fusobacterium*, *Rothia* and *Capnocytophaga* (**Figure S9**; **Table S7**), notably including *F. periodonticum*, *H. parainfluenzae* and *Neisseria, Actinomyces* lineage members already identified as strongly health-associated (**Figure 1B**), pointing to taxa simultaneously health-associated, core, and ecologically well-connected. SStratified CS within cohort types (16S, WGS, Exposure) showed strong reproducibility, correlating significantly with each other and with overall CS (**Figure S10**, **Table S8A**) (16S v/s Overall: R = 0.98, p = 6.2e-165; WGS v/s Overall: R = 0.44, p = 1.1e-5; Exposure v/s Overall: R = 0.76, p = 3.9e-28).

Comparing CS and HS across 499 salivary taxa divided them into groups. The most important, from our study’s perspective, showed simultaneously high CS and HS (≥0.70), defining microbes both reproducibly core-associated and consistently health-associated (Quadrant1) (**Figure 2A**; **Table S8B**), spanning *Prevotella* (*P. pallens*, *P. melaninogenica*, *P. veroralis*, *P. shahii*) *Alloprevotella rava*, *Fusobacterium periodonticum*, *Leptotrichiaceae* (*Leptotrichia buccalis*, *L. hofstadii*), *Campylobacter* (*C. concisus*, *C. showae*), *Neisseria* (*N. elongata*, *N. flavescens*, *N. mucosa*), *Haemophilus* (*H. parainfluenzae*, *H. sputorum*, *H. pittmaniae*), *Cardiobacterium* (*C. hominis*, *C. valvarum*), *Flavobacteriaceae* (*Capnocytophaga gingivalis*), *Lachnoanaerobaculaceae* (*Lachnoanaerobaculum umeaense*, *Catonella morbi*), *Porphyromonas* (*P. catoniae*), *Actinomyces* (*A. graevenitzii*), and *Eubacterium* (*E. sulci*, *E. yurii*). Not all taxa scored high on both properties; we also observed groups with high CS/low HS, high HS/low CS, and low on both (**Figure 2A**; **Table S8B**; **Text S3**). This Q1 group highlights potential hallmarks for measuring oral microbiome health and guiding diagnostics/therapeutics, though an ideal hallmark should also satisfy a third property: association with longitudinal stability (Stability-Association Score; SS).

**Figure 2.**
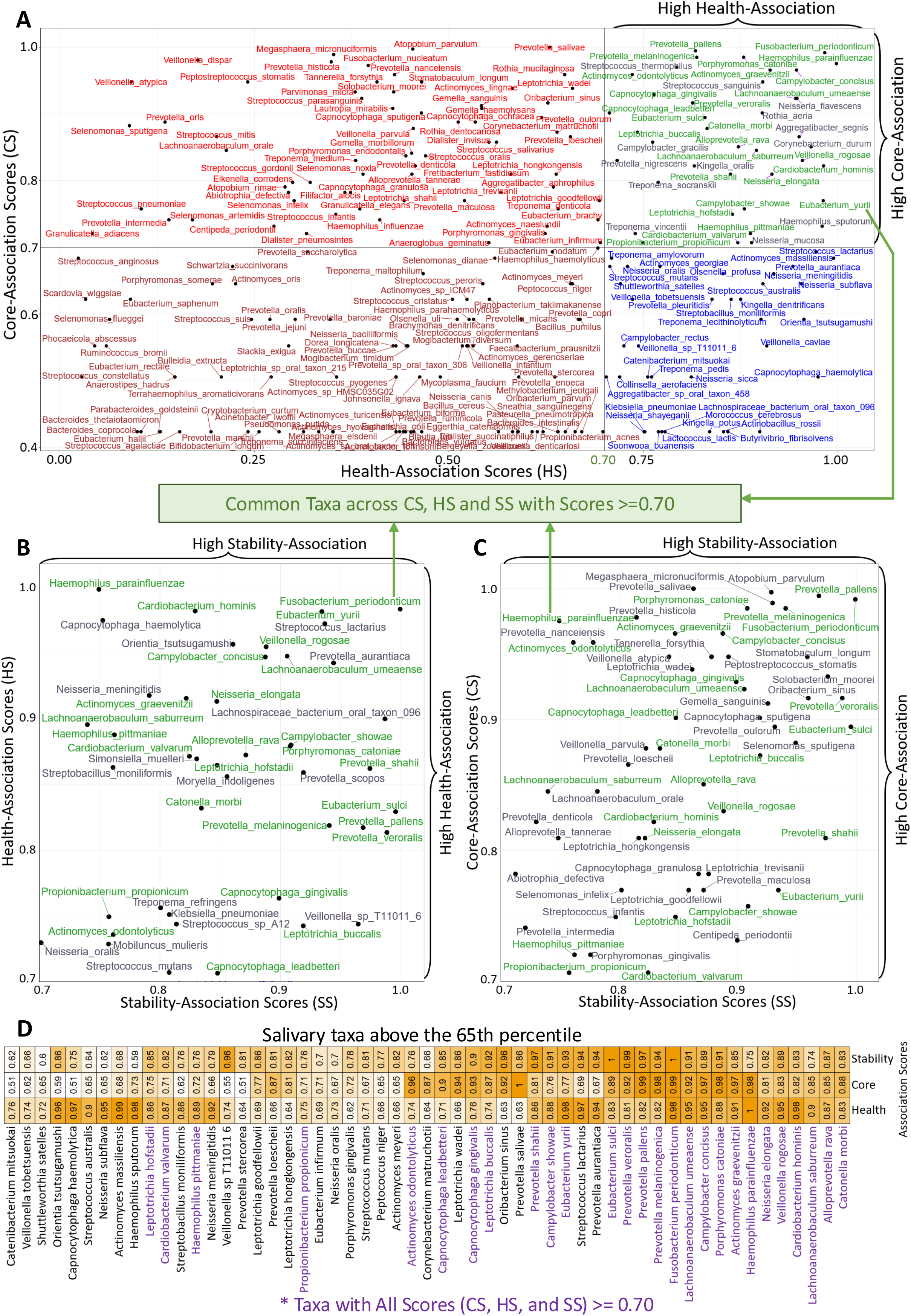
Concordance of health, core, and stability association scores in salivary taxa. **(A) Scatter plot comparing Core-Association Score (CS) and Health-Association Score (HS) for salivary taxa**. Only taxa with non-zero CS values are shown. A score threshold of 0.70 was used to stratify taxa into quadrants and identify taxa that were both recurrently core-associated across control cohorts and consistently health-associated across matched case-control cohorts. **(B) Scatter plot comparing Stability Association Score (SS) and Health-Association Score (HS) for salivary taxa.** Only taxa with both SS and HS ≥ 0.70 are shown, highlighting species associated with reduced longitudinal microbiome change and health-associated disease-control directionality. **(C) Scatter plot comparing Stability-Association Score (SS) and Core-Association Score (CS) for salivary taxa.** Only taxa with both SS and CS ≥ 0.70 are shown, identifying species that combined recurrent core-association with association to longitudinal microbiome stability. For **A-C**: Green labels indicate taxa common across all three scoring dimensions with CS, HS, and SS ≥ 0.70. **(D) Heatmap showing salivary taxa within the top 35 percentile for Core-Association Score, Health-Association Score, and Stability-Association Score.** Values indicate rank-scaled scores for each taxon across the three dimensions. Taxa with all three scores ≥ 0.70 are highlighted, representing species with concordant core, health, and stability associations.

To grade taxa on this, we used 2,692 salivary microbiomes from 12 cohorts with follow-up sampling from 726 subjects (**Table S9**). In this investigation, we first computed “follow-up distances” for each microbiome (with an available follow-up sample) as the variation between a microbiome’s current and follow-up state as an inverse measure of stability. The 499 taxa were then scored for association with longitudinal microbiome stability, using a previously developed iterative meta-analytic framework measuring the extent and consistency of negative association with follow-up distances (**Test S2**; **Figure S11**).

Several taxa were consistently associated with reduced longitudinal change (**Figure S12**). Stratified SS rankings correlated significantly across cohort-types and with overall SS (16S v/s Overall: R = 0.93, p = 1.5e-208; WGS v/s Overall: R = 0.67, p = 6.4e-32) (**Figure S13**; **Table S10A**). Consistent stability-associated taxa included *Fusobacterium periodonticum*, *Eubacterium sulci*, *Prevotella veroralis*, *Lachnospiraceae bacterium oral taxon 096*, *Solobacterium moorei* and multiple Prevotella, Veillonella, Oribacterium, Capnocytophaga and Stomatobaculum members. We identified 27 taxa with all three scores in the top 30% (**Figure 2A-C**; green taxa), namely *Actinomyces graevenitzii*, *A. odontolyticus*, *Alloprevotella rava*, *Campylobacter concisus*, *C. showae*, *Capnocytophaga gingivalis*, *C. leadbetteri*, *Cardiobacterium hominis*, *C. valvarum*, *Catonella morbi*, *Eubacterium sulci*, *E. yurii*, *Fusobacterium periodonticum*, *Haemophilus parainfluenzae*, *H. pittmaniae*, *Lachnoanaerobaculum saburreum*, *L. umeaense*, *Leptotrichia buccalis*, *L. hofstadii*, *Neisseria elongata*, *Porphyromonas catoniae*, *Prevotella melaninogenica*, *P. pallens*, *P. shahii*, *P. veroralis*, *Propionibacterium propionicum* and *Veillonella rogosae* (**Figure 2D**; **Table S10B**).

### Majority of the salivary health-associated core keystone taxa concentrate within a single, co-abundant microbial sub-community

We combined each taxon’s core-association, stability-association, and health-association scores into a single index, the salivary-Health-Associated-Core-Keystone (sHACK) index (**Figure 3A**; **Text S4**; **Table S11**). As in our previous gut microbiome study^14^, ‘health-associated’ reflects consistency of association with healthy individuals across cohorts; ‘Core’ reflects consistency with which a taxon constitutes the prevalent salivary microbiome in healthy individuals; and ‘keystone’ reflects ecological influence together with consistent association with longitudinal stability. The top 10% of sHACK (n = 50) included all 27 taxa with all three scores ≥0.70, plus lineages from Prevotella, Neisseria, Leptotrichia, Oribacterium, Streptococcus, and Corynebacterium clades (**Figure 3B**). sHACK recalculated within cohort subsets (16S and WGS) correlated strongly with overall scores (16S v/s Overall: R = 0.96, p = 5.1e-259; WGS v/s Overall: R = 0.54, p = 6.9e-19) (**Figure S14**), confirming reproducibility across profiling strategy, cohort composition, and exposure.

**Figure 3.**
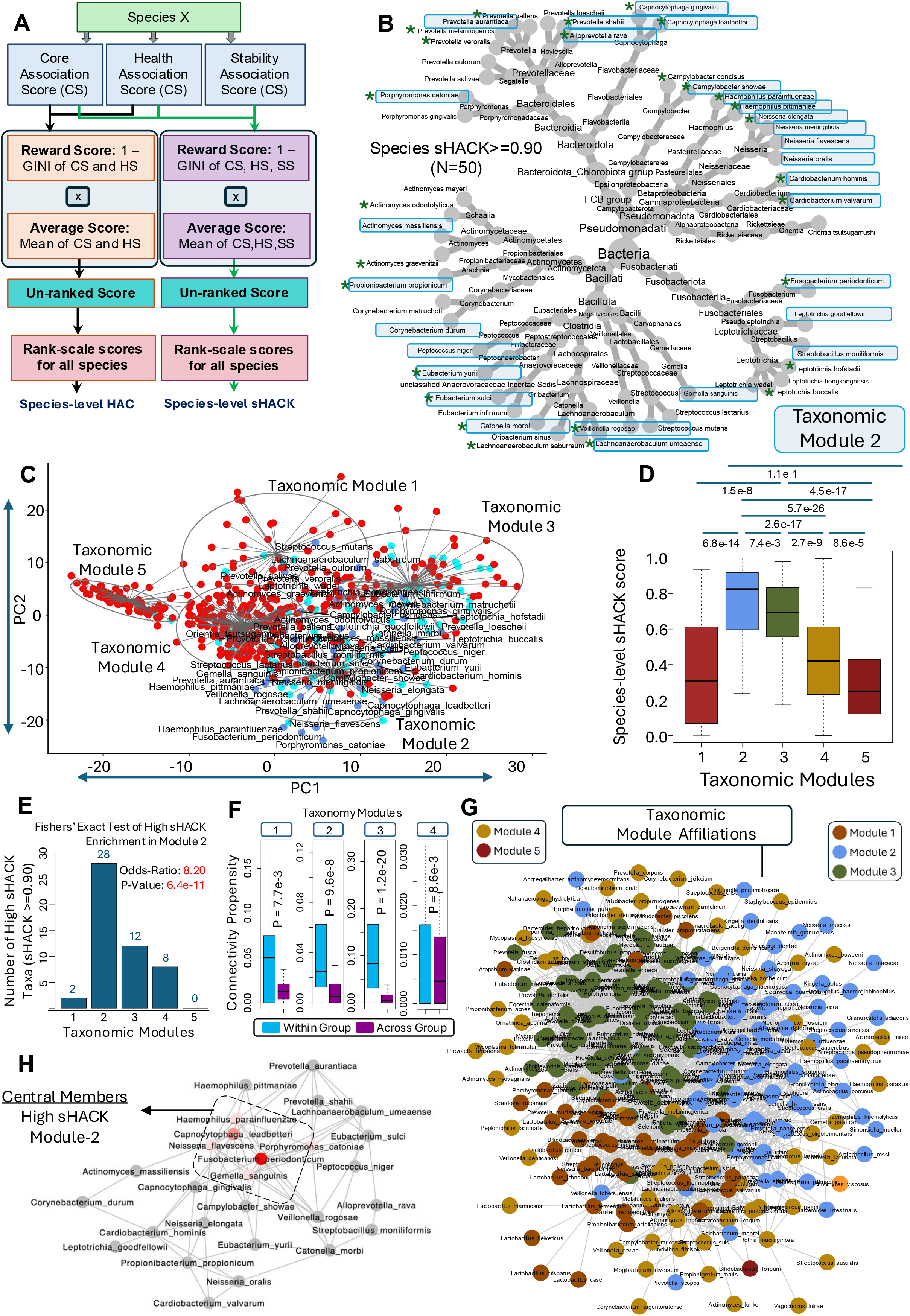
Identification of health-associated core-keystone taxa in the salivary microbiome and their modular organization. **A.** Schematic of salivary Health-Associated Core Keystone (sHACK) score computation, integrating each taxon’s Core-Association (CS), Health-Association (HS), and Stability-Association (SS) scores using a Gini-based concordance term. A separate Health-Associated Core (HAC) score was computed using the same strategy but integrating only CS and HS. HAC scores were calculated for taxa across all five subsites, including saliva, whereas sHACK scores were restricted to the salivary subsite, the only site with available longitudinal data required to compute SS. **B.** Circular phylogenetic tree of top 50 high-sHACK taxa (sHACK ≥ 0.90), showing their distribution across multiple bacterial lineages. Taxa belonging to taxonomic module 2 (described below) are highlighted in blue boxes while taxa with all three scores (CS, HS, SS) >= 0.70 are shown with asterisk. **C.** PCA of salivary taxa based on community-wide taxon–taxon association profiles, identifying five taxonomic modules. Taxa are coloured by sHACK score category: cornflower blue, sHACK ≥ 0.90; cyan, 0.75 ≤ sHACK < 0.90; red, sHACK < 0.75. Module 2 shows enrichment of high-sHACK taxa. The detailed investigation pertaining to the identification of these modules is described in **Text S5** and **Figure S15**. **D.** Distribution of sHACK score across the five taxonomic modules, showing higher sHACK scores in module 2. **E.** Number of high-sHACK taxa (sHACK score ≥ 0.90) in each module. Module 2 showed significant enrichment of high-sHACK taxa by Fisher’s exact test (odds ratio = 8.20, P = 6.4e-11). **F.** Within- and across-module connectivity propensity for taxa belonging to different modules in the co-abundance meta-network shown in **G**, showing significantly higher within-module connectivity for the high sHACK module-2, and the modules 1 and 3. **G.** Co-abundance meta-network of salivary taxa, with nodes coloured by taxonomic module (details in **Text S5**). Modules 2 and 3 form visibly cohesive network regions, indicating stronger co-abundance among taxa within these modules. **(H)** Sub-network of 28 high-sHACK (sHACK score >= 0.90) taxa belonging to module 2, highlighting central members within the salivary health-associated core-keystone (sHACK) sub-community.

Health-association and ecological influence identify individual taxa likely to contribute to oral health but do not reveal how they are organised within the community. A key question is whether health-associated taxa assemble into conserved co-occurring modules forming a core ecological architecture, or act independently. Resolving this offers a systems-level understanding of microbiome organization and a basis for designing therapeutic consortia. We tested this using a two-step framework.

We first clustered salivary taxa by their association patterns with the rest of the community, computing pairwise Association Scores between each of 499 taxa and all others using non-diseased discovery-cohort microbiomes, following our previous approach^28^ (**Text S5**, **Figures S15A-B**). PCA positioned taxa in two-dimensional space (PC1, PC2), with taxa sharing similar association patterns clustering together (**Table S12**, **Figure S15A**). k-means clustering, with cluster number optimised via Calinski-Harabasz index across iterative subsampling (**Text S5**), partitioned taxa into five modules based on 15,127 non-diseased microbiomes across 99 cohorts (**Figure S15B**; **Table S12**). Modules 2 and 3 showed significantly higher CS than others, and module 2 also showed the highest SS and HS (**Figure S15C-E**), mirrored in sHACK, where module 2 taxa scored significantly higher than all others (**Figure 3C-D**). Strikingly, 28 of the 50 high-sHACK taxa (sHACK ≥0.90) belonged to module 2 alone (odds ratio = 8.20; Fisher’s exact test, p = 6.4e-11) (**Figure 3E**), as did 18 of the 27 top-30% taxa (**Figure 3B**). Thus, sHACK taxa concentrate within a single sub-community comprising phylum *Pseudomonadati* (earlier *Proteobacteria*) (*Haemophilus*, *Neisseria*, *Cardiobacterium*), phylum *Bacillota* (earlier *Firmicutes*) (*Gemella*, *Veillonella*, *Lachnoanaerobaculum*, *Catonella*, order *Peptostreptococcales*), and *Fusobacterium periodonticum* (**Figure 3B**). We next tested whether this sub-community is a genuine ecological unit by examining whether its taxa also showed elevated co-occurrence relative to taxa outside the module.

Thus, we next built a co-abundance-based taxon-to-taxon meta-network across non-diseased salivary microbiomes (**Text S5**): for each taxon pair, we performed a random-effects meta-analysis of co-abundance across cohorts, estimating effect size per cohort via robust linear regression and pooling estimates to obtain a summarised effect, p-value, and consistency (fraction of cohorts matching the pooled direction). After FDR correction, edges were drawn for pairs with a positive summarised estimate, Q ≤0.0001, and consistency ≥0.70, yielding a discovery-cohort meta-network of 2,420 edges across 283 taxa (**Table S13**; **Figure 3F-G**).

We then tested whether taxa within each module were preferentially connected to one another. Modules 2 and 3 visibly co-localised in the meta-network (**Figure 3G**). For each taxon, we calculated within- and across-module edge-propensity, compared using paired Wilcoxon signed-rank tests. Modules 1, 2, and 3 showed significantly higher within- than across-module edge-propensity (**Figure 3F**; p = 7.7e-3, 9.6e-8, and 1.2e-20, respectively), confirming these modules are genuine ecological units rather than clustering artifacts.

The sub-network of high-sHACK taxa within module 2 showed strong co-abundance centered on *Fusobacterium periodonticum*, *Capnocytophaga leadbetteri*, *Gemella sanguinis*, *Haemophilus parainfluenzae*, *Neisseria flavescens*, and *Porphyromonas catoniae*, which appeared to drive connectivity within this sub-community (**Figure 3H**). Thus, the predominant HACKs of the salivary microbiome originate from a single, biologically co-occurring ecological unit.

### Ecological arrangement of the salivary microbiome observed in the discovery cohort is also reproduced in validation cohort of 4,149 unseen oral microbiomes

We asked whether the taxonomic modules identified in the discovery cohort reflected a universal feature of the salivary microbiome or were specific to the cohort composition used to define them, using an additional ‘unseen’ validation dataset of 19 study-cohorts (4,149 salivary microbiomes). Using 2,885 non-diseased microbiomes from this cohort, we built Association Maps (as in **Text S5**) (**Table S14**). Co-localization patterns closely resembled the discovery cohort (Procrustes: R = 0.93, p = 1.0e-3) (**Figure 4A**), with taxa from the same discovery-defined modules localizing to similar regions in the validation map, indicating similar ecological arrangements regardless of dataset.

**Figure 4.**
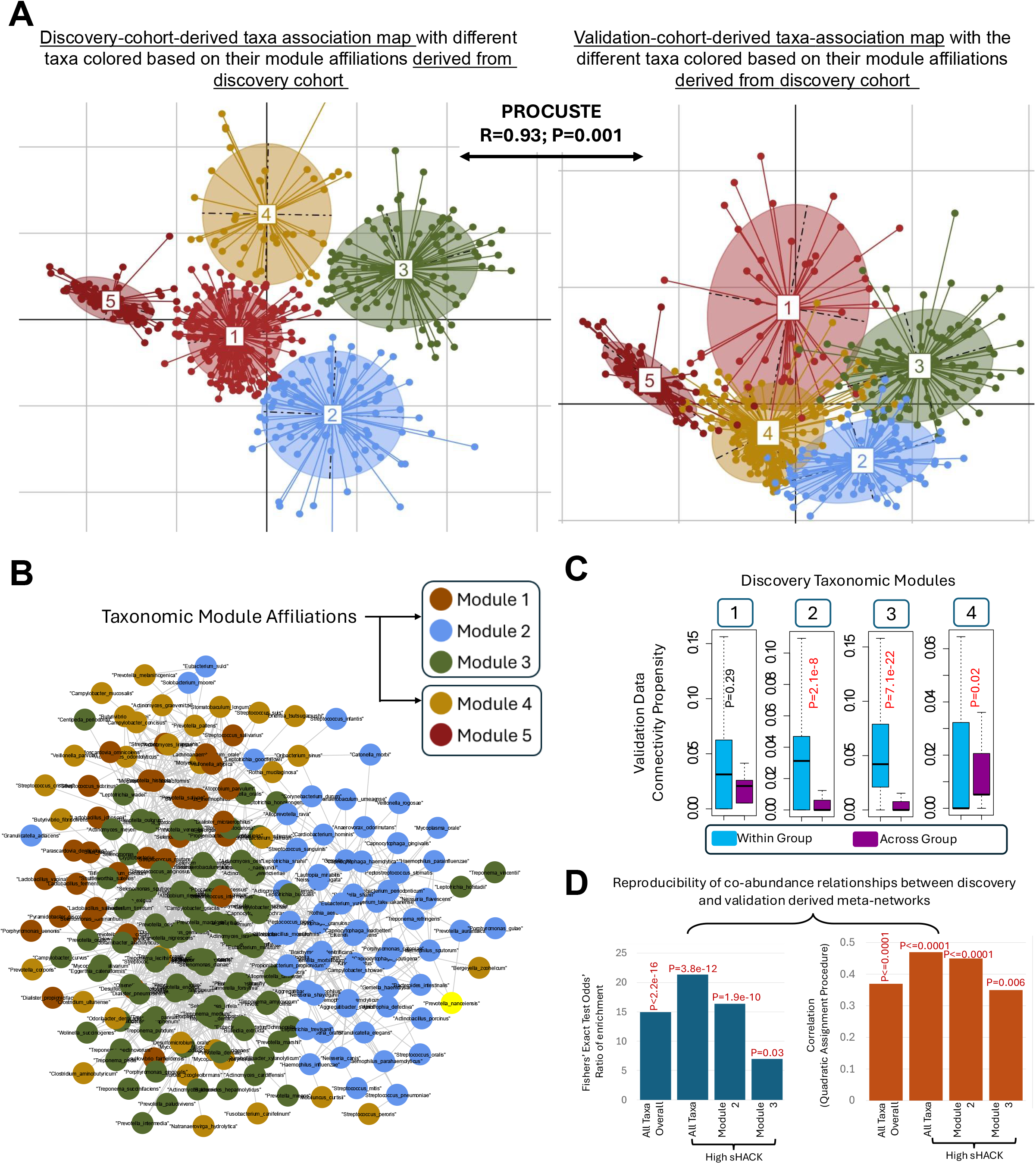
Reproducibility of salivary taxonomic modules in the validation cohort. **A.** Procrustes comparison of salivary taxon association maps (taxonomic modules) generated from discovery and validation cohorts. Five modules identified in the discovery cohort, showing reproducible module-level organization in the validation dataset (Procrustes R = 0.93, P = 0.001). **B.** Co-abundance meta-network reconstructed from validation-cohort salivary microbiomes, with nodes coloured by discovery-defined taxonomic modules showing the similar cohesive pattern in the unseen validation cohort as in the discovery network - Figure 3G. The network shows reproducible co-localization of module-specific taxa, particularly modules 1, 2, and 3. **C.** Within- and across-module connectivity propensity in the validation network, showing stronger within-module co-abundance for major modules. **D.** Reproducibility of co-abundance relationships between discovery and validation meta-networks, evaluated using edge-overlap enrichment and network-topology similarity analyses using Quadratic Assignment Procedure. Significant overlap and similarity indicate that the salivary ecological modules, including the high-sHACK-enriched module 2, are reproducible in unseen validation cohorts.

To test whether these modules also formed co-abundant sub-communities in validation cohorts, we built an analogous meta-network. The resulting network of 1,128 edges again showed modules 1, 2, and 3 (modules defined based on discovery-cohort investigation) colocalizing (**Table S15**, **Figure 4B**), with significantly higher within- than across-module co-abundance-connections (**Figure 4C**). We assessed reproducibility between discovery- and validation-cohort networks using Fisher’s exact test and the quadratic assignment procedure (QAP; correlation between adjacency matrices against a permuted null) (**Text S6**); both showed significant edge enrichment and topological similarity (**Figure 4D**). This was even more pronounced among sHACK taxa, particularly the sHACK sub-community (module 2), indicating this ecological unit remains intact regardless of dataset.

### The high 28 taxa sHACK sub-community (module-2) is also significantly associated with longitudinal stability and consistently distinguishes health from disease in the unseen validation cohort

Given the reproducible organization between discovery and validation cohorts, we tested whether the five modules also showed reproducible Core-, Stability-, and Health-Association, using non-diseased controls (2,885 microbiomes, 19 cohorts), longitudinal cohorts (1,525 microbiomes, 6 cohorts), and matched case-control cohorts (2,782 microbiomes, 15 cohorts; 1,518 controls, 1,217 diseased, 16 disease categories, plus 47 remission/post-treatment samples) (**Table S1F**). Validation patterns mirrored discovery findings (**Figure 5A-C**), where modules 2 and 3 were enriched for core- and stability-associated taxa, while health-associated taxa were enriched specifically in module 2, reaffirming it as the predominant reservoir of sHACK.

**Figure 5.**
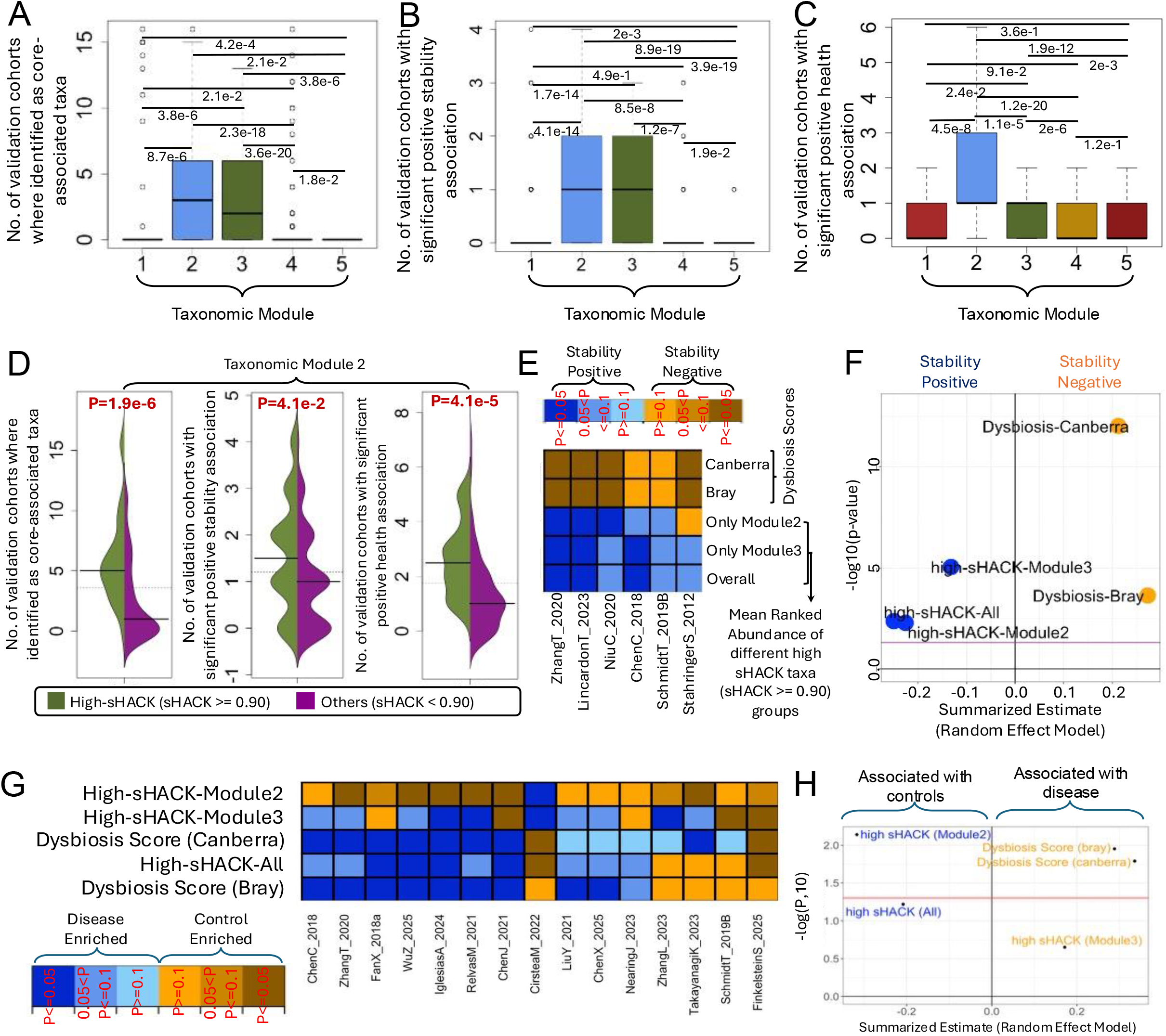
The 28-taxa high-sHACK module-2 panel reproducibly captures health, stability, and ecological associations across ‘unseen’ validation cohorts. **A-C.** Boxplots comparing, across the five taxonomic modules defined in the discovery cohorts, how often member taxa were independently identified in the validation cohorts as (**A**) core-associated, **(B)** stability-associated, and (**C**) health-associated. For each taxon, the y-axis gives the number of validation cohorts in which it was identified as associated with the corresponding property; boxplots summarize these per-taxon counts within each module. **D.** The same comparisons restricted to module 2, split into its high-sHACK (sHACK ≥ 0.90; the 28-taxa panel) and low-sHACK (sHACK < 0.90) subsets. Module 2 shows the strongest associations across all three properties overall, and within module 2 this signal is driven specifically by the 28-member high-sHACK subset. **E.** Cohort-wise associations between longitudinal microbiome stability and five indices, the mean rank-abundance of three high-sHACK taxon groups, plus two dysbiosis measures (Bray–Curtis and Canberra distances) across validation cohorts with longitudinal data. The three high-sHACK groups are: *Overall* (all taxa with sHACK ≥ 0.90), *Only module 2* (the 28-member module 2 high-sHACK subset), and *Only module 3* (module 3 taxa with sHACK ≥ 0.90). **F.** Random-effects meta-analysis summarizing the association of each index in (E) with longitudinal microbiome stability across validation cohorts. **G.** Cohort-wise comparison of the same five indices between control and diseased microbiomes across matched case–control validation cohorts. For the high-sHACK taxon-group indices, higher values in controls reflect enrichment of health-associated high-sHACK taxa (brown shades); for the Bray–Curtis and Canberra dysbiosis indices, disease association instead appears as higher values in diseased microbiomes (blue shades). **H.** Random-effects meta-analysis summarizing control–disease differences for the five indices across matched case–control validation cohorts, comparing the discriminatory ability of the three high-sHACK rank-abundance indices against the two dysbiosis indices. The 28-member module 2 high-sHACK taxon group shows the strongest and most consistent health-discriminatory performance of the five.

We assessed diagnostic potential of high-sHACK taxa (sHACK ≥0.70) in module 2, defining three panels - high-sHACK-All, 28 module-2 taxa, and 10 module-3 taxa-using mean rank-abundance as a diagnostic index, benchmarked against Bray-Curtis and Canberra dysbiosis indices. Across six longitudinal cohorts, dysbiosis indices correlated with reduced stability (significant in 4/6), while sHACK-derived indices showed the expected inverse pattern (significant in 3/6 each, one exception observed for module-2) (**Figure 5D-E**). Meta-analysis confirmed comparable performance overall (**Figure 5F**).

Testing health association across 15 matched cohorts, high-sHACK-module-2 was most consistent, scoring higher in controls in 14/15 cohorts (**Figure 5G**); Dysbiosis-score (with Canberra distances) performed next best (controls lower in 13/15, two reversed). Meta-analysis confirmed module-2 had the largest effect size distinguishing controls from diseased (**Figure 5H**). Thus, the 28-taxon module-2 panel outperformed broader panels and conventional dysbiosis indices for tracking stability and health.

### Health-associated core taxa show subsite-specific but recurrent module-level organization across three non-salivary oral habitats

We next asked whether similar ecological modules of health-associated, ecologically influential microbes could be identified in other oral subsites. As longitudinal data were unavailable, we integrated CS and HS scores into subsite-specific Health-Associated Core (HAC) scores (**Text S2**, **S4**), and identified ecological modules as for saliva-clustering taxa by community-wide association patterns and building co-abundance networks (**Text S5**). Results are summarised below (with details in **Texts S7-S10**).

In the supragingival subsite, 301 consensus taxa were analysed across 10,381 microbiomes from 24 cohorts (9,168 non-diseased, 1,213 diseased; **Table S16**; **Text S7**), ranked by CS, HS, with the two scores integrated into a HAC score (**Figures S16-S17**; **Table S17**), yielding 26 high-HAC taxa (HAC ≥0.90; **Figure S18A**). Module identification revealed three robust sub-communities with strong intra-community connections (**Figure 6A-B**; **Tables S18-S19**; **Figure S18B**). Module-2 showed the highest HAC values, containing 21 of the 26 high-HAC taxa, all with HS and CS ≥0.70 (odds ratio = 21.15, p = 1.4e-11; **Figure S18C**, **Figure 6C**), spanning 13 genera including *Cardiobacterium*, *Actinomyces*, *Capnocytophaga*, *Prevotella*, *Eikenella*, *Neisseria*, *Lautropia*, *Rothia*, *Corynebacterium*, *Gemella*, *Streptococcus*, *Campylobacter*, *Porphyromonas*, and *Aggregatibacter*. 12 of these 21 (∼59%) overlapped with the salivary module-2 (**Figure 3B**).

**Figure 6.**
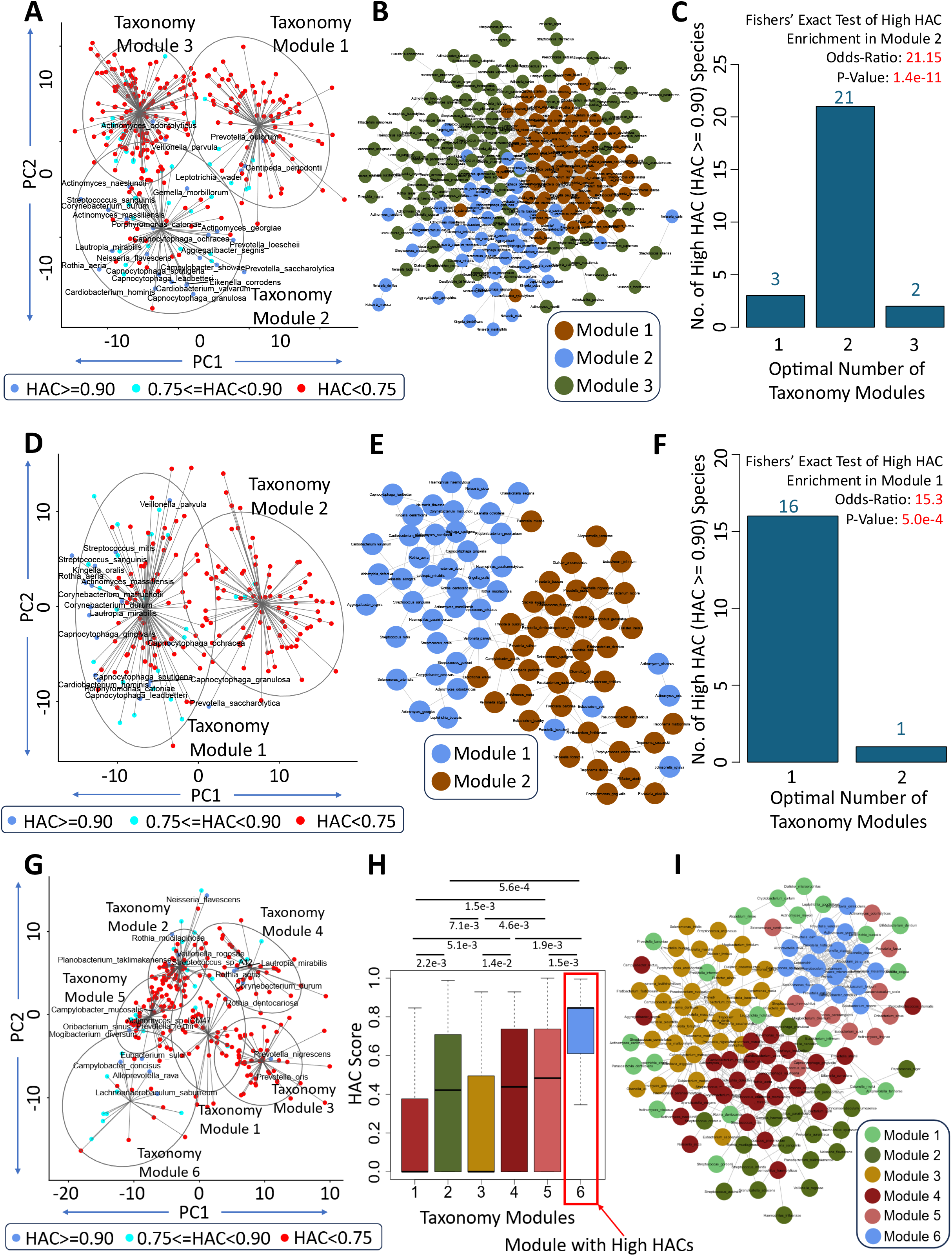
Identification of high-HAC sub-communities (or ecological modules) across non-salivary oral subsites. Module identifications for the supragingival, subgingival and tongue-tonsil subsites are described in A-C, D-F and G-I, respectively. **Community-wide association maps of consensus taxa** for each of the subsites are pictorially shown in **A.** supragingival, **D.** subgingival and **G.** tongue–tonsil subsites. Each community-wide association map is generated from PCA of taxon–taxon association profiles and clustered into taxonomic modules while colour is based on the HAC value. These panels highlight the spread of high HAC across the modules. **Co-abundance meta-networks of consensus taxa** are provided for the different subsites in **B.** Supragingival**, E.** Subgingival **I.** Tongue–tonsil subsites. Networks were generated by random-effects meta-analysis of cohort-wise robust linear regression estimates across non-diseased samples. Taxon pairs with a positive summarized estimate, Benjamini–Hochberg-adjusted Q ≤ 0.05, and directional consistency ≥ 0.70 were connected by edges. Networks were visualized in Cytoscape using the Compound Spring Embedder (CoSE) layout, with nodes coloured according to module identity. **Distribution of taxa with HAC ≥ 0.90 across taxonomic modules** are provided in **C.** for supragingival and **E.** for subgingival subsites. In the supragingival subsite, high-HAC taxa were significantly enriched in module 2 (n = 21; odds ratio = 21.15; Fisher’s exact test, p = 1.4e-11). In the subgingival subsite, high-HAC taxa were significantly enriched in module 1 (n = 16; odds ratio = 15.3; Fisher’s exact test, p = 5.0e-4). **(H) Distribution of all taxa HAC score across the 6 modules in the tongue-tonsil subsite**. Module 6 shows the highest HAC distribution as compared to any other module. The details pertaining to the identification of the high HAC taxa and the ecological modules for each subsite are provided as: Supragingival: **Text S7**, **Figures S17-S18**, **Tables S16-19**; Subgingival: **Text S8**, **Figures S20-21**, **Tables S20-S23**; Tongue-Tonsil: **Text S9**, **Table S24-S27**, **Figure S23-S24**.

A comparable pattern emerged in the subgingival subsite (196 taxa; 1,974 microbiomes, 11 cohorts; 1,418 non-diseased, 556 diseased; **Text S8**; **Table S20**), yielding 17 taxa with HAC ≥0.90 (**Figures S19-S21A**; **Table S21**). Two sub-communities were identified (**Figure 6D**; **Figure S21B**; **Table S22**), further validated via taxa-to-taxa co-abundance meta-network (**Figure 6E**; **Table S23**). Module-1 showed the highest HAC scores (**Figure S21C**), containing 16 of 17 high-HAC taxa (odds ratio = 15.3, p = 5.0e-4; **Figure S21A**, **Figure 6F**), spanning 11 genera including *Capnocytophaga*, *Corynebacterium*, *Streptococcus*, *Veillonella*, *Actinomyces*, *Rothia*, *Cardiobacterium*, *Lautropia*, *Kingella*, and *Porphyromonas*; 11 (∼69%) overlapped with supragingival high-HAC module-2, while only 6 (∼38%) overlapped with salivary high-sHACK module-2.

The tongue-tonsil subsite showed a more distributed organization (266 consensus taxa; 1,693 microbiomes, 18 cohorts; 1,570 non-diseased, 123 diseased; **Text S9**; **Table S24-S27**; **Figures S22-S24**, **6G-I**). Only five taxa, *Mogibacterium diversum*, *Eubacterium sulci*, *Alloprevotella rava*, *Neisseria flavescens*, and *Prevotella jejuni*, showed concordantly high HS and CS (**Figure S23**; **Table S25A**). The 20 top-HAC taxa included these members (**Figure S24A**; **Table S25B**). Ecological module identification revealed six modules (**Table S26**; **Figure S24B**, **6G**), validated via co-abundance meta-network (**Table S27**; **Figure 6I**). Module 6 showed the highest HAC distribution overall (**Figure 6H**), yet only three of the top-HAC taxa colocalized to this module (odds ratio = 1.76, p = 0.26; **Figure S24C**); its elevated HAC values instead primarily reflected a smaller proportion of low-HAC taxa (HAC score ≤ 0.50) within the module.

The buccal-palate-other surface subsite was analysed conservatively due to limited case-control data (749 microbiomes, 11 cohorts; 521 non-diseased, 221 diseased; 166 consensus taxa; **Text S10**; **Table S28**). Core-associated taxa included *Rothia mucilaginosa*, *Fusobacterium nucleatum*, *Actinomyces odontolyticus*, *Haemophilus parainfluenzae*, and *Streptococcus mitis* (**Figure S25**; **Table S29A**); HS and HAC were not computed due to insufficient matched cohorts. Direct disease-control comparison (combining all cohorts) identified *Prevotella shahii* as control-associated, while *Actinomyces naeslundii*, *Capnocytophaga gingivalis*, *C. sputigena*, *C. leadbetteri* and *Streptococcus sanguinis* were disease-associated (**Table S29B**), the opposite of their high-HAC, health-associated status in the gingival (sub/supragingival) and salivary subsites.

Together, these analyses identified health-associated core taxa across oral habitats, with supragingival and subgingival sites showing the clearest ecological modules. Notably, high-HAC taxa overlapped more between gingival sites than with saliva, and several taxa that were high-HAC in gingival and salivary subsites showed reversed health associations in the buccal-palate-other-surface subsite. We next examined these inter-site relationships.

### High HAC taxa show both shared and subsite-specific patterns across oral habitats including Salivary subsite

We examined whether ecologically important, health-associated microbial hallmarks were shared across oral subsites or remained niche-specific, comparing high-HAC taxa lists (HAC≥0.90) across four subsites (buccal-palate-subsite excluded for low sample size). For consistency, salivary taxa were also ranked by HAC (CS and HS only) rather than sHACK (**Figure 3A**); the two measures correlated strongly (R=0.78, P=7.5e-15; **Figure 7A**).

**Figure 7.**
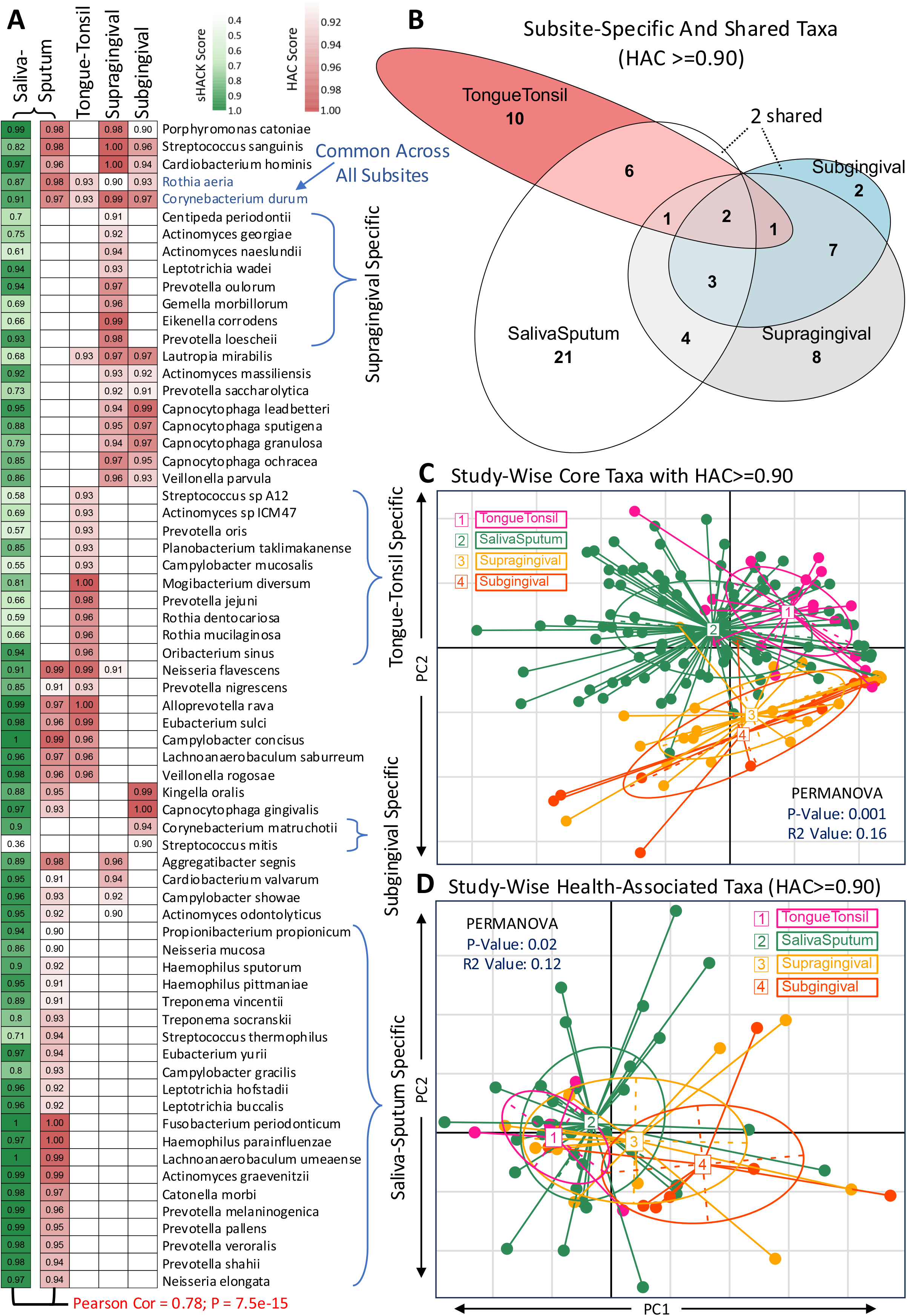
Shared and subsite-specific patterns of high-HAC taxa across oral subsites reflect location- and subsite-specific micro-environments. **A.** Heatmap of HAC values for taxa ranked within the top 10% of HAC scores in at least one oral subsite. Rows represent taxa and columns represent oral subsites, highlighting taxa with broadly shared versus subsite-specific high-HAC patterns. For the salivary subsite, HAC scores (rather than sHACK) were used to allow direct comparison with the non-salivary subsites; the salivary sHACK score is shown separately as a bar on the left. **B.** Venn diagram showing the overlap of top 10% high-HAC taxa across oral subsites, summarizing taxa shared across multiple oral habitats as well as taxa unique to individual subsites. **C.** PCoA of study-cohort-wise core-association patterns, based on the top 10% high-HAC taxa for each subsite. Points represent study-cohorts, coloured by oral subsite. PERMANOVA showed significant subsite-level separation of core-association patterns (R² = 0.16, P = 0.001), indicating that the recurrence of core-associated taxa differs across oral habitats. **D.** PCoA of study-cohort-wise health-association patterns, based on the same top 10% high-HAC taxa. PERMANOVA showed significant subsite-level variation in health-association patterns, though weaker than for core-association (R² = 0.12, P = 0.02). Among the four subsites, the two gingival subsites (subgingival and supragingival) were most similar to each other, and both diverged most strongly from the tongue-tonsil subsite. The salivary subsite overlapped with both the gingival and tongue-tonsil subsites, more closely with tongue-tonsil than either gingival subsite showed with tongue-tonsil, but less closely with the gingival subsites than the two gingival subsites showed with each other.

Only two taxa, *Rothia aeria* and *Corynebacterium durum*, were high-HAC across all four subsites. Site-specific HAC taxa were most numerous in saliva (21), followed by tongue-tonsil (10), supragingival (8), and subgingival (2). Ten taxa were shared between the two gingival subsites, three of these also present in saliva; nine taxa were shared between saliva and at least one gingival subsite, seven between saliva and tongue-tonsil, and only two between tongue-tonsil and either gingival subsite.

Venn-diagram analysis confirmed strong gingival-gingival overlap (subgingival taxa largely nested within supragingival), minimal overlap with the ecologically distinct tongue-tonsil site, and saliva sharing taxa with both while retaining the largest subsite-unique set (**Figure 7B**).

Ordination showed significant study-specific separation based on subsite in core- and health-association patterns (PERMANOVA: Core-Association: R²=0.16, P=0.001; Health-Association: R²=0.12, P=0.02; **Figure 7C-D**), reflecting niche-specific signatures alongside some cross-habitat conservation, tracking microenvironmental similarity between sites.

### Genome-derived functional profiles predict subsite-specific sHACK/HAC scores derived using Microbiome Profile

We next asked whether specific genome-encoded functions underlie the high-HAC and high-sHACK taxa across oral subsites, that is, whether these ecologically important, health-associated microbes have specific functional signatures at each site.

We built genome-derived functional profiles from 71,238 high-quality reference and MAG genomes, annotated via eggNOG-mapper^29^ (with Prodigal-based gene prediction) across six categories: CAZy^30^, COG^31^, BiGG^32^, KEGG^33^, EC^34^, and Pfam^35^ (**Text S11**). Each species was assigned a single profile by averaging annotations across its genomes. Matching these to subsite-specific consensus taxa yielded functionally annotated sets for saliva (366 taxa), supragingival (220), subgingival (140), and tongue-tonsil (190).

For each subsite, a random forest model was trained to predict each taxon’s health-association score (sHACK for saliva; HAC elsewhere) from its functional profile, progressively trimming to the smallest feature set that preserved performance (**Figure 8A**; **Text S11-S12**). The best model per subsite defined that site’s HAC/sHACK-associated functions, and directionality was determined by correlating feature abundances with HAC/sHACK scores.

**Figure 8.**
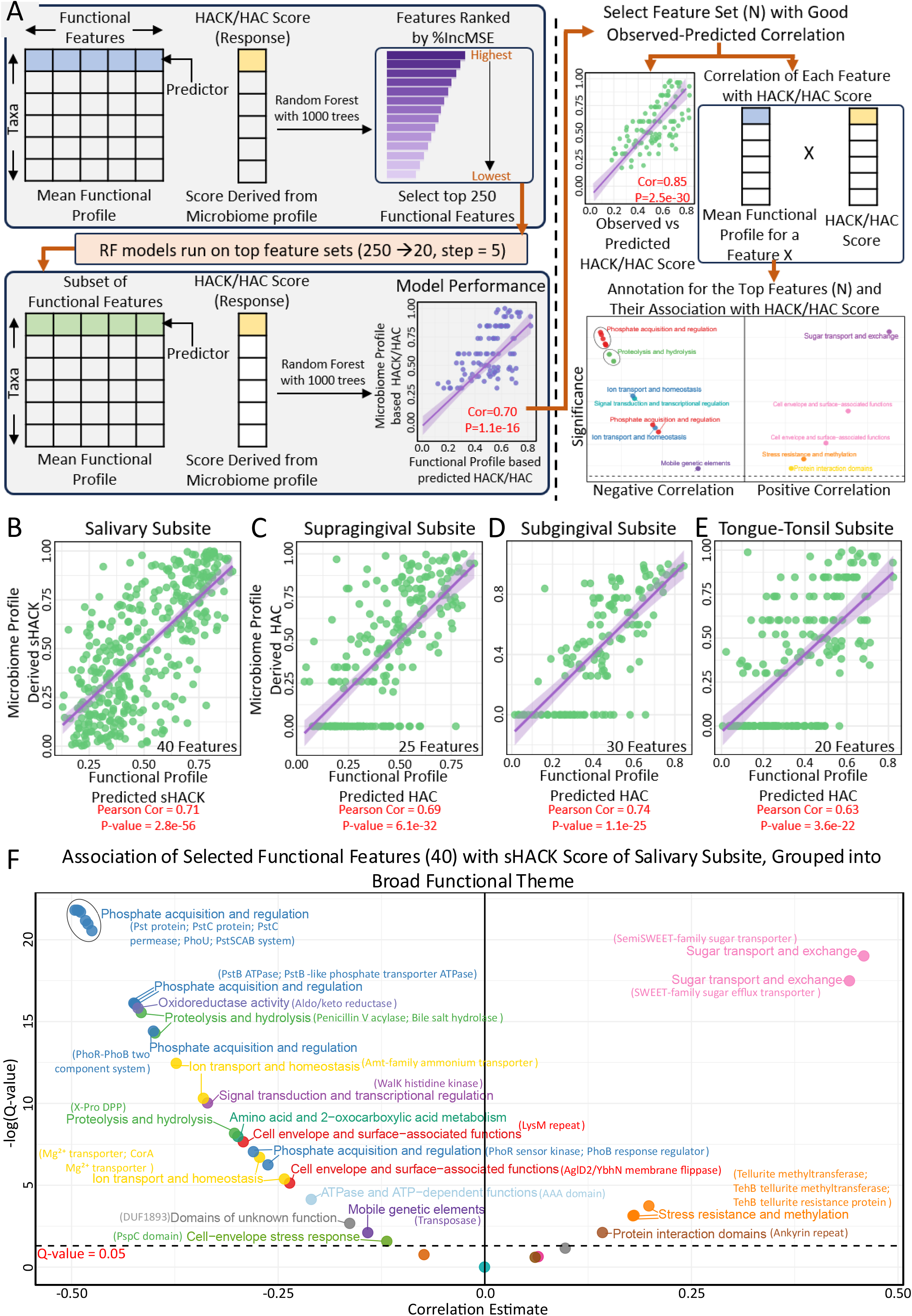
Functional-profile-based prediction and interpretation of oral HACK/HAC scores. **A.** Schematic overview of the functional-profile-based prediction framework. For each subsite, species-level-taxa-specific mean functional profiles derived from our 71,234 genome collection (**Text S11**) were used as predictor variables in a Random Forest model to predict the corresponding taxa-specific sHACK/HAC scores independently obtained using the microbiome abundance profiles for that subsite. An initial Random Forest model was first built taking into consideration all functional features and the features were then ranked in decreasing order by their corresponding feature importance values (defined by the % increase in mean-squared-error values or %IncMSE values). Subsequently, we generated iterative models starting with the top 250 features using reduced feature sets from 250 to 20, decreasing by 5 features at each step. For each feature subset, predicted sHACK/HAC scores were compared with observed scores to evaluate model performance. The final feature set was selected based as that minimum set of features for which the high predicted-versus-observed correlation was either the highest or retained as compared to the top 250 features. After the identification of the top feature subsets for each subsite, the species-taxa-level mean abundances of the corresponding feature sets were then correlated with the corresponding taxon-specific HAC/sHACK scores to identify the functions that are positively or negatively correlated with the corresponding HAC/sHACK scores. **B-E.** Predicted-versus-observed score correlations for the selected functional-feature models across oral subsites. Each point represents a taxon, with the x-axis showing the functional-profile-based predicted score and the y-axis showing the observed microbiome-profile-derived sHACK/HAC score. Pearson correlation coefficients and corresponding p values indicate the predictive performance of the selected functional feature sets in each subsite (Salivary, Supragingival, Subgingival, Tongue-Tonsil subsite respectively). **F. Volcano plot showing correlations between selected 40 functional features and observed sHACK scores in Salivary subsite.** The x-axis represents the correlation coefficient, and the y-axis represents -log10(Q-value). Each point represents a selected functional feature, coloured according to broad functional theme. The dashed horizontal line indicates Benjamini-Hochberg adjusted Q = 0.05. Positively correlated features are associated with higher sHACK scores, whereas negatively correlated features are associated with lower sHACK scores (**Table S30**). Similar results for the other subsites are provided in **Text S12**, **Figures S27-S29**, **Tables S31-S33**.

Taxon-level scores were strongly predictable from functional profiles alone, with significant out-of-bag correlations (Saliva: 40 features, R=0.71, P=2.8e-56; Supragingival: 25 features, R=0.69, P=6.1e-32; Subgingival: 30 features, R=0.74, P=1.1e-25; Tongue-tonsil: 20 features, R=0.63, P=3.6e-22; **Figures 8B-E**, **S26-S29**).

In saliva, high-sHACK taxa were positively linked to sugar transport/exchange (sugar efflux and SemiSWEET transporters), suggesting greater predicted carbohydrate-transport and metabolic-exchange capacity among high-sHACK taxa^36–38^, along with stress-resistance features (tellurite methyltransferase, ankyrin repeats), indicating stress-tolerance and protein-interaction potential^39–43^. Low-sHACK taxa showed enrichment in phosphate acquisition and regulation (PhoB, PhoR, PhoU, ABC-type phosphate transport components)^44–48^, along with ion transport, proteolysis/hydrolysis^49–52^, cell-envelope stress response, signal transduction, and mobile genetic elements, pointing to a contrast between metabolic-exchange capacity in high-sHACK taxa and nutrient-scavenging/stress-adaptation in low-sHACK taxa (**Figure 8F**; **Table S30**; **Text S12**).

Similar predictability held across non-salivary subsites, but functional themes diverged by habitat. Supragingival HAC scores also correlated positively with sugar-transport features, inversely with ammonium-nitrogen transport and translation-related functions (**Figure S27**; **Table S31**). Subgingival HAC scores instead associated positively with amino acid/arginine-polyamine metabolism, oxidative-stress response, and cell-surface interaction domains, with no significant negative associations (**Figure S28**; **Table S32**). Tongue-tonsil HAC scores were positively linked to regulatory, protein-interaction, and ATP-dependent features, and negatively to ribosomal, DNA-replication, and membrane-translocation functions (**Figure S29**; **Table S33**).

Together, these results show that HAC/sHACK rankings capture real, measurable differences in genome-derived functional potential, with saliva and supragingival sites sharing sugar-transport signatures, while subgingival and tongue-tonsil sites carry distinct functional signatures of their own (**Text S12**).

## Discussion

This large multi-cohort study, spanning 37,739 oral microbiomes from 142 cohorts across 41 countries, establishes a subsite-resolved framework for ranking oral taxa by health-association, ecological influence, and prevalence consistency, further incorporating longitudinal stability where available, to derive Health-Associated-Core (HAC) scores across four major subsites and a salivary-specific HAC-Keystone (sHACK) score. Beyond taxon ranking, the framework identifies ecological modules, reproducible sub-communities within each subsite, by grouping taxa based on shared association patterns with the rest of the community and direct co-abundance relationships across cohorts. Convergence between these two independent methods confirms these modules are genuine ecological units rather than analytical artifacts, and reveals that health-associated keystone taxa concentrate within discrete, biologically cohesive guilds rather than distributing independently.

These findings carry translational potential: low-dimensional taxa panels could serve as measures of subsite-specific oral health and stability. Our salivary module-2 (28 taxa) exemplifies this, showing strong associations with both microbiome stability and health status, outperforming conventional dysbiosis indices in the latter. This panel is also a strong candidate consortium for defined-community probiotic or synbiotic development.

This subsite-resolved analysis proved essential. Only two taxa were conserved as high-health-associated across all four subsites, with overlap otherwise tracking anatomical proximity and micro-environmental similarity. Gingival subsites shared strong overlap distinct from tongue-tonsil, while saliva overlapped with all three. Notably, some taxa (e.g., *Capnocytophaga* members) were health-associated in saliva and gingival sites but disease-associated in buccal microbiomes, underscoring that health-association is niche-dependent rather than universal.

Genome-derived functional modelling added a further layer: compact functional feature sets predicted sHACK/HAC scores across subsites, showing that prioritised taxa differ functionally as well as taxonomically and ecologically. Saliva and supragingival taxa shared sugar-transport/exchange associations, while subgingival and tongue-tonsil taxa showed distinct signatures, likely reflecting micro-environmental differences. As these profiles are genome-derived, they capture functional potential rather than confirmed in vivo activity, and given incomplete functional annotations, the mechanistic basis of these associations warrants further validation.

This study has additional limitations: reliance on public datasets introduces variability in geography, protocols, platforms, and metadata completeness; longitudinal stability was assessable only for saliva; co-abundance modules do not establish direct ecological interactions; and functional predictions require validation via metatranscriptomics, metaproteomics, metabolomics, and experimental models, as many findings remain associative and inferred.

Despite these limitations, this work provides a quantitative framework for defining health-associated core taxa and ecological modules across oral micro-environments. These findings offer a foundation for future studies to develop habitat-resolved biomarker panels, functionally probe subsite-specific health-associated pathways, and design microbial consortia for mechanistic testing, microbiome-guided oral health monitoring, and oral-health-associated biotherapeutics.

### Code and Data Availability

This study analyses existing, publicly available data from the European Nucleotide Archive (ENA) and curated Metagenomic Data (see **Methods**); cohort details are listed in **Table S1**. Code used and results generated during the analyses is publicly available at https://github.com/omprakash414/MetaOral_Atlas, with large files hosted on Zenodo (linked from the repository).

## Acknowledgments

T.S.G. and O.S. acknowledge the Department of Biotechnology, Ministry of Science and Technology, Government of India, for the Ramalingaswami Re-entry Fellowship (BT/HRD/35/02/2006). A.A. acknowledges the Department of Science and Technology (DST-INSPIRE) for the INSPIRE fellowship. S.G. acknowledges IIIT-Delhi for the Institute-Fellowship.

## Author Contributions

Conceptualization: T.S.G. Data collection and curation: T.S.G., O.S., M.V., A.P., E.C., A.A.; Resources (genome and functional profile generation): S.G.; Methodology: O.S., T.S.G; Formal analysis: O.S., A.A., T.S.G.; Validation: O.S., A.A., T.S.G.; Investigation: T.S.G., O.S. Writing – original draft: O.S., T.S.G; Writing - review & editing: T.S.G.; Supervision: T.S.G.; Funding acquisition: T.S.G.

## Methods

### Dataset collation, taxonomic profiling, and metadata harmonization

Publicly available human oral microbiome datasets were collated from the European Nucleotide Archive (ENA)^25^, curated Metagenomic Data (cMD)^53^ and PubMed. ENA was used as the primary source for raw sequencing data, while PubMed was used to identify additional relevant studies and verify associated publications. Retrieved studies were screened for human oral microbiome relevance, availability of raw sequencing data, and sufficient metadata describing sample source, disease or control status, country, sequencing strategy, and available demographic or exposure variables.

Raw sequence data were downloaded and processed using a sequencing-technology-specific taxonomic profiling workflow. Whole-genome shotgun metagenomic samples were profiled using MetaPhlAn3^26^, whereas 16S rRNA amplicon datasets were profiled using SPINGO^27^. Studies containing multiple sequencing strategies or multiple oral sampling sites were partitioned into separate analytical study-cohorts.

Metadata were manually harmonised across studies using standardised terms for disease status, sequencing strategy, country, demographic variables, exposure variables, and oral sampling site. The metadata for few studies were collected from their respective authors^54–59^. Author-reported sampling terms were mapped to five standardised oral subsite categories: saliva-sputum-oral wash (salivary), supragingival subsite, subgingival subsite, tongue-tonsil-related sites, and buccal-palate-other surface sites. Detailed study-level information, sample-level metadata, and subsite mapping are provided in **Tables S1-S3**.

### Identification of subsite-resolved consensus oral-associated taxa

Consensus taxa were identified independently within each oral subsite using a two-threshold optimization framework adapted from our previous gut microbiome meta-analysis approach^14^. Detailed threshold selection and subsite-specific retained taxon sets are provided in **Text S1, Figures S1-S5, and Table S4**.

### Computation of taxon-level association scores

Taxon-level association scores were computed to quantify three complementary properties: consistency of association with health status, recurrence as core-associated members (highly prevalent and ecologically influential members of the microbiome, and association with longitudinal microbiome stability (explained in detail in **Text S2**). The scoring frameworks were adapted from our previous gut microbiome meta-analysis study^14^, and applied independently within the different oral subsites where sufficient data were available. Health-Association Scores (HS) were computed from matched case-control cohorts, Core-Association Scores (CS) from non-diseased control cohorts using prevalence and community-structure association criteria, and Stability-Association Scores (SS) from longitudinal salivary cohorts using follow-up microbiome distance. For the salivary subsite, scores were additionally recomputed within sequencing- and exposure-stratified cohort subsets to assess robustness across data-generation modalities. Detailed score computation, thresholds, formulas, and stratified analyses are provided in **Text S2**, and **Figures S6**, **S9**, **S11**.

### Integrated sHACK and HAC indices

Integrated taxon-level indices were computed to prioritize taxa with balanced associations across scoring dimensions. The salivary sHACK index integrated CS, HS, and SS, whereas HAC integrated CS and HS for subsites without longitudinal stability data, using a Gini-based concordance framework. Detailed methodology is provided in **Text S4**, **Figure 3A**.

### Taxonomic module detection in oral-subsites

To investigate whether high-sHACK taxa formed structured ecological sub-communities, pairwise taxon-taxon Association Scores were computed across non-diseased control cohorts for each subsite and used to generate a PCA-based association map. Taxa were then partitioned into modules using k-means clustering, with the optimal number of clusters selected using the Calinski–Harabasz index. Detailed methodology is provided in **Text S5** and **Figure S15**.

### Co-abundance network construction in discovery and validation cohorts

Co-abundance meta-networks were constructed by testing taxon-pair associations within cohorts using robust linear regression and combining cohort-level effects using random-effects meta-analysis. Edges were retained for taxon pairs showing positive summarised effects, FDR-corrected Q-value, and consistency ≥ 0.70. Additionally for salivary subsites discovery and validation-cohort networks were compared using Fisher’s exact test for edge overlap and the quadratic assignment procedure for network-level similarity. Detailed methodology is provided in **Text S5-S6**.

### Genome-derived functional modelling of oral sHACK/HAC scores

Genome-derived species-level functional profiles were used to model the relationship between functional potential and subsite-specific sHACK/HAC scores. For each oral subsite, species matched between the consensus taxon set and the functional-profile matrix were used for random forest regression, with functional groups as predictors and the corresponding sHACK or HAC score as the response. Functional features were ranked by %IncMSE, followed by iterative evaluation of reduced top-feature sets to identify compact predictive models. Selected features were further assessed for their individual associations with sHACK/HAC scores and summarised into broad functional themes; detailed methodology is provided in **Text S11** and **Figure 8A**.

## Supplementary Information

Supplementary Document: Contains Supplementary Texts S1-S12 and Supplementary Figures with Legends S1-S29

Supplementary Tables are provided in multiple sheets - Tables S1-S33

## References

1. Huttenhower, C. et al. Structure, function and diversity of the healthy human microbiome. Nature 486, 207–214 (2012).

2. Baker, J. L., Mark Welch, J. L., Kauffman, K. M., McLean, J. S. & He, X. The oral microbiome: diversity, biogeography and human health. Nat. Rev. Microbiol. 22, 89–104 (2024).

3. Caballero-Flores, G., Pickard, J. M. & Núñez, G. Microbiota-mediated colonization resistance: mechanisms and regulation. Nat. Rev. Microbiol. 21, 347–360 (2023).

4. Lamont, R. J., Koo, H. & Hajishengallis, G. The oral microbiota: dynamic communities and host interactions. Nat. Rev. Microbiol. 16, 745–759 (2018).

5. Cai, L. et al. Integrative analysis reveals associations between oral microbiota dysbiosis and host genetic and epigenetic aberrations in oral cavity squamous cell carcinoma. Npj Biofilms Microbiomes 10, 39 (2024).

6. Tonelli, A., Lumngwena, E. N. & Ntusi, N. A. B. The oral microbiome in the pathophysiology of cardiovascular disease. Nat. Rev. Cardiol. 20, 386–403 (2023).

7. Read, E., Curtis, M. A. & Neves, J. F. The role of oral bacteria in inflammatory bowel disease. Nat. Rev. Gastroenterol. Hepatol. 18, 731–742 (2021).

8. Burcham, Z. M. et al. Patterns of Oral Microbiota Diversity in Adults and Children: A Crowdsourced Population Study. Sci. Rep. 10, 2133 (2020).

9. Manghi, P. et al. Large-scale metagenomic analysis of oral microbiomes reveals markers for autism spectrum disorders. Nat. Commun. 15, 9743 (2024).

10. Zhao, H. et al. Variations in oral microbiota associated with oral cancer. Sci. Rep. 7, 11773 (2017).

11. Belstrøm, D. et al. Periodontitis associates with species-specific gene expression of the oral microbiota. Npj Biofilms Microbiomes 7, 76 (2021).

12. Wolf, A. et al. The salivary microbiome as an indicator of carcinogenesis in patients with oropharyngeal squamous cell carcinoma: A pilot study. Sci. Rep. 7, 5867 (2017).

13. Nearing, J. T., DeClercq, V. & Langille, M. G. I. Investigating the oral microbiome in retrospective and prospective cases of prostate, colon, and breast cancer. Npj Biofilms Microbiomes 9, 23 (2023).

14. Goel, A. et al. Toward a health-associated core keystone index for the human gut microbiome. Cell Rep. 44, (2025).

15. Mark Welch, J. L., Rossetti, B. J., Rieken, C. W., Dewhirst, F. E. & Borisy, G. G. Biogeography of a human oral microbiome at the micron scale. Proc. Natl. Acad. Sci. 113, E791–E800 (2016).

16. Proctor, D. M. & Relman, D. A. The Landscape Ecology and Microbiota of the Human Nose, Mouth, and Throat. Cell Host Microbe 21, 421–432 (2017).

17. Segata, N. et al. Composition of the adult digestive tract bacterial microbiome based on seven mouth surfaces, tonsils, throat and stool samples. Genome Biol. 13, R42 (2012).

18. Hall, M. W. et al. Inter-personal diversity and temporal dynamics of dental, tongue, and salivary microbiota in the healthy oral cavity. Npj Biofilms Microbiomes 3, 2 (2017).

19. Ruan, X., Luo, J., Zhang, P. & Howell, K. The salivary microbiome shows a high prevalence of core bacterial members yet variability across human populations. Npj Biofilms Microbiomes 8, 85 (2022).

20. Xian, P. et al. The Oral Microbiome Bank of China. Int. J. Oral Sci. 10, 16 (2018).

21. Lin, Y. et al. Omics for deciphering oral microecology. Int. J. Oral Sci. 16, 2 (2024).

22. Miller, D. P., Fitzsimonds, Z. R. & Lamont, R. J. Metabolic Signaling and Spatial Interactions in the Oral Polymicrobial Community. J. Dent. Res. 98, 1308–1314 (2019).

23. Wu, C. M. et al. Mucin glycans drive oral microbial community composition and function. Npj Biofilms Microbiomes 9, 11 (2023).

24. O’Toole, P. W., Marchesi, J. R. & Hill, C. Next-generation probiotics: the spectrum from probiotics to live biotherapeutics. Nat. Microbiol. 2, 1–6 (2017).

25. Yuan, D. et al. The European Nucleotide Archive in 2025. Nucleic Acids Res. 54, D120– D127 (2026).

26. Beghini, F. et al. Integrating taxonomic, functional, and strain-level profiling of diverse microbial communities with bioBakery 3. eLife 10, e65088 (2021).

27. Allard, G., Ryan, F. J., Jeffery, I. B. & Claesson, M. J. SPINGO: a rapid species-classifier for microbial amplicon sequences. BMC Bioinformatics 16, 324 (2015).

28. Goswami, S. et al. Gut microbiome features associated with Bifidobacterium colonization predict personalized probiotic persistence patterns. Nat. Commun. 17, 5678 (2026).

29. Cantalapiedra, C. P., Hernández-Plaza, A., Letunic, I., Bork, P. & Huerta-Cepas, J. eggNOG-mapper v2: Functional Annotation, Orthology Assignments, and Domain Prediction at the Metagenomic Scale. Mol. Biol. Evol. 38, 5825–5829 (2021).

30. Drula, E. et al. The carbohydrate-active enzyme database: functions and literature. Nucleic Acids Res. 50, D571–D577 (2022).

31. Galperin, M. Y. et al. COG database update: focus on microbial diversity, model organisms, and widespread pathogens. Nucleic Acids Res. 49, D274–D281 (2021).

32. Norsigian, C. J. et al. BiGG Models 2020: multi-strain genome-scale models and expansion across the phylogenetic tree. Nucleic Acids Res. 48, D402–D406 (2020).

33. Kanehisa, M., Furumichi, M., Sato, Y., Kawashima, M. & Ishiguro-Watanabe, M. KEGG for taxonomy-based analysis of pathways and genomes. Nucleic Acids Res. 51, D587– D592 (2023).

34. Duvaud, S. et al. Expasy, the Swiss Bioinformatics Resource Portal, as designed by its users. Nucleic Acids Res. 49, W216–W227 (2021).

35. Mistry, J. et al. Pfam: The protein families database in 2021. Nucleic Acids Res. 49, D412– D419 (2021).

36. Xu, Y. et al. Structures of bacterial homologues of SWEET transporters in two distinct conformations. Nature 515, 448–452 (2014).

37. Esberg, A., Haworth, S., Hasslöf, P., Lif Holgerson, P. & Johansson, I. Oral Microbiota Profile Associates with Sugar Intake and Taste Preference Genes. Nutrients 12, 681 (2020).

38. Jia, B., Hao, L., Xuan, Y. H. & Jeon, C. O. New Insight Into the Diversity of SemiSWEET Sugar Transporters and the Homologs in Prokaryotes. Front. Genet. 9, (2018).

39. Al-Khodor, S., Price, C. T., Kalia, A. & Abu Kwaik, Y. Functional diversity of ankyrin repeats in microbial proteins. Trends Microbiol. 18, 132–139 (2010).

40. Peng, W. et al. The antibacterial effect of tellurite is achieved through intracellular acidification and magnesium disruption. mLife 4, 423–436 (2025).

41. Gaete-Argel, A. et al. Tellurite Promotes Stress Granules and Nuclear SG-Like Assembly in Response to Oxidative Stress and DNA Damage. Front. Cell Dev. Biol. 9, (2021).

42. Kang, Y. et al. Dental Plaque Microbial Resistomes of Periodontal Health and Disease and Their Changes after Scaling and Root Planing Therapy. mSphere 6, 10.1128/msphere.00162-21 (2021).

43. Sandoval, J. M., Levêque, P., Gallez, B., Vásquez, C. C. & Buc Calderon, P. Tellurite-induced oxidative stress leads to cell death of murine hepatocarcinoma cells. BioMetals 23, 623–632 (2010).

44. Baek, S. et al. PhoU interaction with the PhoR PAS domain is required for repression of the pho regulon and Salmonella virulence, but not for polyphosphate accumulation. J. Microbiol. 63, e2505013 (2025).

45. Martín, J. F. & Liras, P. Molecular Mechanisms of Phosphate Sensing, Transport and Signalling in Streptomyces and Related Actinobacteria. Int. J. Mol. Sci. 22, 1129 (2021).

46. Bruna, R. E., Kendra, C. G. & Pontes, M. H. An intracellular phosphorus-starvation signal activates the PhoB/PhoR two-component system in Salmonella enterica. mBio 15, e01642–24 (2024).

47. Lamarche, M. G., Wanner, B. L., Crépin, S. & Harel, J. The phosphate regulon and bacterial virulence: a regulatory network connecting phosphate homeostasis and pathogenesis. FEMS Microbiol. Rev. 32, 461–473 (2008).

48. Lubin, E. A., Henry, J. T., Fiebig, A., Crosson, S. & Laub, M. T. Identification of the PhoB Regulon and Role of PhoU in the Phosphate Starvation Response of Caulobacter crescentus. J. Bacteriol. 198, 187–200 (2015).

49. Agolino, G. et al. Bile salt hydrolase: The complexity behind its mechanism in relation to lowering-cholesterol lactobacilli probiotics. J. Funct. Foods 120, 106357 (2024).

50. Lucas, L. N. et al. Investigation of bile salt hydrolase activity in human gut bacteria reveals production of conjugated secondary bile acids. Nat. Commun. 17, 3077 (2026).

51. Guo, X. et al. Interactive Relationships between Intestinal Flora and Bile Acids. Int. J. Mol. Sci. 23, 8343 (2022).

52. Di Vincenzo, F. et al. Bile Acid-Related Regulation of Mucosal Inflammation and Intestinal Motility: From Pathogenesis to Therapeutic Application in IBD and Microscopic Colitis. Nutrients 14, 2664 (2022).

53. Pasolli, E. et al. Accessible, curated metagenomic data through ExperimentHub. Nat. Methods 14, 1023–1024 (2017).

54. Poole, A. C. et al. Human Salivary Amylase Gene Copy Number Impacts Oral and Gut Microbiomes. Cell Host Microbe 25, 553–564.e7 (2019).

55. Thomas, A. M. et al. Alcohol and tobacco consumption affects bacterial richness in oral cavity mucosa biofilms. BMC Microbiol. 14, 250 (2014).

56. Kahharova, D. et al. Maturation of the Oral Microbiome in Caries-Free Toddlers: A Longitudinal Study. J. Dent. Res. 99, 159–167 (2020).

57. Moskovitz, M. et al. Characterization of the Oral Microbiome Among Children With Type 1 Diabetes Compared With Healthy Children. Front. Microbiol. 12, (2021).

58. Halboub, E. et al. Tongue microbiome of smokeless tobacco users. BMC Microbiol. 20, 201 (2020).

59. Caselli, E. et al. Defining the oral microbiome by whole-genome sequencing and resistome analysis: the complexity of the healthy picture. BMC Microbiol. 20, 120 (2020).

