## Supplementary Document: Supplementary Texts and Figures for "Multi-cohort analysis of 37,739 oral microbiomes reveals ecologically influential health-associated microbial sub-communities across major oral subsites"

Author Affiliations

#### Text S1

#### Determination of lists of non-sparse consensus taxa associated with the different oral subsite categories

Here, we describe the approach to identify the lists of non-sparse taxa associated with the different subsites. The approach and the results pertaining to the selection of these different lists for the various subsites are described below:

a) ***Salivary subsite***: Consensus salivary-associated taxa were identified using a two-threshold optimisation framework adapted from our previous gut microbiome meta-analysis approach[1]. The framework was applied to identify taxa that were both reproducibly detected across study-cohorts and collectively representative of the salivary microbial community.

   Two thresholds were evaluated as mentioned above.

   (i) The first threshold represented the minimum percentage of samples within a given study-cohort in which a taxon had to be detected (Detection Threshold, which we hereafter refer to as 'X').

   (ii) The second threshold represented the minimum percentage of study-cohorts in which that taxon had to satisfy the within-cohort detection threshold (Study Threshold, which we hereafter refer to as 'Y').

   Thus, for each combination of thresholds, a taxon was retained only if it was detected in at least X% of samples within at least Y% of salivary study-cohorts.

   For each combination of X and Y, a corresponding taxon set was generated. The size of this taxon set was recorded, and its cumulative representation across all the salivary microbiomes was evaluated by summing the relative abundances of the selected taxa within each sample. Threshold combinations were then compared based on two criteria:

   i. Proportion of salivary study-cohorts in which the mean cumulative abundance of the selected taxa combination was $\geq 90\%$ (Property A).

   ii. Proportion of salivary study-cohorts the mean cumulative abundance of the selected taxa combination was $< 70\%$ (Property B).

   The objective was to identify a taxa set with the minimum size that had the highest value of Property A and the lowest value of Property B. The selected taxa set was of 499 taxa, corresponding to taxa detected in at least 5% of samples within at least 15% of salivary study-cohorts. This taxon set cumulatively represented $\geq 90\%$ of the salivary microbial abundance in 95% of study-cohorts (N=94), while representing <70% abundance in only 1% of study-cohorts (N=1). This selected taxon set cumulatively represented approximately 95% of the total salivary microbial abundance, indicating that the dominant fraction of the salivary microbiome could be captured by a reproducible consensus taxon set (**Figure S1; Table S4A**). The same threshold-optimisation framework was then applied independently to the remaining oral subsites (as described below).

b) ***Supragingival subsite***: Adopting the same approach, he optimised threshold retained 301 taxa detected in at least 5% of samples within at least 25% of study-cohorts, which together captured approximately 91.3% of total supragingival microbial abundance (**Figure S2; Table S4B**).

c) ***Subgingival subsite***: The optimised threshold retained 196 taxa detected in at least 5% of samples within at least 45% of study-cohorts which captured approximately 81% of cumulative community abundance, indicating weaker saturation relative to the other oral subsites (**Figure S3; Table S4C**).

d) ***Tongue–tonsil-related subsite***: The above approach retained 266 taxa detected in at least 5% of samples within at least 30% of study-cohorts, cumulatively representing approximately 94% of the total subsite-specific microbial abundance (**Figure S4; Table S4D**).

e) ***Buccal-palate-other surface subsite***: We retained 166 taxa detected in at least 5% of samples within at least 55% of study-cohorts, capturing approximately 90% of the total microbial abundance (**Figure S5; Table S4E**).

Together, these subsite-specific consensus taxon sets provided the taxonomic basis for downstream analyses of health-association, core-association, stability, and integrated taxon ranking.

For clarity, some of the important definitions noted above are clarified below:

- For each taxon, detection was first calculated independently within each study-cohort. For taxon *i* in study-cohort *j*, detection was defined as:

$$Detection_{i,j} = \frac{\textit{number of samples in cohort j with non-zero abundance of taxon i}}{\textit{total number of samples in cohort j}}$$

- Mean abundance was calculated for each taxon within each study-cohort:

$$Mean\ abundance_{ij} = mean\ abundance\ of\ taxon\ i\ across\ all\ samples\ in\ cohort\ j$$

For every candidate taxon set, cumulative community representation was calculated within each study-cohort by summing the mean abundances of all retained taxa:

$$Representation_j = \Sigma\, Mean\ abundance_{ij}, \text{ for all retained taxa } i$$

This value represented how much of the mean microbial community in cohort $j$ was captured by the candidate taxon set.

For each threshold combination, three summary values were calculated:

$$Number\ of\ retained\ taxa = N$$

$$Representation \geq 90\% = \frac{number\ of\ cohorts\ with\ Representation_j \geq 0.90}{total\ number\ of\ cohorts}$$

$$Representation < 70\% = \frac{number\ of\ cohorts\ with\ Representation_j < 0.70}{total\ number\ of\ cohorts}$$

#### Text S2

#### Computation of Health-Association, Core-Association, and Stability-Association Scores

***Health-Association Scores (HS)***: HS were computed to quantify how consistently each taxon was associated with control or disease status across matched case-control study-cohorts. The analysis was performed at the study-cohort level, rather than after pooling all samples, so that each cohort contributed an independent association signal. This design reduced the influence of cohort-specific sample size differences, disease composition, sequencing strategy, and population structure.

For each matched case-control cohort, control and diseased samples were first separated. Cohorts with fewer than 10 control samples or fewer than 10 diseased samples were skipped to avoid unstable statistical comparisons.

For every taxon within each cohort, abundance differences between control and disease samples were tested using the Wilcoxon rank-sum test using the base R (v4.3.1) *wilcox.test* function. The direction of association was determined using Cohen's d effect size, calculated using the *cohen.d* function from the *effsize* (v0.8.1) R package. A positive Cohen's *d* indicated higher abundance in control samples, whereas a negative Cohen's *d* indicated higher abundance in diseased samples.

Each taxon-cohort comparison was assigned a signed association category based on the direction of Cohen's d and the Wilcoxon p value:

+3: significantly increased in controls at $p \leq 0.05$

+2: increased in controls at $0.05 < p \leq 0.10$

+1: increased in controls but not statistically significant at $p > 0.10$

−1: increased in disease but not statistically significant at $p > 0.10$

−2: increased in disease at $0.05 < p \leq 0.10$

−3: significantly increased in disease at $p \leq 0.05$

Only taxon-cohort associations significant at $p \leq 0.05$ were used for HS recurrence counts: values > +2 were counted as significantly control-associated, while values < −2 were counted as significantly disease-associated.

To obtain robust HS rankings, the analysis was repeated over 10 iterations. In each iteration, 65% of matched case-control cohorts were randomly selected. For each taxon, we counted how many selected cohorts showed significant control enrichment (*SP*) and how many showed significant disease enrichment (*SN*). These counts were converted into an iteration-specific unranked HS:

$$HS_{unranked} = \left[\frac{SP - SN}{N}\right] X\ Penalty$$

where *SP* is the number of significantly positive/control-associated cohorts, *SN* is the number of significantly negative/disease-associated cohorts, and *N* is the number of selected study-cohorts in that iteration.

To penalise taxa showing inconsistent bidirectional behavior across cohorts, an imbalance penalty was applied:

$$Penalty = 1 - \left[\frac{\min(SP, SN) + 0.00001}{\max(SP, SN) + 0.00001}\right]$$

This penalty approaches 1 when a taxon shows association in a consistent direction across cohorts and approaches 0 when positive and negative signals are balanced. Therefore, taxa repeatedly increased in controls receive higher positive scores, taxa repeatedly increased in disease receive lower scores, and taxa showing mixed control- and disease-associated signals are down-weighted.

After all iterations, the unranked scores from each iteration were independently rank-scaled to a normalised range of 0–1 using a custom rank_scale function:

$$rank\ scaled\ score = \frac{[rank(x) - \min(rank(x))]}{[\max(rank(x)) - \min(rank(x))]}$$

The final Health-Association Score for each taxon was calculated as the mean of its rank-scaled scores across the 10 iterations. Higher HS values indicate taxa more consistently enriched in control samples across matched case-control cohorts, whereas lower HS values indicate taxa more consistently enriched in diseased samples. Intermediate values represent taxa with weak, inconsistent, or cohort-specific disease-control associations. This framework is also shown in **Figure S6**.

***Core-Association Scores (CS)***: Core-Association Scores (CS) were computed to identify taxa that were reproducibly prevalent and strongly associated with overall community structure across non-diseased study-cohorts. This analysis was performed using only control samples and was applied to the subsite-specific consensus taxon set.

The Core-Association framework was based on two complementary criteria:

(i) Within-cohort prevalence: This captured whether a taxon was consistently detected across samples within a given study-cohort. For each cohort, prevalence was calculated as the proportion of samples in which a taxon had non-zero abundance.

(ii) Community-structure association: This captured whether a taxon was strongly related to the overall composition of the remaining microbial community. This was estimated using the

Remove–Renormalise–Relate (3R) framework. For each taxon, the taxon was removed from the community abundance matrix, the remaining taxa were renormalised within each sample, and Bray-Curtis dissimilarity was calculated on the taxon-removed community profile using *vegdist* from the *vegan* (v2.7.5) package. Principal coordinate analysis was then performed using *dudi.pco* from the *ade4* (v1.7.24) package, and the abundance of the removed taxon was related back to the reconstructed community ordination using *envfit* from the *vegan* package. This generated an envfit-derived $R^2$ value and permutation-based p value for each taxon within each study-cohort. The envfit-derived $R^2$ values were then rank-scaled across all taxa to generate a ranked $R^2$ value for each taxa.

For each cohort, the core-associated taxa were then defined as those that satisfied two properties:

a. They were detected at a prevalence greater than a specific prevalence cutoff threshold.
b. They were influential members of the community. In other words, their ranked envfit-$R^2$ value was above a $R^2$ threshold.

The prevalence and $R^2$ cutoff thresholds were generated as described below:

- ***Prevalence threshold***: For prevalence-threshold selection, multiple detection cutoffs ranging from 0.05 to 0.95 were evaluated. For a given cutoff, taxa detected above that cutoff within a study-cohort were retained as candidate prevalent taxa. Two properties were then calculated across that cohort's samples: the median Jaccard similarity across all sample pairs (using this taxa subset) and the median cumulative abundance of this taxa subset across samples. These two properties were computed for every study-cohort, and their distributions across cohorts are shown as paired boxplots at each cutoff: light-brown for median Jaccard similarity and green for median cumulative abundance (Refer to **Figures S8A**, **S16A**, **S19A**, **S22A** and **S25A** for salivary, supragingival, subgingival, tongue-tonsil and buccal-palate-other surface sites, respectively). For low prevalence thresholds, while the identified core taxa would always account for a large proportion of the microbiome, there is going to be high variation in their detection rates across microbiomes. For stringent or high prevalence thresholds, the selected small taxa set would show minimal variation across samples but would also capture only a small proportion of the microbiome. An ideal core-taxa set should be both uniformly represented across samples (high Jaccard similarity) and account for a large proportion of overall microbiome composition (high cumulative abundance); the optimal cutoff is one that jointly maximises both properties.

- ***Ranked $R^2$ threshold*:** The primary objective to optimise this threshold was to ensure that for a study-cohort, taxa selected as having ranked Ranked $R^2$ above that threshold were also the ones whose abundance variation showed significant association with the community composition of the members. Because absolute $R^2$ values can differ across cohorts, $R^2$ values were first rank-scaled within each cohort. Candidate rank-scaled $R^2$ cutoffs ranging from 0.05 to 0.95 were evaluated. The threshold was thus chosen to balance two aspects: a threshold set too high would retain only taxa consistently associated with other members' abundances, but would miss taxa with significant community-composition associations that nonetheless have lower ranked $R^2$ (reduced sensitivity); a threshold set too low would misclassify taxa with non-significant associations as ecologically influential (increased false positives). The ideal threshold therefore maximises accuracy, the proportion of correctly identified taxa with significant community-composition associations.

The optimised values of the above two parameters were selected independently each subsite specific study-cohorts (as summarised in **Figures S8**, **S16**, **S19**, **S22** and **S25** for salivary, supragingival, subgingival, tongue-tonsil and buccal-mucosa-other sites, respectively).

Using the optimised subsite-specific thresholds, taxa satisfying both criteria were classified as core-associated within that study-cohort. For each taxon, core-associated recurrence was calculated as:

$$\text{core-associated recurrence} = \frac{No.of\ cohorts\ in\ which\ the\ taxon\ is\ core\ associated}{total\ number\ of\ eligible\ cohorts}$$

These recurrence values were then rank-scaled from 0 to 1 to generate the final Core-Association Score (CS). Higher CS values indicate taxa that were more reproducibly identified as core-associated across independent non-diseased study-cohorts. The same framework is shown in **Figure S8C**.

**Stability-Association Scores (SS):** Stability-Association Scores (SS) were computed to identify salivary taxa whose baseline abundance was reproducibly associated with longitudinal microbiome stability. This analysis was restricted to the salivary subsite because longitudinal microbiome profiles were available only for this subsite.

For each longitudinal study-cohort, samples from the same subject were paired with their immediate follow-up sample. Microbiome change between the current and follow-up time point was quantified as follow-up distance. Two distance measures were used for the final stability score: (i) Bray–Curtis distance and (ii) an Aitchison-proxy distance, calculated as Euclidean distance on relative-abundance profiles. Bray–Curtis distances were calculated using *vegdist* from the *vegan* package. Lower follow-up distance indicated higher longitudinal microbiome stability.

For each taxon, the association between baseline abundance and follow-up distance was estimated separately within each study-cohort using robust linear regression:

$$Followup\ distance\ = \beta_0 + \beta_1 \times\ taxon\ abundance$$

Robust linear models were fitted using *rlm* from the *MASS* (v7.3.60) package, and statistical significance of the taxon term was evaluated using robust F-tests. In this framework, a negative association indicated that higher baseline abundance of a taxon was associated with lower follow-up distance, and therefore greater microbiome stability.

To summarise associations across cohorts, cohort-level effects were combined using random-effects meta-analysis. Effect sizes were transformed using Fisher's Z-transformed partial correlation (*ZPCOR*) with *escalc*, and pooled using *rma* from the *metafor* (v5.0.1) package. For each taxon, the meta-analysis generated a pooled effect estimate, p value, false-discovery-rate-adjusted q value, and a consistency measure. Consistency was defined as the proportion of cohorts in which the direction of the cohort-level effect matched the direction of the pooled meta-analytic effect.

The stability scoring procedure was repeated over 10 iterations. In each iteration, 8 longitudinal cohorts were randomly selected, and the robust regression plus random-effects meta-analysis workflow was applied independently for each taxon. For each iteration, taxon-level stability evidence was calculated by combining effect direction, statistical strength, and cross-cohort consistency:

$$Stability\ evidence\ = (-\beta) \times\ [-log_{10}(q)] \times\ consistency$$

Here, β represents the pooled meta-analytic effect estimate. The negative sign was applied so that taxa negatively associated with follow-up distance received higher stability evidence. Each iteration-specific stability evidence vector was rank-scaled from 0 to 1.

The above procedure was performed separately for Bray–Curtis and Aitchison-proxy follow-up distances. For each taxon, the mean rank-scaled stability score across iterations was calculated for each distance metric. The final Stability-Association Score was then computed as the average of the Bray–Curtis-based and Aitchison-proxy-based mean stability scores:

$$SS = mean(SS\ BrayCurtis, SS\ Aitchison\ proxy)$$

Higher SS values indicate taxa whose baseline abundance was more reproducibly associated with reduced longitudinal microbiome change across independent salivary longitudinal cohorts. **Figure S11** depicts this framework.

These Scores calculated for each subsite are given in **Tables S11**, **S17**, **S21**, **S25**, **S29** for salivary, supragingival, subgingival, tongue-tonsil and buccal-palate-other surface site respectively.

**Computation of Scores for stratified study-cohorts in salivary subsite**

For the salivary subsite, association scores were recomputed within stratified cohort subsets to assess reproducibility across sequencing technologies and exposure contexts. Health-Association Scores and Core-Association Scores were recomputed separately using 16S cohorts, WGS cohorts, and exposure-associated cohorts. Stability-Association Scores were recomputed within sequencing-specific longitudinal subsets where sufficient longitudinal data were available.

The stratified scores were compared with the corresponding overall scores using correlation analyses to evaluate whether taxon-level rankings were robust across data-generation modalities and exposure-associated datasets. Stratified analyses were restricted to the salivary subsite because other oral subsites had fewer eligible cohorts after partitioning by sequencing strategy, exposure status, or longitudinal sampling availability. The figures associated with this analysis are depicted in **Figures S7**, **S10**, **S13**, **S14** for HS, CS, SS and sHACK respectively and **Tables S6**, **S8, S10** for HS, CS, SS respectively for salivary subsite.

#### Text S3

##### Description of the different salivary taxa groups that show variable association patterns with Health-Association and Core-Association

Using these two scores, taxa in the saliva-sputum-oral wash subsite were stratified into four groups based on their scores. Of these, the Quadrant1 (or Q1) group, which showed high Health-Association and Core-Association (HS >= 0.70 and CS >= 0.70) was the most important in terms of diagnostic and therapeutic promise. The description of this group is already provided in the main manuscript.

In contrast, Quadrant2 (or Q2) group contained taxa with high Core-Association scores (CS >= 0.70) and lower Health-Association scores (HS < 0.70) (**Table S8B**). This group included several core-associated taxa like *Prevotella* spp. (*P. salivae*, *P. oulorum*, *P. loescheii*, *P. maculosa*, *P. denticola*, *P. nanceiensis*), *Leptotrichia spp.* (*L. wadei*, *L. goodfellowii*, *L. hongkongensis*, *L. trevisanii*, *L. shahii*), and taxa from diverse oral biofilm-associated genera, including *Oribacterium sinus*, *Gemella sanguinis*, *Corynebacterium matruchotii*, *Stomatobaculum longum*, *Atopobium parvulum*, *Capnocytophaga sputigena*, *Veillonella parvula*, and *Dialister invisus* (**Figure 2A**).

The Quadrant3 (Q3) group consisted of taxa with both low CS and low HS (CS & HS <0.7; **Table S8B**). This represented the largest and most heterogeneous group of taxa, comprising species that were neither reproducibly core-associated across control salivary cohorts nor consistently health-associated across matched case-control cohorts. This quadrant included a mixture of peripheral oral taxa, low-recurrence oral commensals, gut-associated anaerobes, environmental or transient taxa, and several host- or niche-associated species not typically considered stable members of the salivary core. The presence of multiple gut-prevalent lineages, including members of *Bacteroides*, *Faecalibacterium*, *Roseburia*, *Ruminococcus*, *Alistipes*, *Parabacteroides*, *Coprococcus*, and *Eubacterium*, suggests that this quadrant captures taxa with limited ecological integration into the salivary microbiome rather than a coherent oral health-associated consortium (**Figure 2A**). Thus, low CS and low HS scores likely reflect weak recurrence, inconsistent disease-control directionality, or transient detection across cohorts.

The last taxa group, Quadrant4 or Q4, contained taxa with low-CS and high-HS (CS <0.7 & HS >=0.7; **Table S8B**). This contained taxa that were strongly associated with health but were not consistently identified as core-associated members across control salivary cohorts. Representative species included *Streptococcus lactarius*, *Streptococcus australis*, *Neisseria* *subflava*, *Neisseria sicca*, *Kingella denitrificans*, *Actinomyces massiliensis*, *Prevotella aurantiaca*, *Capnocytophaga haemolytica*, and *Veillonella tobetsuensis*. Several taxa, such as *Prevotella scopos*, *Moryella indoligenes*, *Haemophilus haemoglobinophilus*, *Prevotella fusca*, and *Desulfomicrobium orale*, showed high health-association despite near-zero core association, suggesting context-dependent or cohort-specific health-linked patterns.

#### Text S4

#### Integrated sHACK and HAC score computation

Integrated taxon-level scores were computed to prioritise taxa showing concordant associations across multiple microbiome properties. For the salivary subsite, where all three component scores were available, the salivary Health-Associated Core Keystone score (sHACK) was computed by integrating Core-Association Score (CS), Health-Association Score (HS), and Stability-Association Score (SS). For each taxon, the three scores were first combined by taking their mean, representing the overall strength of association across the three dimensions.

To ensure that high sHACK values were assigned only to taxa with balanced contributions across all three properties, a Gini-based concordance term was applied. The Gini coefficient measures inequality among the component scores; therefore, taxa with one high score but low values for the remaining scores were penalised. The sHACK score was calculated as:

$$sHACK = mean(CS, HS, SS) \times [1 - Gini(CS, HS, SS)]$$

where CS, HS, and SS represent the Core-Association Score, Health-Association Score, and Stability-Association Score, respectively. Gini coefficients were computed using the *Gini* function from the *DescTools* (v0.99.60) package. The resulting sHACK values were then rank-scaled from 0 to 1 to generate the final salivary sHACK ranking.

For all oral subsites, an analogous Health-Associated Core score (HAC) was computed using only CS and HS:

$$HAC = mean(CS, HS) \times [1 - Gini(CS, HS)]$$

Thus, sHACK prioritised taxa that were simultaneously core-associated, health-associated, and stability-associated in the salivary microbiome, whereas HAC prioritised taxa that were simultaneously core-associated and health-associated in the remaining oral subsites. However, to compare the salivary subsite with other subsites, we also calculated the HAC score for salivary subsite was also calculated. This framework has been depicted in the **Figure 3A**. The results for this analysis have been given in the **Tables S11**, **S17B**, **S21B**, **S25B** for salivary, supragingival, subgingival, and tongue-tonsil site respectively.

**Text S5**

**Complete protocol for the identification of taxonomic modules (or robust clusters of taxa) in the salivary microbiomes**

We employed two complementary approaches for this purpose. First, we identified groups of taxa exhibiting similar patterns of association with all other members of the salivary microbial community across studies, using an association-score-based approach adopted from a previous study by our group[2]. Second, we constructed taxon-to-taxon networks linking pairs of taxa that showed significant co-abundance relationships with each other (**Figure S15A**). For this, we generated abundance-based, across-study "meta-networks" by performing random-effects-model-based meta-analysis across all study-cohorts to identify taxon pairs showing significant correlations in abundance[2]. These approaches are described in detail below

a. *Approach 1 – Creation of two-dimensional maps:* For any given pair of taxa (i, j), we first computed Spearman correlations between their abundances independently within each study-cohort. The number of cohorts in which the taxon pair showed a significant positive correlation (Spearman's $\rho > 0$; FDR $\leq 0.1$) and a significant negative correlation (Spearman's $\rho < 0$; FDR $\leq 0.1$) were denoted as SP and SN, respectively. The Association-Score between taxa i and j was then calculated as:

   Association-Score(i,j)
   = ((SP – SN)/T) * (1-((Min(SP,SN)+0.0001)/(Max(SP,SN)+0.0001))

   Where, T = Total number of study-cohorts, Min(SP,SN) = Minimum of SP and SN and Max(SP,SN) = Maximum of SP and SN.

This procedure was applied to all pairwise combinations of the 499 salivary microbiome taxa, yielding a 499 × 499 Association-Score matrix capturing the pairwise association pattern of each taxon with every other taxon. Study-specific correlation analysis was performed using *corr.test* functionality of the *psych* package version-2.6.1 in R (https://cran.r-project.org/web/packages/psych/index.html). Principal Component Analysis (PCA) was then applied to this Association-Score matrix, and each taxon was represented as a point in two-dimensional space defined by the first two principal components (PC1 on the x-axis, PC2 on the y-axis). In this representation, taxa with similar Association-Score profiles across all other taxa, i.e., similar patterns of co-occurrence with the broader community, are positioned closer together. Principal Component Analysis was performed using the *dudi.pca* function of the ade4 package version-1.7.23 in R (https://cran.r-project.org/web/packages/ade4/index.html).

Clustering taxa based on their proximity in this 2-D space therefore is expected to identify modules of taxa likely to share similar ecological roles or functional traits within the salivary microbial community. We utilised the k-means clustering approach for this purpose. The k-means clustering approach is a method of grouping points (or observations) into 'k' non-overlapping clusters, where in each observation is assigned to the cluster whose centroid is having the least distance to the observation. The key parameter to decide here is selection of the appropriate 'k' that achieves the best partitioning of the observations (or points).

To decide this, we performed 20 iterations. In each iteration, we selected 50% of the taxa and performed the clustering of the points corresponding to the selected taxa (in the 2-D space obtained as above) using various values of k, ranging from 2 to 10. For each value of k, we measured the efficiency of clustering of the points using Calinski-Harabasz (CH) index. The CH indices for each value of k across the 20 iterations were then compared using boxplots to select minimum value of 'k' achieving the highest CH index across the iterations with the minimum variation across iterations. Based on the patterns observed here, we selected k=5 for partitioning the taxa into five taxonomic modules, where in each taxa belonging to each of the modules showed similar patterns of association with all other taxa across the 15,127 salivary microbiomes from non-diseased individuals across 99 cohorts. The R packages, *cluster* version-2.1.8.2 and *clusterSim* version-0.51.6, were utilised for performing the k-means clustering of the points and the related investigations (https://cran.r-project.org/web/packages/cluster/index.html; https://cran.r-project.org/web/packages/clusterSim/index.html). The results pertaining to this clustering or module are given in **Figures S15B**, **S18B**, **S21B**, **S24B** for salivary, supragingival, subgingival, and tongue-tonsil site respectively where the number of clusters or modules are selected based on the highest CH value.

**b.** *Approach 2 – Creation of taxa-to-taxa meta-networks*: Here we generated co-abundance-based taxon-to-taxon meta-network. Given a pair of taxa, we identified whether the two exhibited significant co-abundance relationships (higher abundance associated with higher abundance of the other) by performing a Random Effect Model based meta-analysis of their abundances across the different study-cohorts. We specifically considered only the non-diseased gut microbiomes for this purpose.

Within each cohort, the effect-size of the co-abundance relationships was obtained by performing a robust linear regression of the abundance of one microbe based on the abundance of the other. The robust linear regression model was built using the *rlm* function of the *MASS* package version-7.3.65 in R (https://cran.r-project.org/web/packages/MASS/index.html). The robust linear model outputs obtained for the individual study-cohorts were then provided as inputs to the *rma* function of the *metafor* package version-4.8.0 in R (https://cran.r-project.org/web/packages/metafor/index.html). The *mutate* function of the *dplyr* package version-1.2.0 was utilised to parse the outputs of the individual *rlm* models into the input format required by *rma* function for computing the Random Effect Models.

For a given taxa-pair, the summarised estimate, the P-value, and the consistency (i.e the fraction of the study-cohorts where the directionality robust linear regression estimate of the relationships were the same as that of the summarised estimate) were obtained. The above steps were repeated for all taxa-pairs. The P-values obtained for all the individual pairs were then corrected to Q-values using the Benjamini-Hochberg correction. All taxa-pairs having a summarised Random Effect Model estimate > 0, Q-value <= 0.0001 for salivary subsite (Q-value <= 0.05 for non-salivary subsite) and consistency >= 0.70, were identified to have a significant co-abundance relationship and were connected by an edge. This resulted in the creation of the taxa-to-taxa meta-network in the discovery cohort. This edge list was further used to get the graphical meta-network using Cytoscape (v3.10.4) using Compound Spring Embedder (CoSE) layout. These network results are depicted in **Figures 3G, 6B, 6E, and 6I** for the salivary, supragingival, subgingival, and tongue–tonsil subsites, respectively. The corresponding edge lists are provided in **Tables S13, S19, S23, and S27**, respectively.

**Text S6**

**Complete protocol for investigation the network similarity between the discovery and validation cohort derived meta-network**

Investigation of the similarity between the discovery cohort derived and validation cohort derived meta-networks was performed using two approaches:

**Fishers' Exact Test based evaluation for edge-overlap between the two meta-networks**

To assess whether the co-abundance edges identified in the discovery-cohort meta-network were reproduced in the validation-cohort meta-network more than expected by chance, we performed a Fisher's exact test on edge overlap between the two networks. Each network was represented as an edge list restricted to a common node universe, and edges were compared as unordered node pairs to determine the number of edges present in both networks (overlap), present in only one network, or absent from both. Given n shared nodes, the total number of possible node pairs was calculated as $^{n}C_{2}$., and edge presence/absence in each network was tabulated into a 2×2 contingency table (edge present/absent in the discovery network × edge present/absent in the validation network).

A one-sided Fisher's exact test (alternative = "greater") was applied to this contingency table to test whether the observed edge overlap between the two networks exceeded that expected under the null hypothesis of independence between the two networks, yielding a P-value and an odds ratio quantifying the strength of enrichment. Fisher's exact tests were implemented in base R using the *fisher.test* function.

**Quadratic Assignment Procedure (QAP) for network similarity testing**

The QAP approach was utilised to assess the correspondence between the discovery- and validation-cohort meta-networks. For this purpose, each meta-network was first represented as a binary adjacency matrix, and both matrices were restricted to the set of nodes common to both networks, with node ordering aligned identically across matrices. The Pearson correlation between the two adjacency matrices was computed as the observed test statistic, capturing the degree of correspondence between edge patterns across the two networks.

To assess the significance of this observed correlation, a null distribution was generated by randomly permuting node labels within each network (10,0000 permutations). The approach preserved each network's internal structure, while recomputing the matrix correlation at each permutation. This node-label permutation strategy accounted for the non-independence of edges within a network, an assumption violated by simpler edge-overlap tests. This makes QAP a more conservative and appropriate test of network-level correspondence. A one-sided empirical P-value was calculated as the proportion of permuted correlations equal to or exceeding the observed correlation, testing the alternative hypothesis that the two networks are more similar than expected under random node correspondence. QAP tests were implemented using the *qaptest* function from the *sna* package version-2.8 in R.

The results pertaining to this validation is shown in **Figure 4A**.

#### Text S7

#### Computation of Health-Association, Core-Association, and HAC Rankings, and Identification of Ecological Modules in the Supragingival Subsite

We next examined the supragingival subsite to evaluate whether health-associated core taxa could also be identified in a distinct tooth-associated oral habitat. The supragingival dataset included 10,381 microbiomes from 24 study-cohorts, comprising 9,168 non-diseased and 1,213 diseased microbiomes. Using the subsite-specific consensus taxon selection framework, 301 supragingival consensus taxa were retained for downstream analyses (**Text S1**; **Table S4B**).

Core-Association Scores were computed using 9,168 non-diseased supragingival microbiomes from 23 study-cohorts (**Table S16A**), following the framework described in **Text S2**. Using the subsite-specific threshold optimisation framework, the optimised prevalence and community-association thresholds for supragingival CS computation were set at 0.80 and 0.75, respectively (**Figure S16A–B**). The highest core-associated taxa included *Streptococcus sanguinis*, *Fusobacterium nucleatum*, *Rothia dentocariosa*, *Rothia aeria*, and *Porphyromonas catoniae* (**Figure S16C**; **Table S17A**). Additional taxa from *Corynebacterium*, *Capnocytophaga*, *Actinomyces*, and *Neisseria* lineages also showed high Core-Association Scores, indicating their recurrent presence and community association across non-diseased supragingival cohorts.

Health-Association Scores were computed using matched case-control supragingival cohorts spanning 7 disease categories from 2558 non-disease control and matched 1213 diseased microbiome (**Table S16B**), following the iterative cohort-wise framework described in **Text S2**. Taxa with high HS included *Cardiobacterium hominis*, *Leptotrichia goodfellowii*, *Lautropia mirabilis*, *Streptococcus sanguinis*, *Aggregatibacter segnis*, *Prevotella nanceiensis*, *Eikenella corrodens*, *Corynebacterium durum*, *Porphyromonas catoniae*, *Gemella morbillorum*, *Neisseria oralis*, *Haemophilus parahaemolyticus*, and *Lachnoanaerobaculum umeaense* (**Table S17A**). In contrast, low-HS taxa included *Treponema lecithinolyticum*, *Prevotella baroniae*, *Prevotella salivae*, *Prevotella histicola*, *Prevotella nigrescens*, *Fretibacterium fastidiosum*, *Selenomonas sputigena*, *Streptococcus constellatus*, *Neisseria bacilliformis*, and *Actinomyces israelii*.

Because longitudinal data were not sufficiently available for the supragingival subsite, taxa were prioritised using the Health-Associated Core framework, which integrates Health-Association and Core-Association Scores using the Gini-based concordance approach described in **Text S4**. Comparison of HS and CS highlighted 30 taxa with concordant high values across both dimensions, including *Cardiobacterium hominis*, *Streptococcus sanguinis*, *Eikenella corrodens*, *Corynebacterium durum*, *Porphyromonas catoniae*, *Lautropia mirabilis*, *Capnocytophaga ochracea*, and *Veillonella parvula* (**Figure S17**; **Table S17A**).

We identified 26 high-HAC taxa (HAC>=0.90) supragingival taxa. These included *Cardiobacterium hominis*, *Streptococcus sanguinis*, *Eikenella corrodens*, *Corynebacterium durum*, *Porphyromonas catoniae*, *Prevotella loescheii*, *Lautropia mirabilis*, *Capnocytophaga ochracea*, *Prevotella oulorum*, and *Veillonella parvula* (**Figure S18A**; **Table S17B**). Overall, these results suggest that the supragingival HAC framework identifies taxa that are both recurrently core-associated in non-diseased cohorts and consistently health-associated across matched case-control cohorts.

We next asked whether supragingival high-HAC taxa were organised into coherent ecological modules. Using the same community-wide association framework applied for the salivary subsite (**Text S5**), pairwise association profiles among supragingival consensus taxa were first used to generate a two-dimensional association map based on PC1 and PC2 coordinates (**Table S18**). The optimal number of clusters was then determined by iterative clustering analysis, which identified 3 supragingival taxonomic modules as the most stable solution (**Figure S18B**). Projection of these taxa onto the association map showed a clear modular arrangement of the supragingival microbiome (**Figure 6A**).

We then examined the distribution of HAC scores across the three supragingival modules. Module-level comparison showed that module 2 contained taxa with comparatively higher HAC values (**Figure S18C**). To further evaluate whether these modules also reflected underlying ecological structure, we constructed the supragingival co-abundance meta-network using 845 edges (**Table S19**), visualised using the Compound Spring Embedder (CoSE) layout in Cytoscape and coloured taxa by module identity. The resulting network showed cluster-wise organisation consistent with the association-map-derived modules, providing additional support that these supragingival modules represent biologically meaningful microbial sub-communities rather than arbitrary clusters (**Figure 6B**). Consistent with this pattern, 21 high-HAC taxa (HAC $\geq$ 0.90) were significantly enriched in module 2 (odds ratio = 21.15, Fisher's exact test, P= 1.4e-11; **Figure 6C**). Notably, all 21 of these taxa also showed concordantly high component scores, with both HS and CS $\geq$ 0.70. These 21 taxa belonging to the high HAC supragingival ecological module-2 were *Cardiobacterium hominis*, *Cardiobacterium valvarum*, *Eikenella corrodens*, *Neisseria flavescens*, *Lautropia mirabilis*, *Rothia aeria*, *Corynebacterium durum*, *Actinomyces georgiae*, *Actinomyces naeslundii*, *Gemella morbillorum*, *Streptococcus sanguinis*, *Campylobacter showae*, *Capnocytophaga ochracea*, *Capnocytophaga sputigena*, *Capnocytophaga granulosa*, *Capnocytophaga leadbetteri*, *Porphyromonas catoniae*, *Prevotella loescheii*, *Prevotella asaccharolytica* and *Aggregibacter segnis*.

#### Text S8

#### Computation of Health-Association, Core-Association, and HAC Rankings, and Identification of Ecological Modules in the Subgingival Subsite

We next examined the subgingival subsite, a distinct tooth-associated habitat with a more anaerobe-enriched community structure. The subgingival dataset included 1,974 microbiomes from 11 study-cohorts, comprising 1,418 non-diseased and 556 diseased microbiomes. Using the subsite-specific consensus taxon selection framework, 196 subgingival consensus taxa were retained for downstream analyses (**Text S1**; **Table S4C**).

Core-Association Scores were computed using 1,418 non-diseased subgingival microbiomes from 11 study-cohorts (**Table S20A**), following the framework described in **Text S2**. Using the subsite-specific threshold optimisation framework, the optimised prevalence and community-association thresholds for CS computation were set at 0.75 and 0.75, respectively (**Figure S19A-B**). The highest core-associated taxa included *Rothia dentocariosa*, *Fusobacterium nucleatum*, *Capnocytophaga* spp., *Parvimonas micra*, *Eubacterium brachy*, and *Porphyromonas* spp. (**Figure S19C**; **Table S21A**). These results indicate that the subgingival core-associated set included taxa recurrently detected and strongly associated with community structure across non-diseased subgingival cohorts.

Health-Association Scores were computed using matched case-control subgingival cohorts of containing 990 non-disease control and 556 diseased microbiomes spanning 4 diseases (**Table S20B**), following the iterative cohort-wise framework described in **Text S2**. The highest health-associated taxa included *Capnocytophaga sputigena*, *Rothia aeria*, *Neisseria elongata*, *Capnocytophaga gingivalis*, *Capnocytophaga leadbetteri*, *Corynebacterium matruchotii*, *Lautropia mirabilis*, *Neisseria bacilliformis*, *Cardiobacterium hominis*, *Corynebacterium durum*, *Leptotrichia buccalis*, *Capnocytophaga granulosa*, and *Kingella oralis* (**Table S21A**).

Because longitudinal data were not sufficiently available for the subgingival subsite, taxa were prioritised using the Health-Associated Core (HAC) framework, which integrates Health-Association and Core-Association Scores using the Gini-based concordance approach described in **Text S4**. Comparison of HS and CS highlighted taxa with concordant high values across both dimensions, including *Capnocytophaga gingivalis*, *Kingella oralis*, *Capnocytophaga leadbetteri*, *Lautropia mirabilis*, *Capnocytophaga granulosa*, *Capnocytophaga sputigena*, *Corynebacterium durum*, and *Corynebacterium matruchotii* (**Figure S20**; **Table S21A**).

The top HAC-ranked subgingival taxa included *Capnocytophaga gingivalis*, *Kingella oralis*, *Capnocytophaga leadbetteri*, *Lautropia mirabilis*, *Capnocytophaga granulosa*, *Capnocytophaga sputigena*, *Corynebacterium durum*, *Streptococcus sanguinis*, *Capnocytophaga ochracea*, and *Corynebacterium matruchotii* (**Figure S21A**; **Table S21B**). Overall, these results suggest that the subgingival HAC framework identifies taxa that combine recurrent Core-Association with consistent health-association across matched case-control cohorts.

We next evaluated whether high-HAC taxa in the subgingival subsite also showed module-level organisation similar to supragingival and salivary subsite. Using the same community-wide association framework described for the salivary and supragingival analyses (**Text S5**), pairwise association profiles among subgingival consensus taxa using the non-disease control cohort, were constructed to get a two-dimensional association map based on PC1 and PC2 coordinates (**Table S22**). Iterative clustering analysis identified 2 subgingival taxonomic modules as the optimal clustering solution (**Figure S21B**). Mapping these modules onto the association space showed a clear separation of subgingival taxa into two major ecological groups (**Figure 6D**).

We then compared HAC score distributions between the two subgingival modules. This analysis showed that module 1 contained taxa with comparatively higher HAC values (**Figure S21C**). To further assess whether these association-map-derived modules were reflected in co-abundance structure, we constructed a subgingival co-abundance meta-network containing 131 edges (**Table S23**), visualised using the Compound Spring Embedder (CoSE) layout in Cytoscape and coloured taxa by module identity. The resulting network showed module-wise organisation consistent with the association-map clustering, supporting the presence of distinct subgingival microbial sub-communities (**Figure 6E**). In agreement with this pattern, 16 taxa with HAC $\geq 0.90$ were significantly enriched in module 1 (odds ratio =15.3, Fisher's exact test, P= 5.0e-4) (**Figure 6F**) of which all taxa have both the scores (HS & CS) >=0.70 (**Figure S21A**). These 16 taxa were *Veillonella parvula*, *Streptococcus mitis*, *Streptococcus sanguinis*, *Actinomyces massiliesis*, *Rothia aeria*, *Corynebacterium durum*, *Corynebacterium matruchotii*, *Cardiobacterium hominis*, *Lautropia mirabilis*, *Kingella oralis*, *Porphyromonas catoniae*, *Capnocytophaga sputigena*, *Capnocytophaga granulosa*, *Capnocytophaga gingivalis*, *Capnocytophaga ochracea* and *Capnocytophaga leadbetteri*.

#### Text S9

#### Tongue-tonsil Health Association, Core Association, and HAC ranking

We next examined the tongue–tonsil-related subsite, a mucosal oral habitat with a distinct microbial composition compared with tooth-associated plaque sites. The tongue-tonsil dataset included 1,693 microbiomes from 18 study-cohorts, comprising 1,570 non-diseased and 123 diseased microbiomes. Using the subsite-specific consensus taxon selection framework, 266 tongue–tonsil consensus taxa were retained for downstream analyses (**Text S1**; **Table S4D**).

Core-Association Scores were computed using 1,570 non-diseased tongue-tonsil microbiomes from 18 study-cohorts (**Table S24A**), following the framework described in **Text S2**. Using the subsite-specific threshold optimisation framework, the optimised prevalence and community-association thresholds for CS computation were set at 0.80 and 0.75, respectively (**Figure S22A-B**). The highest core-associated taxa included members of *Veillonella*, *Prevotella*, *Actinomyces*, *Megasphaera*, and *Streptococcus* lineages, including *Veillonella dispar*, *Veillonella atypica*, *Prevotella salivae*, and *Actinomyces odontolyticus* (**Figure S22C**; **Table S25A**). These results indicate that the tongue–tonsil core-associated set was dominated by taxa recurrently detected and strongly associated with community structure across non-diseased cohorts.

Health-Association Scores were computed using matched case-control tongue–tonsil cohorts spanning 3 diseases across 4 cohorts (**Table S24B**), following the iterative cohort-wise framework described in **Text S2**. The highest health-associated taxa included *Mogibacterium diversum*, *Porphyromonas gulae*, *Alloprevotella rava*, *Neisseria subflava*, *Neisseria perflava*, *Lachnospiraceae bacterium oral taxon 096*, *Prevotella sp. oral taxon 306*, *Tannerella sp. oral taxon HOT 286*, *Actinomyces sp. ICM47*, and *Streptococcus sp. A12* (**Table S25A**).

Because longitudinal data were not sufficiently available for the tongue–tonsil subsite, taxa were prioritised using the Health-Associated Core framework, which integrates Health-Association and Core-Association Scores using the Gini-based concordance approach described in **Text S4**. Comparison of HS and CS highlighted a very smaller set of taxa with concordant high values across both dimensions, including *Mogibacterium diversum*, *Eubacterium sulci*, *Alloprevotella rava*, *Neisseria flavescens*, and *Prevotella jejuni* (**Figure S23**; **Table S25A**).

The top HAC-ranked tongue–tonsil taxa (HAC >=0.90) included *Mogibacterium diversum*, *Alloprevotella rava*, *Eubacterium sulci*, *Neisseria flavescens*, *Prevotella jejuni*, and *Rothia mucilaginosa* (**Figure S24A**; **Table 25B**).

We next evaluated whether high-HAC taxa in the tongue–tonsil subsite also showed module-level organisation. Using the same community-wide association framework described for the salivary, supragingival, and subgingival analyses (**Text S5**), pairwise association profiles among tongue-tonsil consensus taxa from non-diseased control cohorts were constructed to generate a two-dimensional association map based on PC1 and PC2 coordinates (**Table S26**). Iterative clustering analysis identified 6 tongue–tonsil taxonomic modules as the optimal clustering solution (**Figure S24B**). Projection of these modules onto the association space showed that tongue–tonsil taxa were organised into multiple distinct ecological groups (**Figure 6G**).

We then compared HAC score distributions across the six tongue–tonsil modules. Although module 6 showed comparatively higher HAC values at the module level (**Figure 6H**), high-HAC taxa were not significantly enriched in this module when using the HAC $\geq$ 0.90 cutoff (odds ratio = 1.76, Fisher's exact test, P= 0.26; **Figure S24C**). This reflects the small number of high-HAC taxa in module 6, with only 3 taxa meeting the HAC $\geq$ 0.90 threshold. Thus, while module 6 showed the highest overall HAC distribution, the enrichment of high-HAC taxa did not reach statistical significance. To further assess whether these association-map-derived modules were reflected in co-abundance structure, we constructed a tongue-tonsil co-abundance meta-network containing 420 edges (**Table S27**), visualised using the Compound Spring Embedder (CoSE) layout in Cytoscape and coloured taxa by module identity. The resulting network showed module-wise organisation consistent with the association-map clustering, supporting the presence of distinct tongue–tonsil microbial sub-communities (**Figure 6I**).

#### Text S10

#### Buccal–palate-other mucosal surface Core-Association and disease-control association patterns

The last subsite with very a smaller number of samples, buccal–palate-other mucosal surface subsite, representing oral epithelial surfaces distinct from tooth-associated plaque and tongue–tonsil habitats were also examined. The buccal-palate-other surface dataset included 749 microbiomes from 11 study-cohorts, comprising 521 non-diseased and 221 diseased microbiomes. Using the subsite-specific consensus taxon selection framework, 166 buccal–palate-other mucosal consensus taxa were retained for downstream analyses (**Table S4E**).

Core-Association Scores were computed using 521 non-diseased buccal–palate-other mucosal microbiomes from 10 study-cohorts (**Table 28A**), following the framework described in **Text S2**. Using the subsite-specific threshold optimisation framework, the optimised prevalence and community-association thresholds for CS computation were set at 0.75 and 0.75, respectively (**Figure S25A–B**). The highest core-associated taxa included *Rothia mucilaginosa*, *Fusobacterium nucleatum*, *Actinomyces odontolyticus*, *Haemophilus parainfluenzae*, and *Streptococcus mitis* (**Figure S25C**; **Table S29A**). These results indicate that the buccal–palate-other mucosal core-associated set included taxa recurrently detected and associated with community structure across non-diseased mucosal cohorts.

Because only two matched disease-control cohorts with total 275 microbiome (non-diseased = 54; diseased = 221; **Table S28B**) were available for this subsite, Health-Association Scores and HAC rankings were not computed. Instead, we examined cohort-wise disease-control association patterns directly within the available matched cohorts. Taxa showing control-associated patterns included only one species - *Prevotella shahii*, whereas taxa showing disease-associated patterns included *Actinomyces naeslundii*, *Capnocytophaga gingivalis*, *Capnocytophaga sputigena*, *Capnocytophaga leadbetteri*, and *Streptococcus sanguinis*. (**Figure S25D**; **Table S29B**).

No other analysis was carried due to limited number of samples in this subsite.

#### Text S11

#### Random Forest-based Prediction of sHACK/HAC Score Using Genome-Derived Species Level Functional profile

Genome-derived species-level functional profiles were used to evaluate whether the functional potential of oral taxa could explain their integrated sHACK/HAC rankings. Briefly, 71,238 high-quality reference genomes and metagenome-assembled genomes were curated from publicly available genome resources, including EMBL-EBI MGnify and the UNINA MAGs Genome Repository[3]. Genome annotation was performed using the eggNOG-mapper[4] pipeline against the eggNOG database[5], with protein-coding sequences predicted using Prodigal[6] as implemented within the annotation workflow. Functional features were summarised across multiple annotation categories, including CAZy[7], COG[8], BiGG[9], KEGG[10] orthologs, KEGG reactions, KEGG modules, EC numbers[11], and Pfam domains[12]. For species represented by multiple genomes, feature values were averaged across genomes to generate a single species-level functional profile. Thus, each species was represented by a vector of predicted functional capacities, reflecting its mean functional potential across its available genomes.

For the oral microbiome analysis, these species-level functional profiles were matched to the subsite-specific consensus taxa. This yielded matched functional profiles for 366 salivary taxa, 220 supragingival taxa, 140 subgingival taxa, and 190 tongue-tonsil taxa, for which both species-level-taxa functional profile as well as the subsite-specific HAC/sHACK scores were available.

For each subsite, a functional input matrix was defined as:

$$F = [f_{ij}]$$

where each row $i$ represented a species and each column $j$ represented a functional group. The response vector was defined as:

$$Y = [y_i]$$

where $y_i$ represented the observed integrated taxon-level score for species $i$. For the salivary subsite, $Y$ corresponded to the sHACK score, whereas for supragingival, subgingival, and tongue-tonsil subsites, $Y$ corresponded to the HAC score.

For each subsite, an initial random forest regression model was fitted using all matched functional groups as predictors and the observed sHACK/HAC score as the response:

$$Y_{\mathrm{i}} = RF(f_{\mathrm{i}1}, f_{\mathrm{i}2}, \ldots, f_{\mathrm{i}p})$$

where $p$ represents the total number of functional groups available in the matched functional matrix. Random forest regression was performed using the *randomForest* (v4.7.1.2) package in R, and functional-feature importance was estimated using percentage increase in mean squared error (%IncMSE). Features with higher %IncMSE values were considered more important because permuting those features resulted in a larger reduction in predictive performance. Model performance was evaluated by comparing the random forest predicted scores with the observed sHACK/HAC scores using Pearson correlation:

$$r = cor\left(Y_{predicted}, Y_{observed}\right)$$

To identify a compact predictive functional signature, functional groups were ranked according to %IncMSE, and random forest models were re-trained using progressively smaller sets of top-ranked features. Feature-set sizes were evaluated from 250 to 20 features, decreasing by 5 features at each step:

$$N = 250, 245, 240, \ldots, 20$$

For each value of *N*, the top *N* functional groups were retained and a new random forest model was fitted:

$$Y_{\mathrm{i}} = RF(top\ N\ functional\ groups)$$

For every model, the correlation between predicted and observed sHACK/HAC scores was calculated, along with the corresponding P value. The final feature set was selected by balancing predictive performance and parsimony, prioritising the smallest feature set that retained high predicted-versus-observed correlation. This procedure selected 40 functional groups for the salivary subsite, 25 functional groups for the supragingival subsite, 30 functional groups for the subgingival subsite, and 20 functional groups for the tongue-tonsil subsite and their corresponding selection is depicted in **Figures S26**, **S27A**, **S28A**, **S29A** respectively.

Using the final selected feature set, a random forest model was refitted, and predicted-versus-observed sHACK/HAC scores were visualised to assess model performance. This Predicted vs actual results are given in **Tables S30A**, **S31A**, **S32A**, **S33A** for salivary, supragingival, subgingival, tongue-tonsil, buccal-palate-other surface site respectively. While their correlation of actual vs predicted using selected features is visualised in **Figures 8B-E**.

To interpret the selected features, each selected functional group was then correlated independently with the observed sHACK/HAC score across species:

$$r_j = cor(F_j, Y)$$

where $F_j$ represents the values of functional group $j$ across species. The resulting p values were corrected using false-discovery-rate (fdr) adjustment to obtain Q-values. Functional groups with $Q \leq 0.05$ were considered significantly associated with the score. Positive correlations indicated functional groups enriched among higher-scoring taxa, whereas negative correlations indicated functional groups associated with lower-scoring taxa.

Selected functional groups were summarised into broad biological categories based on their annotations, including phosphate acquisition and regulation, ion transport and homeostasis, proteolysis and hydrolysis, sugar transport and exchange, stress resistance, DNA repair and replication, cell-envelope and surface-associated functions, signal transduction and transcriptional regulation, mobile genetic elements, oxidative stress response, carbon and amino-acid metabolism, translation and tRNA metabolism. These associations were visualised using volcano plots, with correlation coefficient on the x-axis and $-\log_{10}$(Q-value) on the y-axis. This framework is also depicted in **Figure 8A**. The Volcano plots showing association of selected features with sHACK/HAC is given in the **Figures 8F**, **S27B**, **S28B**, **S29B** for salivary, supragingival, subgingival, tongue-tonsil, buccal-palate-other surface site respectively.

#### Text S12

##### Functional-profile-based prediction and interpretation of subsite-specific sHACK/HAC scores

To determine whether genome-derived functional potential could explain the integrated taxon-level prioritisation scores, we used species-level functional profiles to predict sHACK/HAC scores across oral subsites. Functional profiles were available for 366 salivary taxa, 220 supragingival taxa, 140 subgingival taxa, and 190 tongue-tonsil taxa after matching subsite-specific consensus taxa with the genome-derived functional-profile matrix. For each subsite, random forest regression models were trained using functional groups as predictors and the corresponding sHACK/HAC score as the response. Functional features were first ranked using %IncMSE, and reduced models were then evaluated across progressively smaller top-feature sets from 250 to 20 features. The final selected models retained high observed–predicted score correlations while using comparatively compact functional feature sets. For detailed methodology see **Text S11** and **Figure 8A**.

##### *Salivary sHACK scores were predicted by a compact set of genome-derived functional features*

For the salivary subsite, 366 taxa had matched genome-derived functional profiles and sHACK scores. Random forest models trained across progressively reduced feature sets showed consistently high observed-predicted correlations across multiple feature numbers. Although larger models also performed well, the 40-feature model retained high predictive performance while using a substantially smaller feature set (**Figure S26**). This selected 40-feature model achieved an observed-predicted Pearson correlation of $r = 0.71$ with a highly significant association between predicted and observed sHACK scores ($P = 2.8e-56$) (**Figure 8B**; **Table S30A**). The selected model therefore indicated that a limited subset of genome-derived functional groups was sufficient to capture a major component of the salivary sHACK ranking.

Correlation analysis of these selected 40 functional features with observed salivary sHACK scores identified 35 FDR-significant features at $Q \leq 0.05$ (**Table S30B**). Among these, six features were positively associated with sHACK scores, whereas 29 features showed negative associations (**Figure 8B**). The strongest positive associations were observed for Sugar transport and exchange features, including sugar efflux transporter for intercellular exchange ($r = 0.46$, $Q = 9.8e-20$) and SemiSWEET-family sugar transporter ($r = 0.44$, $Q = 3.2e-18$). SemiSWEET/SWEET transporters are known mono- and disaccharide transporters, and bacterial SemiSWEET homologs have been structurally shown to form transporter pores with conserved residues required for glucose transport activity[13,14]. Therefore, the positive association of sugar efflux and SemiSWEET-family transporter annotations with sHACK suggests that high-sHACK salivary taxa may possess greater predicted capacity for carbohydrate movement, intercellular substrate exchange, and metabolic flexibility within salivary microbial communities.

Additional positive associations involved Stress resistance and methylation features, including tellurite methyltransferase, Tellurite resistance protein TehB, SAM-dependent methylase, Tellurite resistance protein TehB, as well as Ankyrin repeat domains classified under Protein interaction domains. Tellurite-related methyltransferase and TehB-like annotations are generally associated with bacterial detoxification or tolerance to toxic metalloid stress[15–17]. The bacteria *Haemophilus parainfluenzae* has the most potential to metal stress resistance[18] which is one of the highest sHACK taxa. Therefore, their positive association with sHACK suggests that high-sHACK taxa may carry selected protective stress-tolerance capacities that could support persistence under chemically or oxidatively variable salivary conditions. Ankyrin repeat domains may further reflect protein-protein interaction potential, which could contribute to host interaction, surface adaptation, or community-level persistence[19], although their specific role in salivary health-associated taxa requires experimental validation.

In contrast, negatively associated features were dominated by Phosphate acquisition and regulation. Thirteen significant negatively associated features belonged to this category (**Table S30B**), including phosphate transport system permease proteins, ABC-type phosphate transport components, phosphate uptake regulator PhoU, phosphate transport system ATP-binding components, and PhoR/PhoB phosphate-regulon elements. The strongest negative associations included phosphate transport system permease protein ($r = -0.50$, $Q = 1.6e-22$), ABC-type phosphate transport system permease component ($r = -0.50$, $Q = 1.6e-22$), phosphate transport system module ($r = -0.50$, $Q = 1.7e-22$), and PhoU phosphate uptake regulator ($r = -0.49$, $Q = 2.1e-22$). These features represent conserved bacterial phosphate-starvation and high-affinity phosphate-acquisition systems, including the PhoR/PhoB regulatory system, PhoU phosphate uptake regulator, and ABC-type phosphate transport machinery[20–24]. Their inverse association with sHACK suggests that high-sHACK salivary taxa were less enriched for functions typically required under phosphate limitation or nutrient-stress conditions. The reasons for these associations are not immediately evident and will require deeper functional investigations pertaining to the mechanistic basis of these associations. One possible explanation is that taxa with greater phosphate acquisition or regulatory potential may be better equipped to persist under nutrient-limited or environmentally perturbed conditions, including inflammatory or dysbiotic environments. Although direct evidence linking phosphate acquisition to persistence in such environments is lacking, analogous phosphorus-acquisition and adaptation strategies have been documented in microbial communities from phosphorus-limited terrestrial and aquatic ecosystems[25–27]

Other negatively associated themes included Ion transport and homeostasis, Proteolysis and hydrolysis, Cell-envelope-associated functions, Cell-envelope stress response, Signal transduction, ATP-dependent functions, Oxidoreductase activity, Mobile genetic elements, and Amino acid/2-oxocarboxylic acid metabolism. Within the proteolysis and hydrolysis, choloylglycine hydrolase and related taurocholate and glycocholate amidohydrolase annotations correspond to bile salt hydrolase-like enzymes that hydrolyse conjugated bile acids into unconjugated bile acids and glycine or taurine[28].

Together, these results indicated that salivary sHACK scores were functionally structured. Higher-scoring taxa were associated with predicted functional capacities related to sugar transport and exchange, selected metalloid/stress-tolerance mechanisms, and protein-interaction domains, whereas several phosphate-starvation, ion-transport, proteolytic, envelope-stress, and nutrient-scavenging functions were inversely associated with sHACK ranking. However, because these annotations were derived from genome-based functional profiles, they should be interpreted as predicted functional potential rather than direct evidence of in vivo expression or activity. We require further investigations to validate and establish the mechanistic bases behind their sHACK association patterns.

##### *Supragingival HAC scores were predicted by 25 functional features enriched for sugar transport and regulatory domains*

For the supragingival subsite, 220 taxa had matched genome-derived functional profiles and HAC scores. Random forest models trained using progressively reduced feature sets showed robust observed-predicted correlations across the feature-reduction series. The selected 25-feature model achieved an observed-predicted Pearson correlation of $r = 0.69$ ($P = 6.1e-32$), providing a parsimonious functional model for supragingival HAC score prediction (**Figure S27A**; **Figure 8C**; **Table S31A**). Although models with larger feature sets showed similar or slightly higher correlations, the 25-feature model retained high predictive performance with a much smaller functional signature.

Among the selected 25 functional features, 20 were significantly associated with supragingival HAC scores after FDR correction (**Table S31B**; **Figure S27B**). Eight significant features showed positive correlations with HAC scores. Similar to the salivary subsite, the strongest positive associations were related to Sugar transport and exchange, including a SemiSWEET-family sugar transporter (r = 0.37, Q = 1.9e-7) and a sugar efflux transporter for intercellular exchange (r = 0.36, Q = 3.2e-7). These features indicate predicted capacity for carbohydrate movement across bacterial membranes and intercellular substrate exchange[13,14]. The positive association of these features with supragingival HAC suggests that higher-HAC taxa may carry functional potential related to carbohydrate uptake, sugar efflux, and substrate exchange within tooth-associated microbial communities.

This result is biologically relevant because supragingival plaque is a structured biofilm that is repeatedly exposed to host- and diet-derived carbohydrates[29]. In this environment, carbohydrate transport does not necessarily indicate a single health or disease state; rather, it reflects an important ecological capacity that can contribute to microbial growth, substrate sharing, and biofilm-level metabolic interactions[30,31]. Oral biofilm studies have shown that microbial communities respond dynamically to sugar availability, and that carbohydrate metabolism, cross-feeding, and metabolic exchange can shape community organisation[32,33]. Since HAC integrates Health-Association and Core-Association Scores, these results suggest that supragingival health-associated core taxa may be functionally characterised, in part, by predicted capacities for sugar transport and intercellular metabolic exchange.

Additional positive associations included a GyrI-like small-molecule binding domain, a cyclopropanoid cyclopropyl hydrolase/GyrI-like domain linked to lipid modification, and isoleucyl-tRNA synthetase. These features may reflect complementary functional capacities related to small-molecule binding, membrane or lipid adaptation, and translation-associated cellular maintenance. In a surface-attached supragingival biofilm, such functions may help bacteria respond to changing local conditions, maintain cellular activity, and persist within a stable microbial community[34,35]. Although the specific roles of these features in supragingival health-associated taxa require experimental validation, their positive association with HAC suggests that higher-HAC taxa are not only taxonomically distinct but also carry a coherent set of predicted functional capacities related to substrate exchange, cellular maintenance, and biofilm adaptation.

Twelve significant features were inversely associated with supragingival HAC scores. The strongest inverse associations were related to Nitrogen and ammonium transport, including an ammonium transporter family feature (r = -0.37, Q = 1.9e-7) and an ammonia channel protein AmtB (r = -0.37, Q = 1.9e-7) (**Table S31B**). Again, while the exact mechanistic bases behind these associations require further validation, taxa with lower HAC scores may rely more strongly on ammonium transport and nitrogen-scavenging strategies, whereas higher-HAC taxa may be characterised more by sugar transport and exchange-related functions.

Other inversely associated features included methanogenesis (r = -0.32, Q = 6.0e-6), aspartate, methionine, tyrosine aminotransferase (r = -0.30, Q = 1.6e-5), N-ribosyl hydrolase reaction class (r = -0.30, Q = 1.6e-5) (**Table S31B**), and sugar-acid dehydration reaction class (r = -0.30, Q = 2.6e-5). Additional inverse functional themes included nucleotide and nucleoside metabolism, carbohydrate metabolism, fatty acid biosynthesis, acetate production and energy metabolism, and ribosome-associated translation features. Together, these annotations suggest that supragingival taxa differed in predicted biosynthetic activity, nitrogen handling, energy metabolism, nucleotide turnover, and translational capacity across the HAC ranking. These functions may reflect alternative metabolic strategies used by different members of the supragingival biofilm, particularly under changing nutrient, oxygen, or community-structure conditions.

Overall, supragingival HAC prediction was supported by a compact functional signature in which higher-HAC taxa were positively associated with Sugar transport and exchange, together with selected small-molecule binding, lipid-adaptation, and translation-related features. Inversely associated features highlighted differences in nitrogen/ammonium transport, energy metabolism, amino-acid transformation, nucleotide metabolism, and ribosome-associated functions across HAC-ranked taxa. These results suggest that supragingival HAC scores capture biologically meaningful variation in genome-derived functional potential, with high-HAC taxa showing predicted capacities consistent with substrate exchange, metabolic flexibility, and persistence within structured tooth-associated biofilms. As these annotations were derived from genome-based functional profiles, they should be interpreted as predicted functional potential rather than direct evidence of in vivo activity.

***Subgingival HAC scores showed the strongest functional predictability and were associated with amino acid, polyamine, redox, and cell-surface functions***

For the subgingival subsite, 140 taxa had matched genome-derived functional profiles and HAC scores. Random forest models across the feature-reduction series showed strong predictive performance, and the selected 30-feature model achieved the highest observed-predicted correlation among the tested compact models. This model produced an observed–predicted Pearson correlation of r = 0.74 (P = 1.1e-25) (**Figure S28A**; **Figure 8D**; **Table S32A**). This represented the strongest functional prediction performance among the four analysed oral subsites, suggesting that subgingival HAC scores were particularly well captured by genome-derived functional features.

Correlation analysis of the 30 selected functional features identified 16 FDR-significant features, all of which were positively associated with subgingival HAC scores (**Table S32B**; **Figure S28B**). The strongest positive associations involved amino acid and tetrapyrrole biosynthesis, arginine and polyamine metabolism, oxidative stress response, regulatory domains, and cell-surface-associated features. Glutamate-1-semialdehyde 2,1-aminomutase showed the strongest positive association ($r = 0.33$, $Q = 7.0e-4$), followed closely by arginase family ($r = 0.33$, $Q = 7.0e-4$) and arginase/agmatinase family enzyme ($r = 0.32$, $Q = 7.0e-4$). These features suggested a positive relationship between higher subgingival HAC scores and functional capacities related to amino acid conversion, arginine metabolism, and polyamine-linked processes.

The enrichment of arginine-related features among higher-HAC subgingival taxa is biologically relevant because arginine metabolism is an important component of oral biofilm physiology. Arginine can be metabolised by oral bacteria to generate alkali, contributing to pH homeostasis and helping buffer acidification within dental biofilms. Experimental oral biofilm studies have also shown that arginine can modulate microbial metabolism and community structure, supporting its broader role in maintaining biofilm ecological balance. Therefore, the positive association of arginase and arginase, agmatinase-family features with subgingival HAC scores may indicate that health-associated core taxa in the subgingival niche carry predicted capacities linked to nitrogen handling, arginine conversion, and polyamine-related metabolism[36,37].

The association with glutamate-1-semialdehyde 2,1-aminomutase and glutamate-1-semialdehyde aminotransferase suggests an additional link between higher subgingival HAC scores and amino-acid, tetrapyrrole biosynthetic potential. These enzymes are involved in aminolevulinate formation, a precursor step connected to tetrapyrrole and heme-related biosynthesis. In the subgingival environment, where oxygen gradients, heme availability, redox conditions, and host-derived inflammatory products strongly shape microbial ecology, such biosynthetic capacities may support cellular maintenance and adaptation among health-associated core taxa. Prior subgingival and periodontal microbiome studies have shown that periodontal status is associated with functional shifts in microbial gene expression and metabolic potential, highlighting the relevance of metabolic activity and site-specific function in this habitat[38,39].

Additional positively associated features included periplasmic deferrochelatase/peroxidase EfeB (r = 0.32, Q = 7.0e-4), a WYL_2 domain protein (r = 0.32, Q = 7.0e-4), a PKD domain (r = 0.32, Q = 7.0e-4), and glutamate-1-semialdehyde aminotransferase (r = 0.31, Q = 7.5e-4) (**Table S32B**; **Figure S28B**). The EfeB feature is particularly notable because EfeB-like proteins have been described as peroxidase/deferrochelatase-related proteins involved in oxidative-stress defense and iron-associated processes. In the subgingival niche, bacteria are exposed to fluctuating oxygen tension, host inflammatory responses, heme/iron sources, and reactive oxygen species. Therefore, the positive association of EfeB with HAC suggests that higher-HAC taxa may carry predicted functional potential for redox buffering or oxidative-stress tolerance[40,41], which could support persistence in the dynamic subgingival environment.

The WYL_2 and TetR-associated WHG domains point toward regulatory and environmental-response capacities among higher-HAC taxa. WYL domains are commonly associated with bacterial regulatory proteins, while TetR-family regulators are broadly involved in bacterial transcriptional control, stress responses, antimicrobial resistance, and biofilm-associated traits. In oral bacteria, TetR-family regulation has been linked to biofilm formation and stress-response pathways, supporting the interpretation that these features may contribute to adaptive regulation in structured plaque communities[42,43].

The PKD domain and cell-surface carbohydrate biosynthesis feature further suggest that higher-HAC subgingival taxa may possess capacities related to surface interaction, adhesion, and cell-envelope organisation. These functions are relevant in subgingival plaque, where stable colonisation depends on adhesion to tooth/root surfaces, interaction with neighboring microbes, and adaptation to the gingival crevice environment. Consistent with this, prior subgingival microbiome studies have emphasised that periodontal health and disease states are associated not only with taxonomic shifts but also with changes in microbial functional potential, biofilm-associated traits, signal transduction, and host-microbiome interactions[39].

Other significant positive associations included MFS multidrug transporter, DNA polymerase III alpha, predicted SnoaL-like aldol condensation enzyme, phosphohistidine swiveling domain, cytochrome/quinol oxidase subunit, dihydrolipoyl dehydrogenase, and alkyl/prenyl-group transfer reaction class. These annotations indicate that the subgingival HAC-associated functional signature extended beyond a single pathway and included transport, replication, redox metabolism, regulatory signaling, membrane/cell-surface biology, and biosynthetic functions. This is consistent with the ecological complexity of subgingival plaque, where bacteria must respond to host-derived nutrients, oxygen gradients, inflammatory pressure, redox changes, and dense biofilm interactions. Studies integrating subgingival microbiome and metabolome data have similarly shown that periodontal states are associated with changes in amino-acid biosynthesis, carbohydrate-related metabolism, pyrimidine metabolism, and other metabolic pathways, supporting the biological relevance of these functional categories[44,45].

Overall, the subgingival predictive signature was characterised by positive associations with metabolism, redox activity, cell-surface interaction, regulatory functions, and transport-related features, with no selected feature showing a significant negative association after FDR correction. These results suggest that higher subgingival HAC scores were linked to a coordinated genome-derived functional profile involving amino-acid conversion, arginine, polyamine-related metabolism, oxidative-stress tolerance, regulatory adaptation, surface interaction, and cellular maintenance. Since HAC integrates Health-Association and Core-Association Scores, these findings indicate that subgingival health-associated core taxa may be distinguished not only by their recurrence and health association, but also by predicted functional capacities relevant to persistence and adaptation within the subgingival biofilm. As these annotations were derived from genome-based functional profiles, they should be interpreted as predicted functional potential rather than direct evidence of in vivo activity.

***Tongue-tonsil HAC scores were predicted by a minimal 20-feature model with mixed positive and negative functional associations***

For the tongue-tonsil subsite, 190 taxa had matched genome-derived functional profiles and HAC scores. Across progressively reduced functional feature sets, random forest models retained moderate-to-high predictive performance. The selected 20-feature model achieved an observed-predicted Pearson correlation of $r = 0.63$ ($P = 3.6e-22$) while using the smallest final feature set among the analysed subsites (**Figure S29A**; **Figure 8E**; **Table S33A**).

Among the selected 20 features, 11 were significantly associated with tongue-tonsil HAC scores after FDR correction (**Table S33B**; **Figure S29B**). Three features showed positive associations. These included an ArsR family arsenate, arsenite, antimonite-responsive transcriptional regulator ($r = 0.20$, $Q = 0.02$), an ankyrin repeat domain ($r = 0.18$, $Q = 0.03$), and a predicted ATPase ($r = 0.17$, $Q = 0.04$). These features linked higher tongue-tonsil HAC scores to metal-resistance-associated transcriptional regulation, protein-interaction repeat domains, and ATP-dependent functions.

The positive association of the ArsR family arsenate, arsenite, antimonite-responsive transcriptional regulator suggests that higher-HAC tongue-tonsil taxa may carry regulatory capacity for responding to metalloid stress. ArsR-family regulators are widely involved in bacterial transcriptional responses to toxic metal ions and metalloids, including arsenite and related compounds, and therefore indicate predicted potential for environmental sensing and stress-responsive regulation. In the tongue-tonsil habitat, where microbes are exposed to saliva, dietary compounds, oxygen gradients, epithelial shedding, host immune factors, and transient environmental inputs, such regulatory functions may support persistence of health-associated core taxa under fluctuating conditions. This interpretation is consistent with the broader role of tongue-associated microbiota as a dynamic oral habitat with measurable interpersonal and temporal variation in healthy individuals[35].

The ankyrin repeat domain positively associated with HAC may reflect protein–protein interaction or adaptation-related potential among higher-HAC tongue–tonsil taxa. Because bacterial ankyrin-repeat proteins can contribute to host interaction, environmental adaptation, and surface-associated processes, this feature was interpreted as a broad protein-interaction signal rather than evidence of a specific mechanism[46,47]. Interestingly, similar functions were also found to be high-sHACK-associated for the salivary microbiomes. Similarly, the predicted ATPase may indicate ATP-dependent cellular processes such as transport, protein remodelling, stress adaptation, or cellular maintenance. Given the broad nature of this annotation, it was interpreted as a general ATP-dependent functional signal among higher-HAC taxa rather than a pathway-specific marker.

Eight significant features were inversely associated with tongue-tonsil HAC scores (**Table S33B**; **Figure S29B**). These included an AAA domain of the Cdc48 subfamily (r = -0.22, Q= 0.02), ribosome-associated features such as ribosomal protein L31 (r = -0.20 to -0.21, Q= 0.02) and ribosomal protein L28 (r = -0.20, Q == 0.02), as well as putative DNA primase/helicase (r = -0.20, Q= 0.02) and preprotein translocase subunit SecG (r = -0.16, Q= 0.05). These features indicate that tongue-tonsil taxa differed across the HAC gradient in predicted ATP-dependent protein remodelling, translation, DNA replication/repair, and Sec-dependent protein translocation capacities.

Overall, the tongue-tonsil functional signature was distinct from the other oral subsites. Higher tongue-tonsil HAC scores were positively associated with metalloid-responsive transcriptional regulation, protein-interaction domains, and ATP-dependent functions, whereas inverse associations highlighted variation in protein remodelling, ribosome-associated translation, DNA primase-helicase activity, and Sec-dependent protein translocation. These results suggest that tongue-tonsil HAC scores capture habitat-specific differences in genome-derived functional potential related to environmental sensing, regulatory adaptation, protein interaction, and cellular processing. As with the other subsites, these annotations should be interpreted as predicted functional potential rather than direct evidence of in vivo activity.

***Cross-subsite comparison of functional predictive signatures***

Across oral subsites, genome-derived functional profiles consistently predicted integrated taxon-level scores with significant observed-predicted correlations. This demonstrated that sHACK/HAC rankings were not only taxonomically structured but also reflected measurable differences in species-level genome-encoded functional potential.

Despite this shared predictability, the functional themes associated with high scores differed across subsites. Salivary and supragingival high-score taxa shared positive associations with sugar transport and exchange features, particularly SemiSWEET-family and sugar efflux transporter annotations (**Figures 8F**, **S27B**). In contrast, subgingival high-HAC taxa were characterised by positive associations with amino acid/tetrapyrrole biosynthesis, arginine and polyamine metabolism, oxidative stress response, cell-surface interaction domains, and redox/energy-related functions. Tongue-tonsil high-HAC taxa showed positive associations with metal-responsive transcriptional regulation, ankyrin-repeat protein interaction domains, and ATPase-related functions. These results indicate that high-priority oral taxa have subsite-specific functional signatures, with distinct functional capacities contributing to sHACK/HAC score prediction in different oral habitats.

**References:**


1. Goel, A. *et al.* Toward a health-associated core keystone index for the human gut microbiome. *Cell Rep.* **44**, (2025).
2. Ghosh, T. S., Shanahan, F. & O'Toole, P. W. Toward an improved definition of a healthy microbiome for healthy aging. *Nat. Aging* **2**, 1054–1069 (2022).
3. Mitchell, A. L. *et al.* MGnify: the microbiome analysis resource in 2020. *Nucleic Acids Res.* **48**, D570–D578 (2020).
4. Cantalapiedra, C. P., Hernández-Plaza, A., Letunic, I., Bork, P. & Huerta-Cepas, J. eggNOG-mapper v2: Functional Annotation, Orthology Assignments, and Domain Prediction at the Metagenomic Scale. *Mol. Biol. Evol.* **38**, 5825–5829 (2021).
5. Huerta-Cepas, J. *et al.* eggNOG 5.0: a hierarchical, functionally and phylogenetically annotated orthology resource based on 5090 organisms and 2502 viruses. *Nucleic Acids Res.* **47**, D309–D314 (2019).
6. Hyatt, D. *et al.* Prodigal: prokaryotic gene recognition and translation initiation site identification. *BMC Bioinformatics* **11**, 119 (2010).
7. Drula, E. *et al.* The carbohydrate-active enzyme database: functions and literature. *Nucleic Acids Res.* **50**, D571–D577 (2022).
8. Galperin, M. Y. *et al.* COG database update: focus on microbial diversity, model organisms, and widespread pathogens. *Nucleic Acids Res.* **49**, D274 (2020).
9. Norsigian, C. J. *et al.* BiGG Models 2020: multi-strain genome-scale models and expansion across the phylogenetic tree. *Nucleic Acids Res.* **48**, D402–D406 (2020).
10. Kanehisa, M., Furumichi, M., Sato, Y., Kawashima, M. & Ishiguro-Watanabe, M. KEGG for taxonomy-based analysis of pathways and genomes. *Nucleic Acids Res.* **51**, D587 (2022).
11. Duvaud, S. *et al.* Expasy, the Swiss Bioinformatics Resource Portal, as designed by its users. *Nucleic Acids Res.* **49**, W216 (2021).
12. Mistry, J. *et al.* Pfam: The protein families database in 2021. *Nucleic Acids Res.* **49**, D412 (2020).
13. Xu, Y. *et al.* Structures of bacterial homologues of SWEET transporters in two distinct conformations. *Nature* **515**, 448–452 (2014).
14. Jia, B., Hao, L., Xuan, Y. H. & Jeon, C. O. New Insight Into the Diversity of SemiSWEET Sugar Transporters and the Homologs in Prokaryotes. *Front. Genet.* **9**, (2018).

15. Peng, W. *et al.* The antibacterial effect of tellurite is achieved through intracellular acidification and magnesium disruption. *mLife* **4**, 423–436 (2025).
16. Gaete-Argel, A. *et al.* Tellurite Promotes Stress Granules and Nuclear SG-Like Assembly in Response to Oxidative Stress and DNA Damage. *Front. Cell Dev. Biol.* **9**, (2021).
17. Sandoval, J. M., Levêque, P., Gallez, B., Vásquez, C. C. & Buc Calderon, P. Tellurite-induced oxidative stress leads to cell death of murine hepatocarcinoma cells. *BioMetals* **23**, 623–632 (2010).
18. Kang, Y. *et al.* Dental Plaque Microbial Resistomes of Periodontal Health and Disease and Their Changes after Scaling and Root Planing Therapy. *mSphere* **6**, 10.1128/msphere.00162-21 (2021).
19. Al-Khodor, S., Price, C. T., Kalia, A. & Abu Kwaik, Y. Functional diversity of ankyrin repeats in microbial proteins. *Trends Microbiol.* **18**, 132–139 (2010).
20. Baek, S. *et al.* PhoU interaction with the PhoR PAS domain is required for repression of the pho regulon and Salmonella virulence, but not for polyphosphate accumulation. *J. Microbiol.* **63**, e2505013 (2025).
21. Martín, J. F. & Liras, P. Molecular Mechanisms of Phosphate Sensing, Transport and Signalling in Streptomyces and Related Actinobacteria. *Int. J. Mol. Sci.* **22**, 1129 (2021).
22. Bruna, R. E., Kendra, C. G. & Pontes, M. H. An intracellular phosphorus-starvation signal activates the PhoB/PhoR two-component system in Salmonella enterica. *mBio* **15**, e01642-24 (2024).
23. Lamarche, M. G., Wanner, B. L., Crépin, S. & Harel, J. The phosphate regulon and bacterial virulence: a regulatory network connecting phosphate homeostasis and pathogenesis. *FEMS Microbiol. Rev.* **32**, 461–473 (2008).
24. Lubin, E. A., Henry, J. T., Fiebig, A., Crosson, S. & Laub, M. T. Identification of the PhoB Regulon and Role of PhoU in the Phosphate Starvation Response of Caulobacter crescentus. *J. Bacteriol.* **198**, 187–200 (2015).
25. Wu, X. *et al.* Genome-Resolved Metagenomics Reveals Distinct Phosphorus Acquisition Strategies between Soil Microbiomes. *mSystems* **7**, e01107-21 (2022).
26. Dai, T. *et al.* Nutrient supply controls the linkage between species abundance and ecological interactions in marine bacterial communities. *Nat. Commun.* **13**, 175 (2022).
27. Peñuelas, J., Zheng, B., Tariq, A. & Sardans, J. Microbial phosphorus cycling in terrestrial ecosystems. *Nat. Rev. Microbiol.* **24**, 478–495 (2026).
28. Agolino, G. *et al.* Bile salt hydrolase: The complexity behind its mechanism in relation to lowering-cholesterol lactobacilli probiotics. *J. Funct. Foods* **120**, 106357 (2024).

29. Edlund, A. *et al.* Meta-omics uncover temporal regulation of pathways across oral microbiome genera during in vitro sugar metabolism. *ISME J.* **9**, 2605–2619 (2015).
30. Wu, C. M. *et al.* Mucin glycans drive oral microbial community composition and function. *Npj Biofilms Microbiomes* **9**, 11 (2023).
31. Miller, D. P., Fitzsimonds, Z. R. & Lamont, R. J. Metabolic Signaling and Spatial Interactions in the Oral Polymicrobial Community. *J. Dent. Res.* **98**, 1308–1314 (2019).
32. Miller, D. P., Fitzsimonds, Z. R. & Lamont, R. J. Metabolic Signaling and Spatial Interactions in the Oral Polymicrobial Community. *J. Dent. Res.* **98**, 1308–1314 (2019).
33. Esberg, A., Haworth, S., Hasslöf, P., Lif Holgerson, P. & Johansson, I. Oral Microbiota Profile Associates with Sugar Intake and Taste Preference Genes. *Nutrients* **12**, 681 (2020).
34. Wake, N. *et al.* Temporal dynamics of bacterial microbiota in the human oral cavity determined using an in situ model of dental biofilms. *Npj Biofilms Microbiomes* **2**, 16018 (2016).
35. Hall, M. W. *et al.* Inter-personal diversity and temporal dynamics of dental, tongue, and salivary microbiota in the healthy oral cavity. *Npj Biofilms Microbiomes* **3**, 2 (2017).
36. Huang, X., Schulte, R. M., Burne, R. A. & Nascimento, M. M. Characterization of the Arginolytic Microflora Provides Insights into pH Homeostasis in Human Oral Biofilms. *Caries Res.* **49**, 165–176 (2015).
37. Agnello, M. *et al.* Arginine Improves pH Homeostasis via Metabolism and Microbiome Modulation. *J. Dent. Res.* **96**, 924–930 (2017).
38. Belstrøm, D. *et al.* Periodontitis associates with species-specific gene expression of the oral microbiota. *Npj Biofilms Microbiomes* **7**, 76 (2021).
39. Shi, B. *et al.* The subgingival microbiome associated with periodontitis in type 2 diabetes mellitus. *ISME J.* **14**, 519–530 (2020).
40. Zhang, C. *et al.* Identification of a Novel Dye-Decolourizing Peroxidase, EfeB, Translocated by a Twin-Arginine Translocation System in Streptococcus thermophilus CGMCC 7.179. *Appl. Environ. Microbiol.* **81**, 6108–6119 (2015).
41. Sheldon, J. R., Laakso, H. A. & Heinrichs, D. E. Iron Acquisition Strategies of Bacterial Pathogens. *Microbiol. Spectr.* **4**, 10.1128/microbiolspec.vmbf-0010–2015 (2016).
42. Boutrin, M.-C. *et al.* A putative TetR regulator is involved in nitric oxide stress resistance in Porphyromonas gingivalis. *Mol. Oral Microbiol.* **31**, 340–353 (2016).
43. Liu, J. *et al.* TetR Family Regulator brpT Modulates Biofilm Formation in Streptococcus sanguinis. *PLOS ONE* **12**, e0169301 (2017).

44. Shi, M. *et al.* Alterations and Correlations in Microbial Community and Metabolome Characteristics in Generalized Aggressive Periodontitis. *Front. Microbiol.* **11**, (2020).
45. Lu, X. *et al.* Subgingival microbiome in periodontitis and type 2 diabetes mellitus: an exploratory study using metagenomic sequencing. *J. Periodontal Implant Sci.* **52**, 282–297 (2022).
46. Jernigan, K. K. & Bordenstein, S. R. Tandem-repeat protein domains across the tree of life. *PeerJ* **3**, e732 (2015).
47. Hamilton, W. C. & Newton, I. L. G. crANKing up the infection: ankyrin domains in Rickettsiales and their role in host manipulation. *Infect. Immun.* **92**, e00059-24 (2024).

### Supplementary Figures

#### Multi-cohort analysis of 37,739 oral microbiomes reveals ecologically influential health-associated microbial sub-communities across major oral subsites

Omprakash Shete, Alisha Ansari, Mithun Verma, Abhishek P, Esha Chauhan, Sourav Goswami, Tarini Shankar Ghosh*

Author Affiliations

Department of Computational Biology, Indraprastha Institute of Information Technology (IIIT-Delhi), Okhla Phase III, New Delhi, India

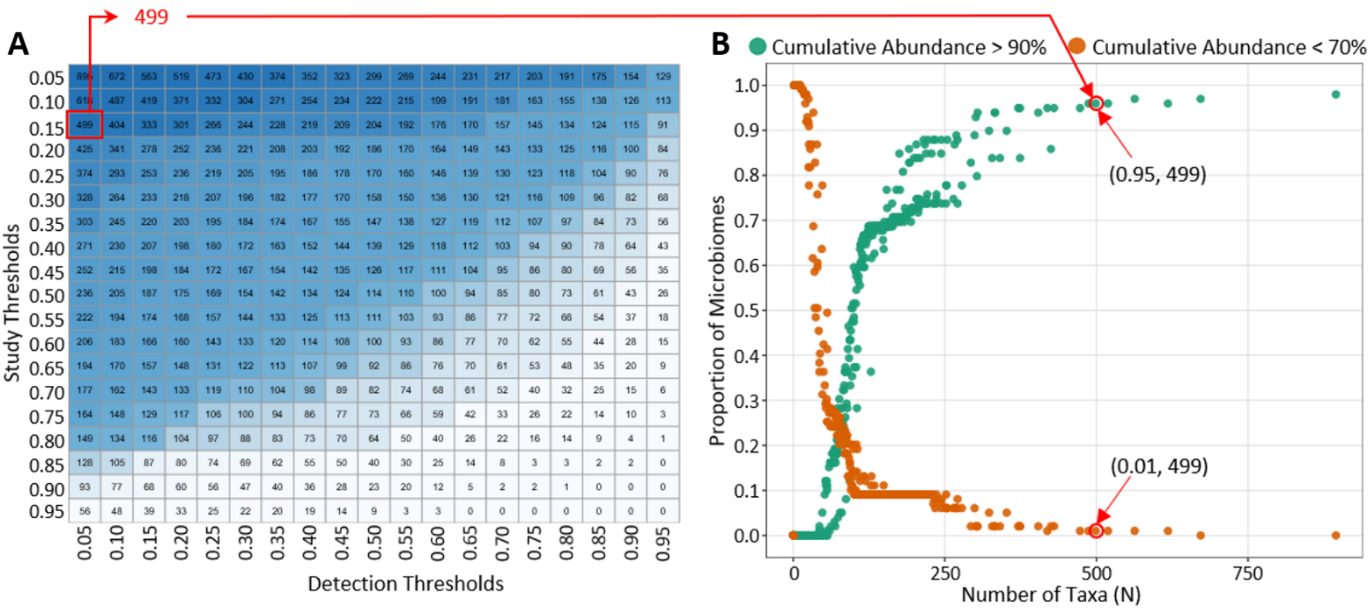


**Figure S1. Threshold optimization for identifying non-sparse consensus taxa in the salivary subsite. A.** Heatmap showing the sizes of the different candidate taxa sets when the taxa are selected based on different thresholds of the percentage of study-cohorts in which they are detected (Study Threshold, denoted on the rows) and the percentage of samples within each study-cohorts in which they are detected (Detection Threshold, denoted on columns). **B.** Cumulative representation of candidate salivary taxon sets (denoted in the cells of the heatmap above) across all threshold combinations. The x-axis denotes the sizes of the taxa sets. For each candidate taxa set combination, we plot two values in the scatterplot (denoted in the y-axis) denotes as points in green and red colours. The point in green colour shows the proportion of salivary study-cohorts in which the retained taxa combination cumulatively represented ≥ 90% and the point in orange colour shows the proportion of salivary study-cohorts where the cumulative abundance of the same taxa set was < 70%. The objective was to identify a taxa set with the minimum size that had the highest value of the green point and the lowest value corresponding to the orange point. The selected taxa set was of 499 taxa, corresponding to taxa detected in at least 5% of samples within at least 15% of salivary study-cohorts. This taxon set cumulatively represented ≥90% of the salivary microbial abundance in 95% of study-cohorts (N=94), while representing <70% abundance in only 1% of study-cohorts (N=1)**.**

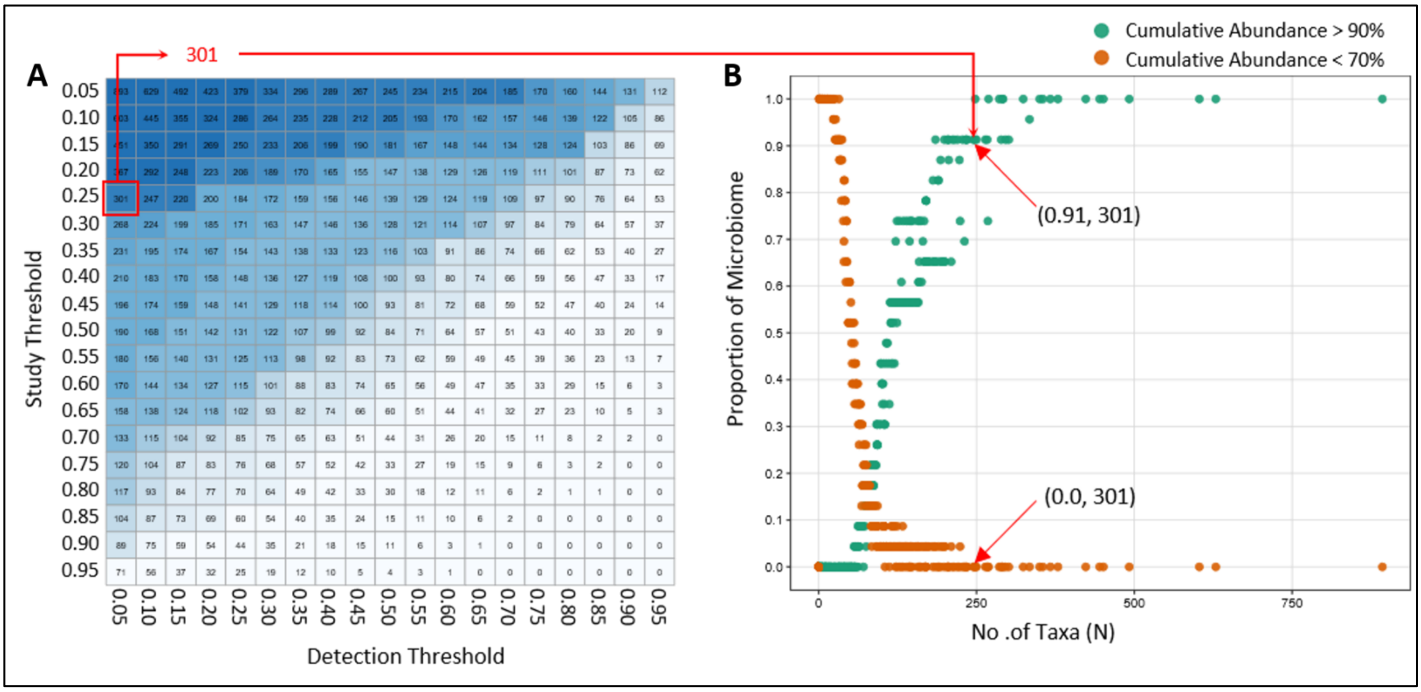


**Figure S2. Threshold optimization for identifying non-sparse consensus taxa in the supragingival subsite. A.** Heatmap showing the sizes of the different candidate taxa sets when the taxa are selected based on different thresholds of the percentage of study-cohorts in which they are detected (Study Threshold, denoted on the rows) and the percentage of samples within each study-cohorts in they are detected (Detection Threshold, denoted on columns). **B.** Cumulative representation of candidate salivary taxon sets (denoted in the cells of the heatmap above) across all threshold combinations. The x-axis denotes the sizes of the taxa sets. For each candidate taxa set combination, we plot two values in the scatterplot (denoted in the y-axis) denotes as points in green and red colours. The point in green colour shows the proportion of salivary study-cohorts in which the retained taxa combination cumulatively represented ≥ 90% and the point in orange colour shows the proportion of salivary study-cohorts where the cumulative abundance of the same taxa set was < 70%. The objective was to identify a taxa set with the minimum size that had the highest value of the green point and the lowest value corresponding to the orange point. The selected set contained 301 taxa, corresponding to taxa detected in at least 5% of samples within at least 25% of supragingival study-cohorts. This taxon set cumulatively represented ≥90% of the mean supragingival microbial abundance in approximately 91.3% of study-cohorts (N = 21), and no study-cohort showed cumulative representation below 70%.

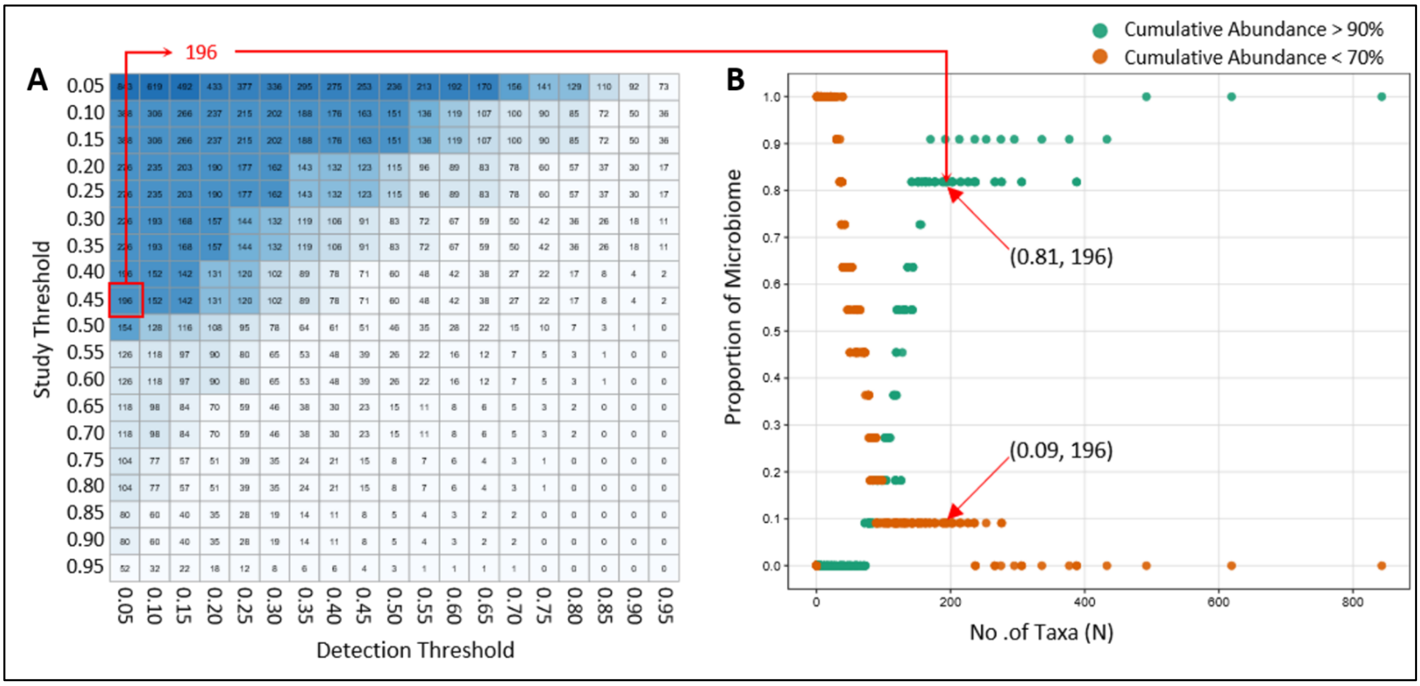


**Figure S3. Threshold optimization for identifying non-sparse consensus taxa in the subgingival subsite. A.** Heatmap showing the sizes of the different candidate taxa sets when the taxa are selected based on different thresholds of the percentage of study-cohorts in which they are detected (Study Threshold, denoted on the rows) and the percentage of samples within each study-cohorts in they are detected (Detection Threshold, denoted on columns). **B.** Cumulative representation of candidate salivary taxon sets (denoted in the cells of the heatmap above) across all threshold combinations. The x-axis denotes the sizes of the taxa sets. For each candidate taxa set combination, we plot two values in the scatterplot (denoted in the y-axis) denotes as points in green and red colours. The point in green colour shows the proportion of salivary study-cohorts in which the retained taxa combination cumulatively represented ≥ 90% and the point in orange colour shows the proportion of salivary study-cohorts where the cumulative abundance of the same taxa set was < 70%. The objective was to identify a taxa set with the minimum size that had the highest value of the green point and the lowest value corresponding to the orange point. The selected contained 196 taxa, corresponding to taxa detected in at least 5% of samples within at least 45% of subgingival study-cohorts. This taxon set cumulatively represented ≥90% of the mean subgingival microbial abundance in approximately 81% of study-cohorts (N=9), while representing <70% abundance in 9% of study-cohorts (N=1), indicating weaker saturation relative to the other oral subsites.

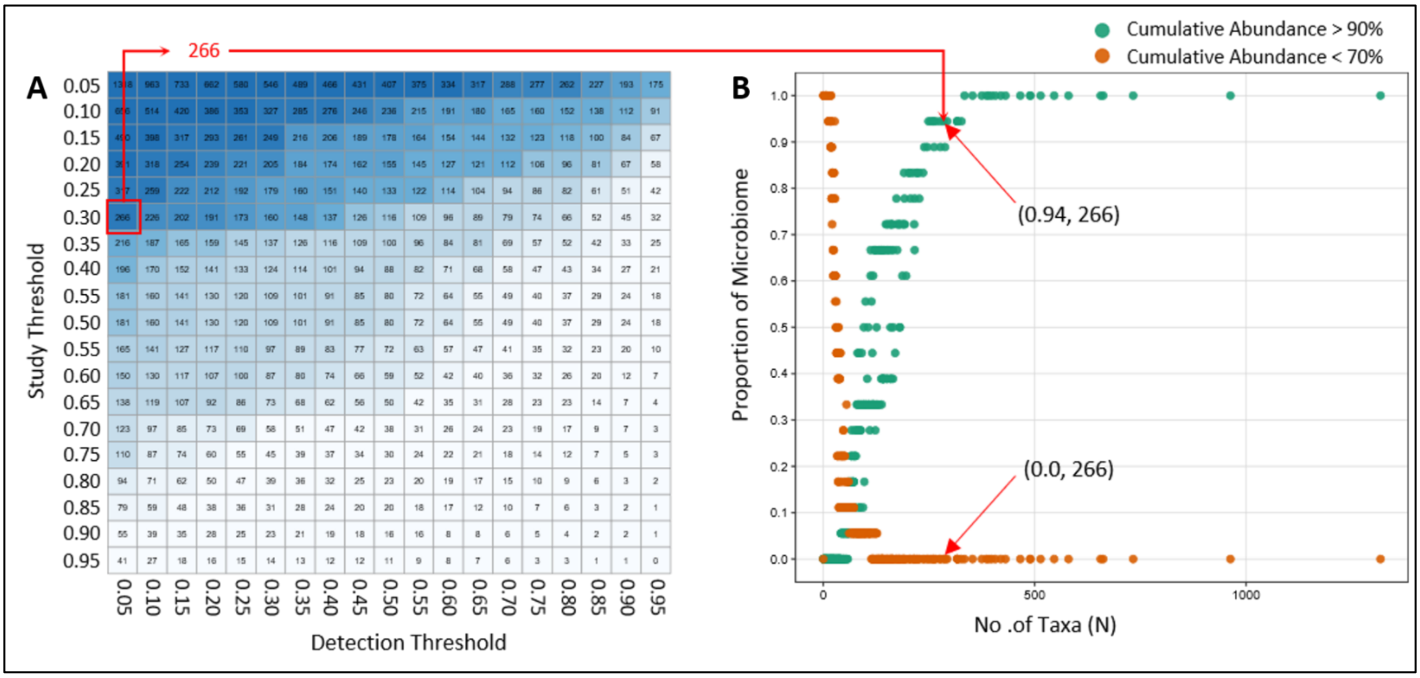


**Figure S4. Threshold optimization for identifying non-sparse consensus taxa in the Tongue-tonsil-related subsite. A.** Heatmap showing the sizes of the different candidate taxa sets when the taxa are selected based on different thresholds of the percentage of study-cohorts in which they are detected (Study Threshold, denoted on the rows) and the percentage of samples within each study-cohorts in they are detected (Detection Threshold, denoted on columns). **B.** Cumulative representation of candidate salivary taxon sets (denoted in the cells of the heatmap above) across all threshold combinations. The x-axis denotes the sizes of the taxa sets. For each candidate taxa set combination, we plot two values in the scatterplot (denoted in the y-axis) denotes as points in green and red colours. The point in green colour shows the proportion of salivary study-cohorts in which the retained taxa combination cumulatively represented ≥ 90% and the point in orange colour shows the proportion of salivary study-cohorts where the cumulative abundance of the same taxa set was < 70%. The objective was to identify a taxa set with the minimum size that had the highest value of the green point and the lowest value corresponding to the orange point. The selected set retained 266 taxa, corresponding to taxa detected in at least 5% of samples within at least 30% of tongue–tonsil study-cohorts. This taxon set cumulatively represented ≥90% of the mean tongue-tonsil microbial abundance in approximately 94% of study-cohorts (N=17), while representing <70% abundance in none of the study-cohorts.

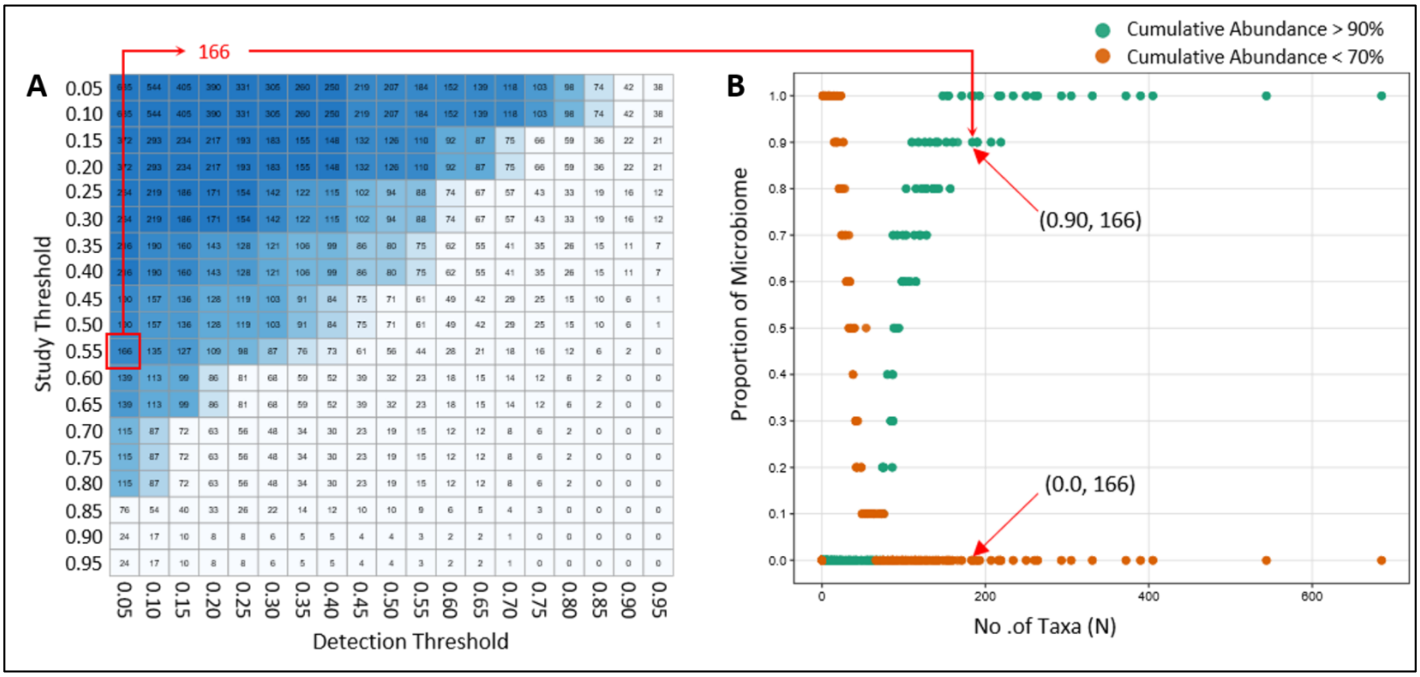


**Figure S5. Threshold optimization for identifying non-sparse consensus taxa in the buccal–palate-other surface subsite. A.** Heatmap showing the sizes of the different candidate taxa sets when the taxa are selected based on different thresholds of the percentage of study-cohorts in which they are detected (Study Threshold, denoted on the rows) and the percentage of samples within each study-cohorts in they are detected (Detection Threshold, denoted on columns). **B.** Cumulative representation of candidate salivary taxon sets (denoted in the cells of the heatmap above) across all threshold combinations. The x-axis denotes the sizes of the taxa sets. For each candidate taxa set combination, we plot two values in the scatterplot (denoted in the y-axis) denotes as points in green and red colours. The point in green colour shows the proportion of salivary study-cohorts in which the retained taxa combination cumulatively represented ≥ 90% and the point in orange colour shows the proportion of salivary study-cohorts where the cumulative abundance of the same taxa set was < 70%. The objective was to identify a taxa set with the minimum size that had the highest value of the green point and the lowest value corresponding to the orange point. The selected threshold retained 166 taxa, corresponding to taxa detected in at least 5% of samples within at least 55% of buccal–palate study-cohorts. This taxon set cumulatively represented ≥90% of the mean buccal–palate microbial abundance in approximately 90% of study-cohorts, while representing <70% abundance in none of the study-cohorts.

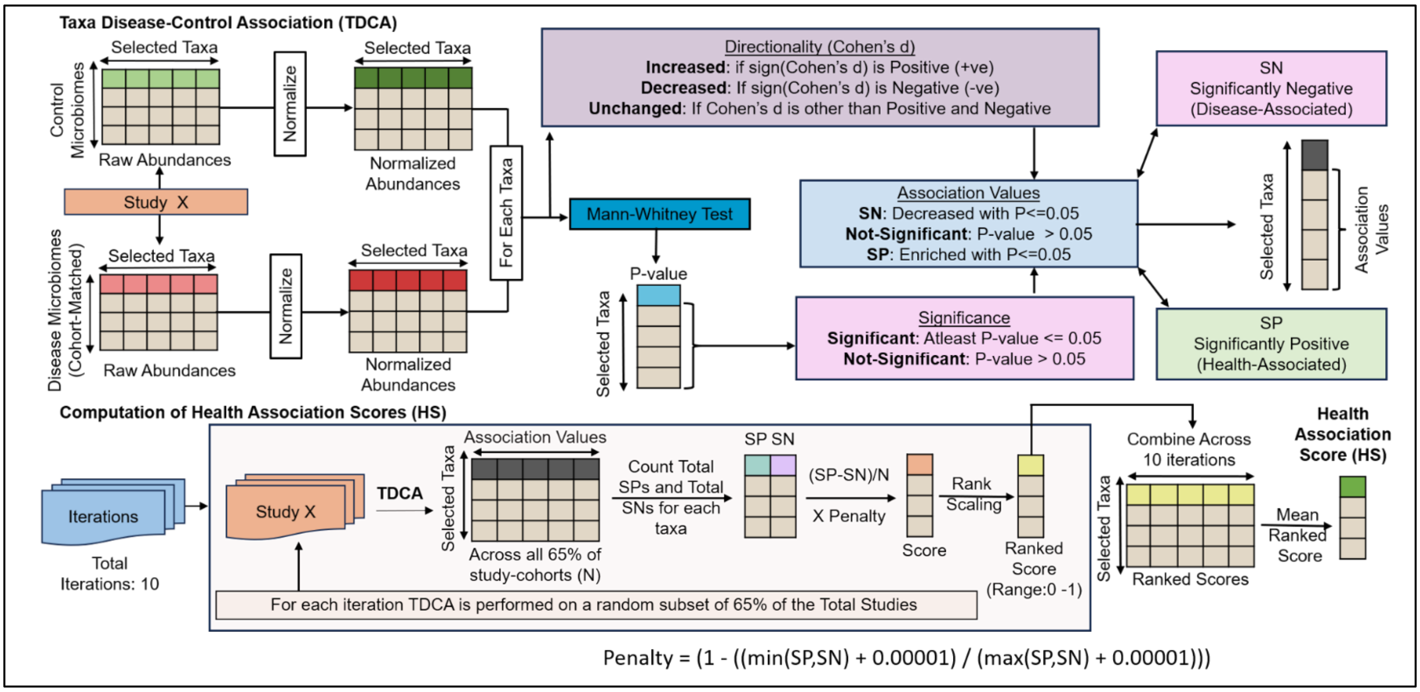


**Figure S6. Schematic overview of Health-Association Score (HS) computation.** It shows the modified HS computation schema adapted from the Taxa Disease-Control Association (TDCA) workflow in our previous study[1]. In summary, for each matched case-control study-cohort, taxon abundance profiles were first normalized within samples. For each taxon, abundance differences between control and disease samples were tested using the Mann–Whitney test, and the direction of association was determined using Cohen's *d*. Taxa with positive Cohen's *d* and enrichment (at least with P≤0.05) in control samples were classified as positive associations (SP; health-associated), whereas taxa with negative Cohen's *d* and depletion or enrichment in disease (at least with P≤0.05) in disease samples were classified as significantly associations (SN; disease-associated). Taxa without sufficient statistical evidence were classified as non-significant. To compute the Health Association Score, the scoring procedure was repeated across the study-cohorts for 10 iterations. In each iteration, a random subset of 65% of matched case-control study-cohorts was selected. For each taxon, the total number of SP and SN classifications was counted across the selected cohorts. An iteration-specific score was then calculated as (SP-SN)/N multiplied by an imbalance penalty, where *N* represents the number of selected study-cohorts. The penalty [Penalty = (1 - ((min(SP,SN) + 0.00001) / (max(SP,SN) + 0.00001)))] reduced scores for taxa showing inconsistent bidirectional associations across cohorts. Iteration-specific scores were then rank-scaled to a normalized range of 0–1, and the final Health-Association Score was calculated by averaging the rank-scaled scores across iterations.

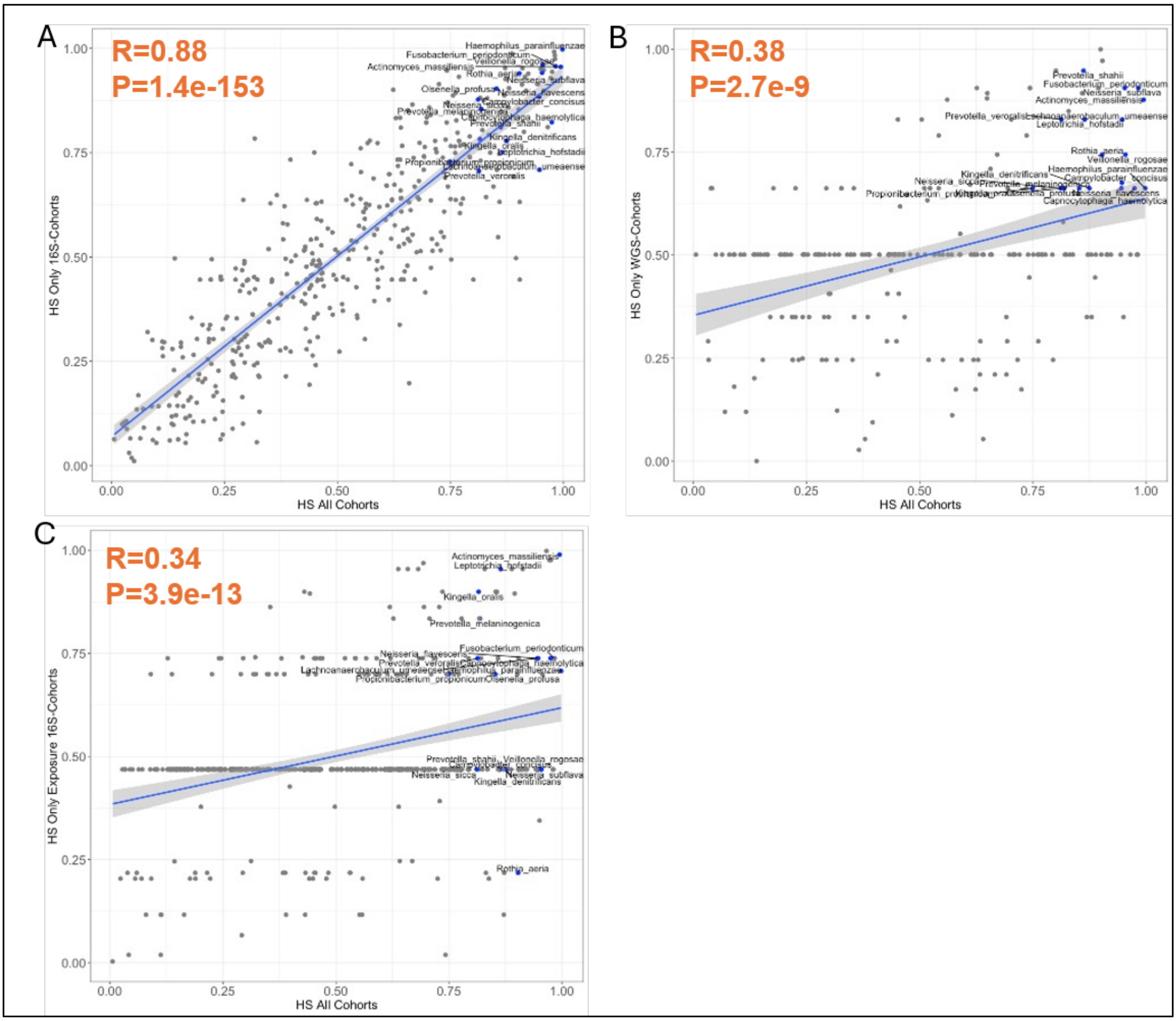


**Figure S7. Reproducibility of salivary Health-Association Scores (HS) across cohort-subsets using different profiling strategies (or sequencing types) as well as cohort-subsets investigating specific environmental exposure.** Scatter plots comparing Health-Association Scores (HS) computed using all salivary case-control cohorts (38) with HS computed from stratified cohort subsets. Each point represents one salivary taxon. Labelled taxa represent consistent health-associated taxa identified in the overall cohort analysis that also ranked within the top 35% across all stratified HS analyses. **A.** Comparison of overall HS computed from all salivary cohorts (on x-axis) with HS computed using only 16S sequencing cohorts (on y-axis), showing strong concordance (Spearman Rho=0.88, P=1.4e-153) between the overall and 16S-derived HS rankings. **B.** Comparison of overall HS computed from all salivary cohorts (on x-axis) with HS computed using only WGS cohorts (on y-axis), showing a significant positive correlation (Spearman Rho=0.38, P=2.7e-9) between the overall and WGS-derived rankings. **C.** Comparison of overall HS computed from all salivary cohorts (on x-axis) with HS computed using exposure-associated 16S cohorts (on y-axis), showing significant positive concordance (Spearman Rho=0.34, P=3.9e-13) between the overall and exposure-associated health-association rankings. Taxa showing an HS ≥ 0.70 in either the overall or corresponding stratified analysis are highlighted in plots **A**-**C**.

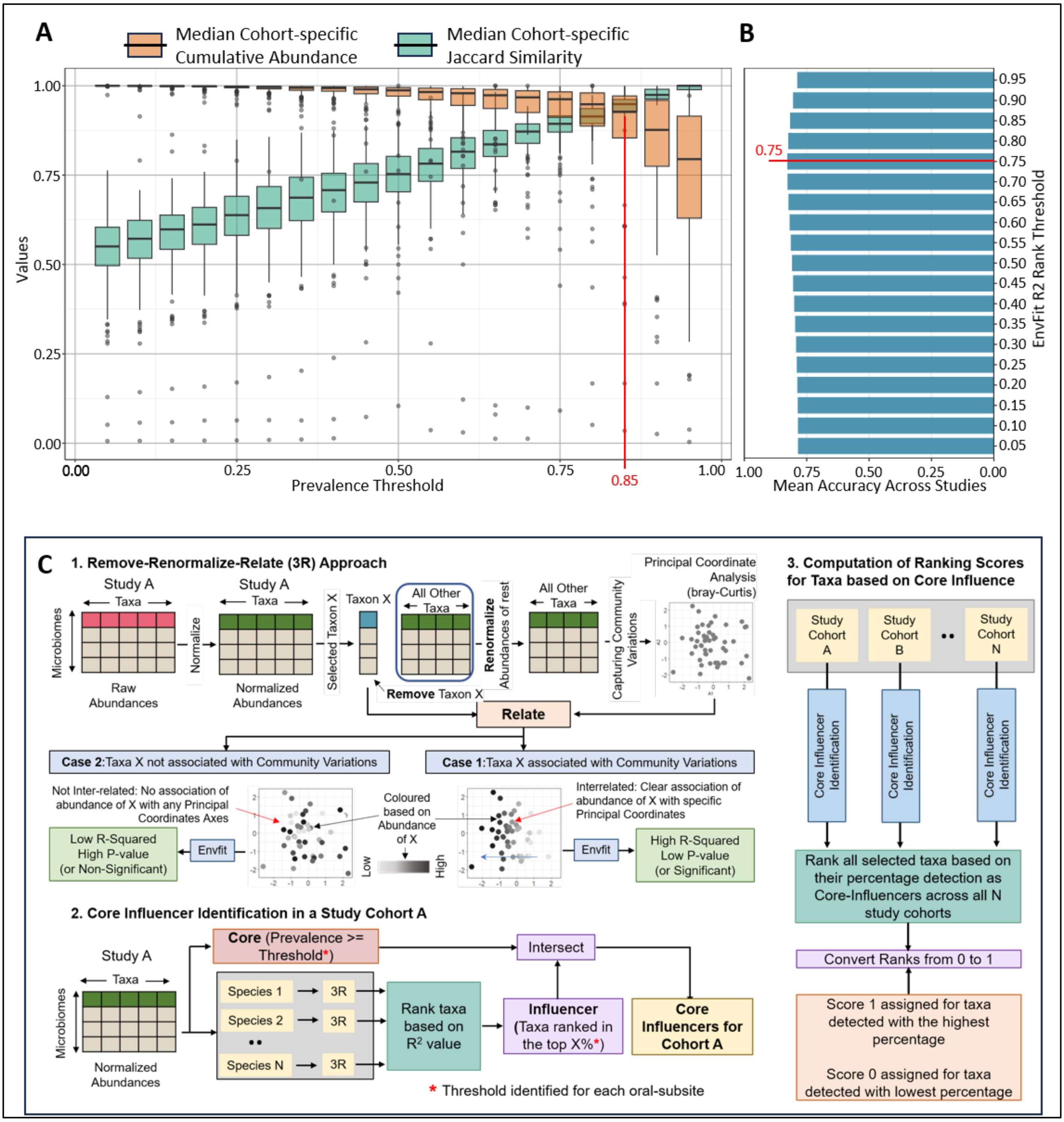


**Figure S8. Optimization and schematic description for the salivary Core-Association Score (CS) computation framework. A.** Selection of the prevalence threshold for defining candidate core taxa in salivary control cohorts. For each prevalence cutoff along the x-axis, taxa detected above that cutoff within a study-cohort were selected, and two properties were then calculated across that cohort's samples: the median Jaccard similarity across all sample pairs (using this taxa subset) and the median cumulative abundance of this taxa subset across samples. These two properties were computed for every study-cohort, and their distributions across cohorts are shown as paired boxplots at each cutoff: light-brown for median cumulative abundance and green for median Jaccard similarity. For low prevalence thresholds, while the identified core taxa would always account for a large proportion of the microbiome, there is going to be high variation in their detection rates across microbiomes. For stringent or high prevalence thresholds, the selected small taxa set would show minimal variation across samples but would also capture only a small proportion of the microbiome. An ideal core-taxa set should be both uniformly represented across samples (high Jaccard similarity) and account for a large proportion of overall microbiome composition (high cumulative abundance); the optimal cutoff is one that jointly maximizes both properties. Based on this trade-off across cutoff thresholds, a cutoff of 0.85 was selected. **B.** Selection of the community-association threshold for identifying taxa strongly linked to salivary community structure. Taxa were ranked by $R^2$, and a threshold was chosen to balance two failure modes: a threshold set too high would retain only taxa consistently associated with other members' abundances, but would miss taxa with significant community-composition associations that nonetheless have lower ranked $R^2$ (reduced sensitivity); a threshold set too low would misclassify taxa with non-significant associations as ecologically influential (increased false positives). The ideal threshold therefore maximizes accuracy, the proportion of correctly identified taxa with significant community-composition associations. Based on this evaluation, a cutoff of 0.75 was selected, as it maintained high mean accuracy across studies while retaining taxa with strong envfit-derived community-association signals. **C.** Schematic overview of Core-Association Score computation using the Remove-Renormalize-Relate (3R) framework adopted from our previous study[1]. Within each salivary control cohort, taxa were evaluated using the 3R framework. Each taxon was removed, the remaining community was renormalized, and Bray–Curtis PCoA was used to reconstruct community variation. The abundance of the removed taxon was then related to this ordination using envfit, and taxa with high, significant $R^2$ values were considered community-associated. Taxa passing both the optimized prevalence threshold and the community-association threshold were defined as core-influencer taxa within each cohort. Across cohorts, taxa were ranked by how often they were identified as core-influencers, and these recurrence values were rank-scaled from 0 to 1 to generate the final Core-Association Score.

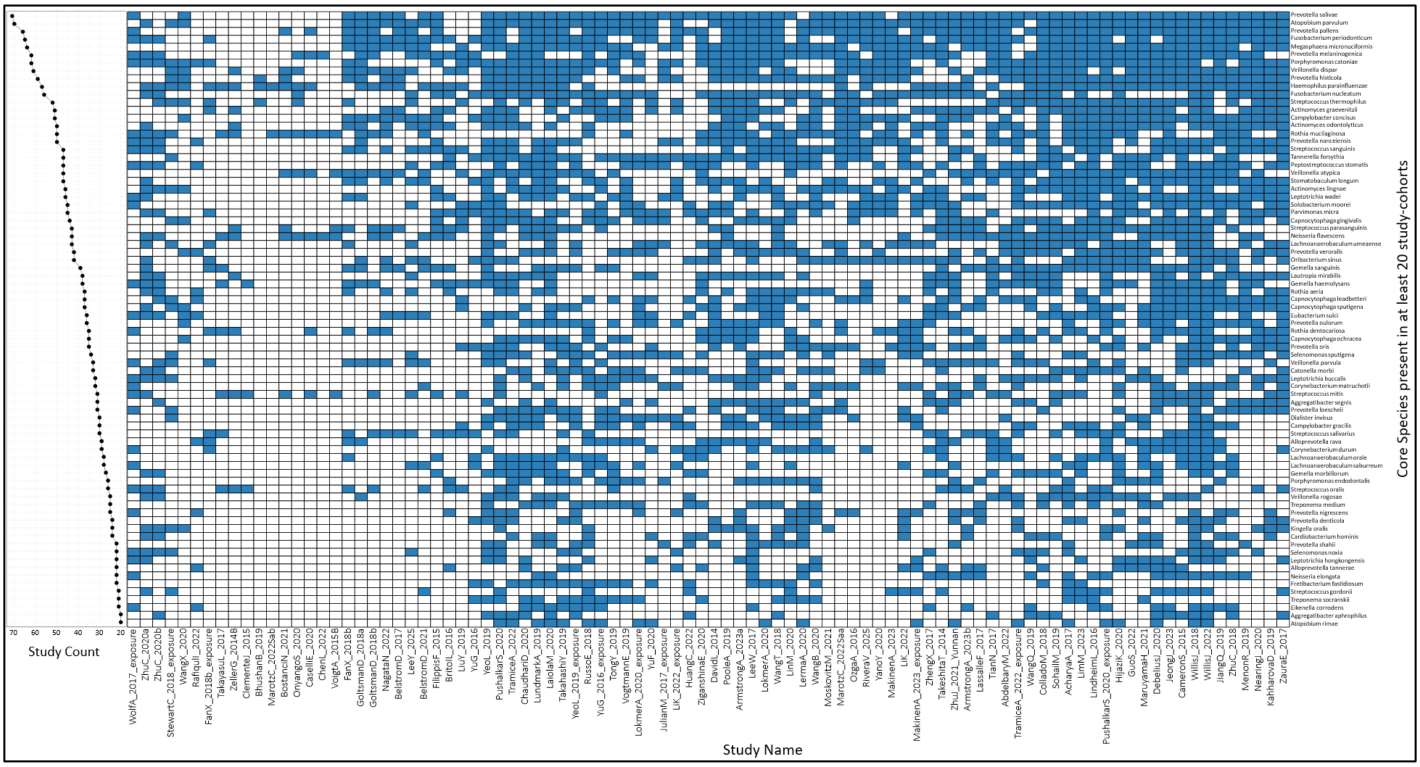


**Figure S9. Study-cohort-wise identification of salivary core-associated taxa.** Heatmap showing the identification of core-associated taxa across salivary control study-cohorts. Rows represent salivary taxa and columns represent individual control study-cohorts. Blue cells indicate that a taxon was classified as core-associated within that cohort after passing the optimized prevalence threshold of 0.85 and $R^2$ rank threshold of 0.75. Taxa are ordered according to their Core-Association Score, and the left-side line plot shows their recurrence across cohorts. Only taxa identified as core-associated in at least 20 study-cohorts out of 99 are shown.

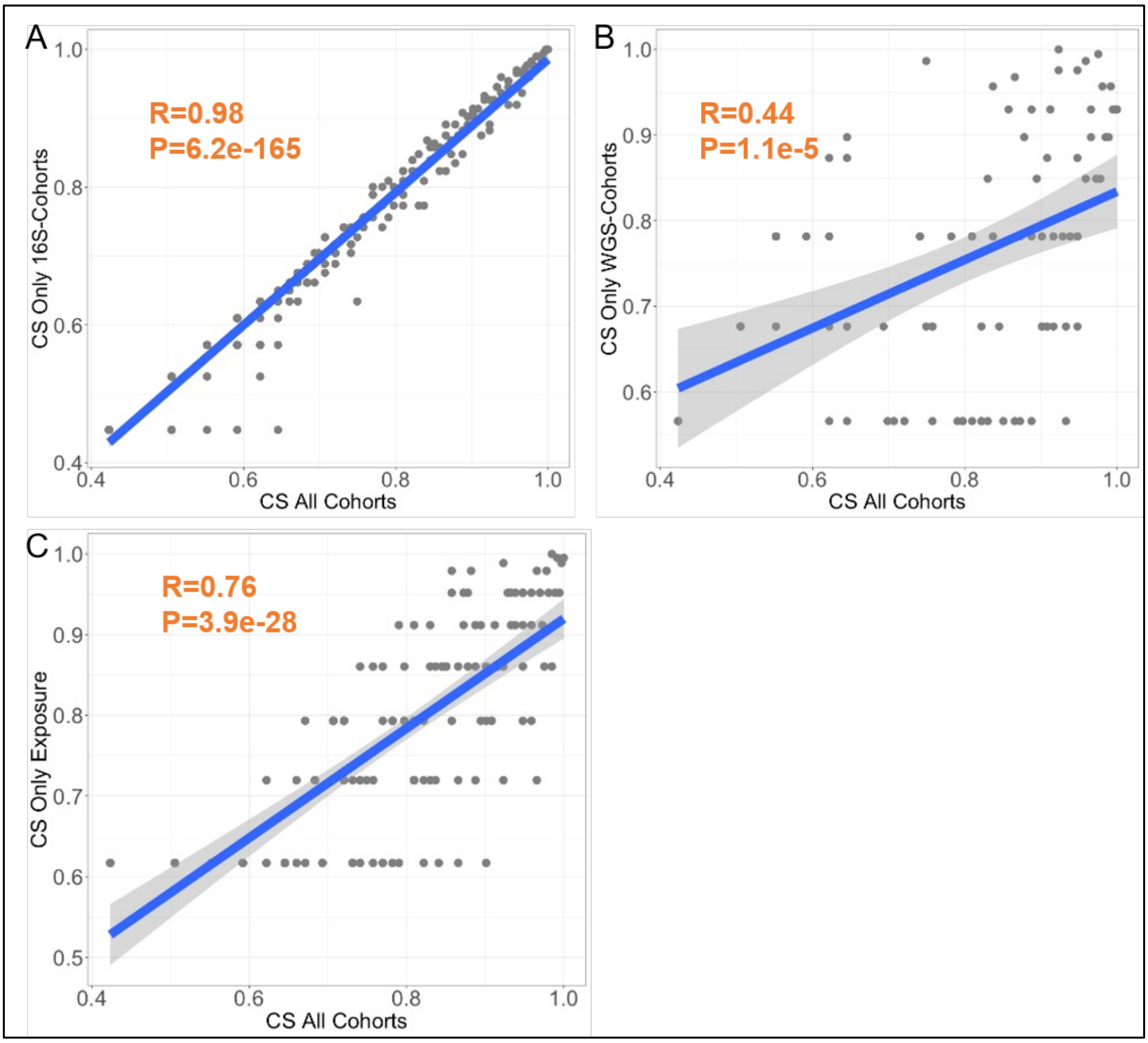


**Figure S10. Reproducibility of salivary Core-Association Scores (CS) across cohort-subsets using different profiling strategies (or sequencing types) as well as cohort-subsets investigating specific environmental exposure. A.** Scatterplot comparison the overall CS computed from all salivary cohorts (x-axis) with CS computed using only 16S sequencing cohorts (y-axis), showing strong concordance (Spearman Rho=0.98, P=6.2e-165) between the overall and 16S-derived core-association rankings. **B.** Similar comparison of overall CS computed from all salivary cohorts with CS computed using only WGS cohorts, showing a significant positive correlation (Spearman Rho=0.44, P=1.1e-5) between the overall and WGS-derived core-association rankings. **C.** Comparison of CS computed from all salivary cohorts with CS computed using exposure-associated 16S cohorts, showing significant positive concordance (Spearman Rho=0.76, P=3.9e-28) between the overall and exposure-associated core-association rankings.

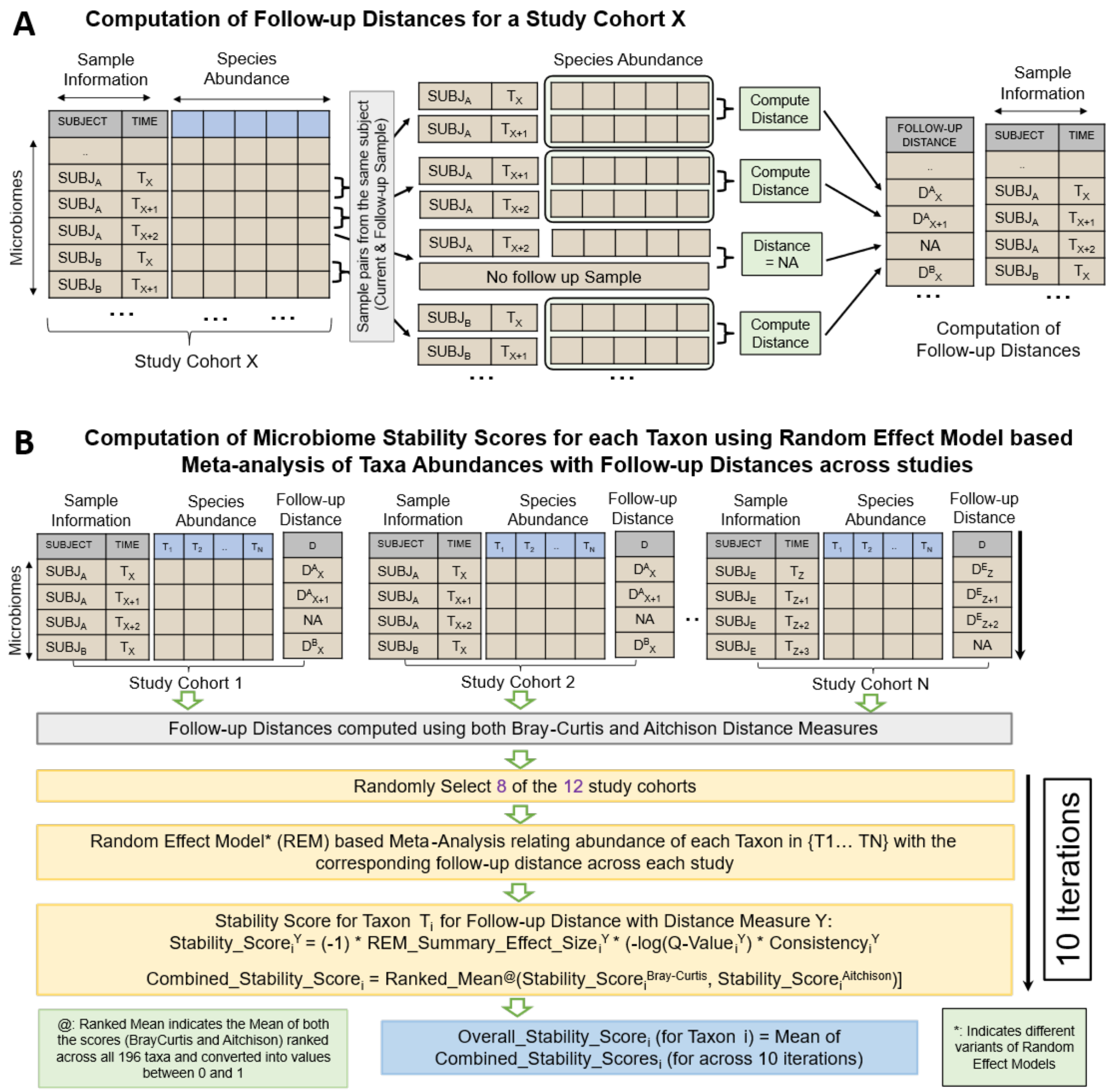


**Figure S11. Schematic overview of Stability-Association Score computation.** Schematic showing the workflow used to compute salivary Stability-Association Scores, adapted from Goel et al 2025[1]. For each longitudinal salivary cohort, samples from the same subject were paired across consecutive time points, and follow-up distance was computed between each sample and its immediate follow-up using Bray–Curtis and Aitchison distance measures. In each iteration, 8 of the 12 longitudinal cohorts were randomly selected, and taxon abundance was related to follow-up distance using random-effects meta-analysis. Taxa with stronger negative associations with follow-up distance were considered more stability-associated. Bray–Curtis- and Aitchison-based stability scores were rank-scaled and averaged, and the final Stability-Association Score was calculated as the mean combined score across 10 iterations. This Analysis was specific to salivary subsite only.

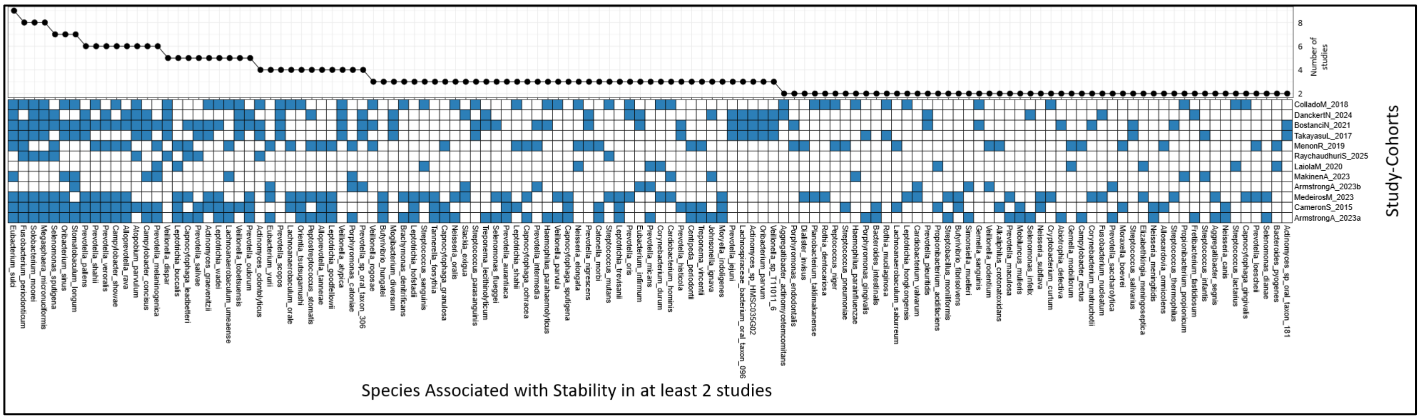


**Figure S12. Study-cohort-wise identification of stability-associated taxa in salivary microbiomes.** Heatmap showing the identification of stability-associated taxa across 12 longitudinal salivary study-cohorts. Rows represent longitudinal cohorts and columns represent salivary taxa. Blue coloured cells indicate that a taxon was classified as stability-associated within that cohort based on its negative association with follow-up microbiome distance. Taxa are ordered according to their Stability-Association Score, and the upper line plot shows the recurrence of stability-associated classification across cohorts. Only Taxa present in at least 2 study-cohorts are shown here.

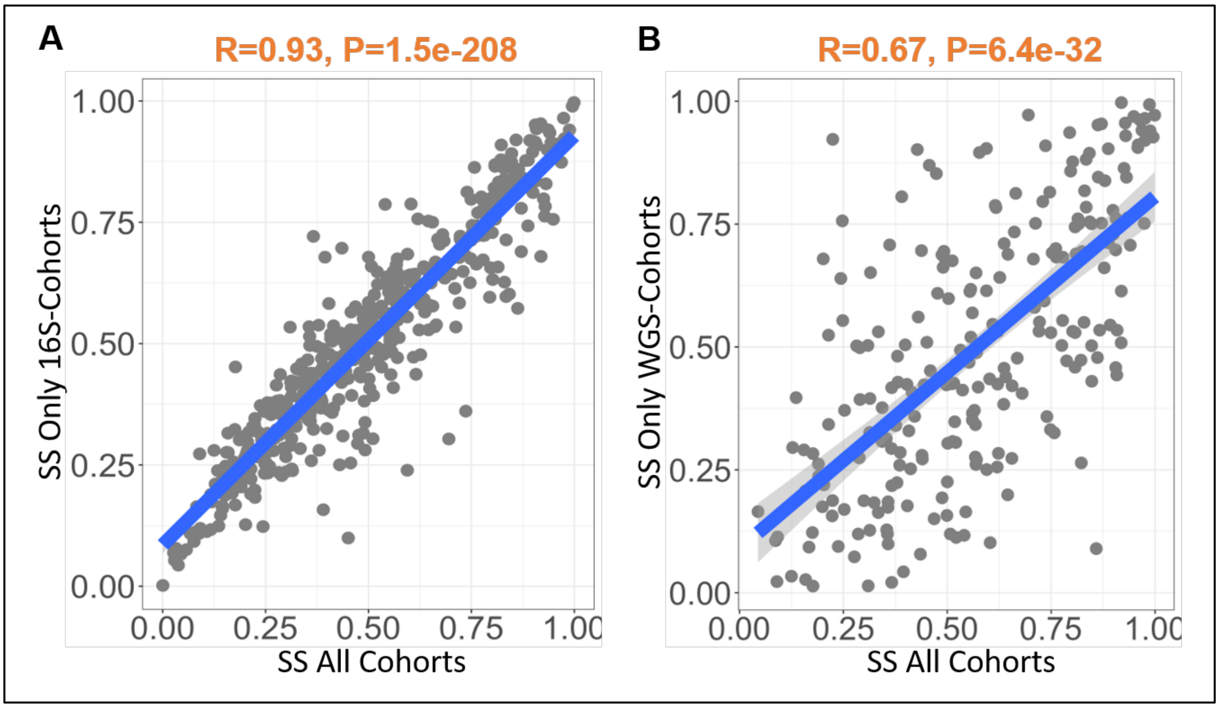


**Figure S13. Reproducibility of salivary Stability-Association Scores (SS) across cohort-subsets using different profiling strategies (or sequencing types).** Scatter plots comparing salivary Stability-Association Scores (SS) computed using all longitudinal salivary cohorts with SS computed from sequencing-specific longitudinal cohort subsets. Each point represents one salivary taxon. **A.** Comparison of SS computed from all longitudinal salivary cohorts with SS computed using only 16S sequencing cohorts, showing strong concordance (Spearman Rho = 0.93, P= 1.5e-208) between the overall and 16S-derived stability-association rankings. **B.** Comparison of SS computed from all longitudinal salivary cohorts with SS computed using only WGS cohorts, showing significant positive concordance (Spearman Rho = 0.67, P= 6.4e-32) between the overall and WGS-derived stability-association rankings.

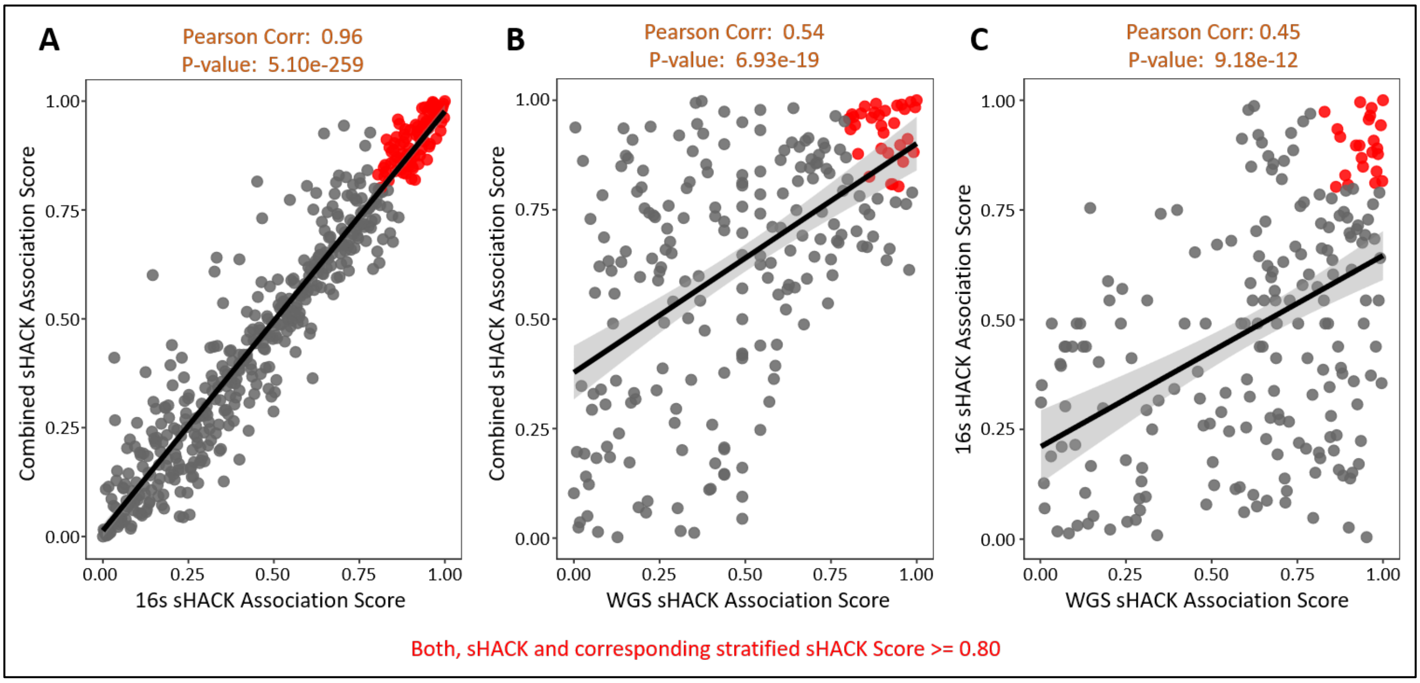


**Figure S14. Reproducibility of salivary Health-Associated Core Keystone (sHACK) scores across sequencing-type-specific cohort subsets.** Scatter plots comparing salivary Health-Associated Core Keystone scores (sHACK) computed using all salivary cohorts (by integrating Core-Associated score, Health-Associated Score and Stability-Associated Score - **Figure 3A**) with sHACK scores computed from sequencing-specific cohort subsets. Each point represents one salivary taxon and only taxa with both the scores >= 0.80 are highlighted in red. **A.** Comparison of sHACK scores computed from all salivary cohorts with sHACK scores computed using only 16S sequencing cohorts, showing strong concordance (Spearman Rho=0.96, P=5.1e-259) between the overall and 16S-derived sHACK rankings. **B.** Comparison of sHACK scores computed from all salivary cohorts with sHACK scores computed using only WGS cohorts, showing significant positive concordance (Spearman Rho=0.54, P=6.9e-19) between the overall and WGS-derived sHACK rankings. **C.** Direct comparison of sHACK scores computed from 16S cohorts and WGS cohorts, showing significant concordance (Spearman Rho=0.45, P=9.2e-12) between sequencing-specific sHACK rankings.

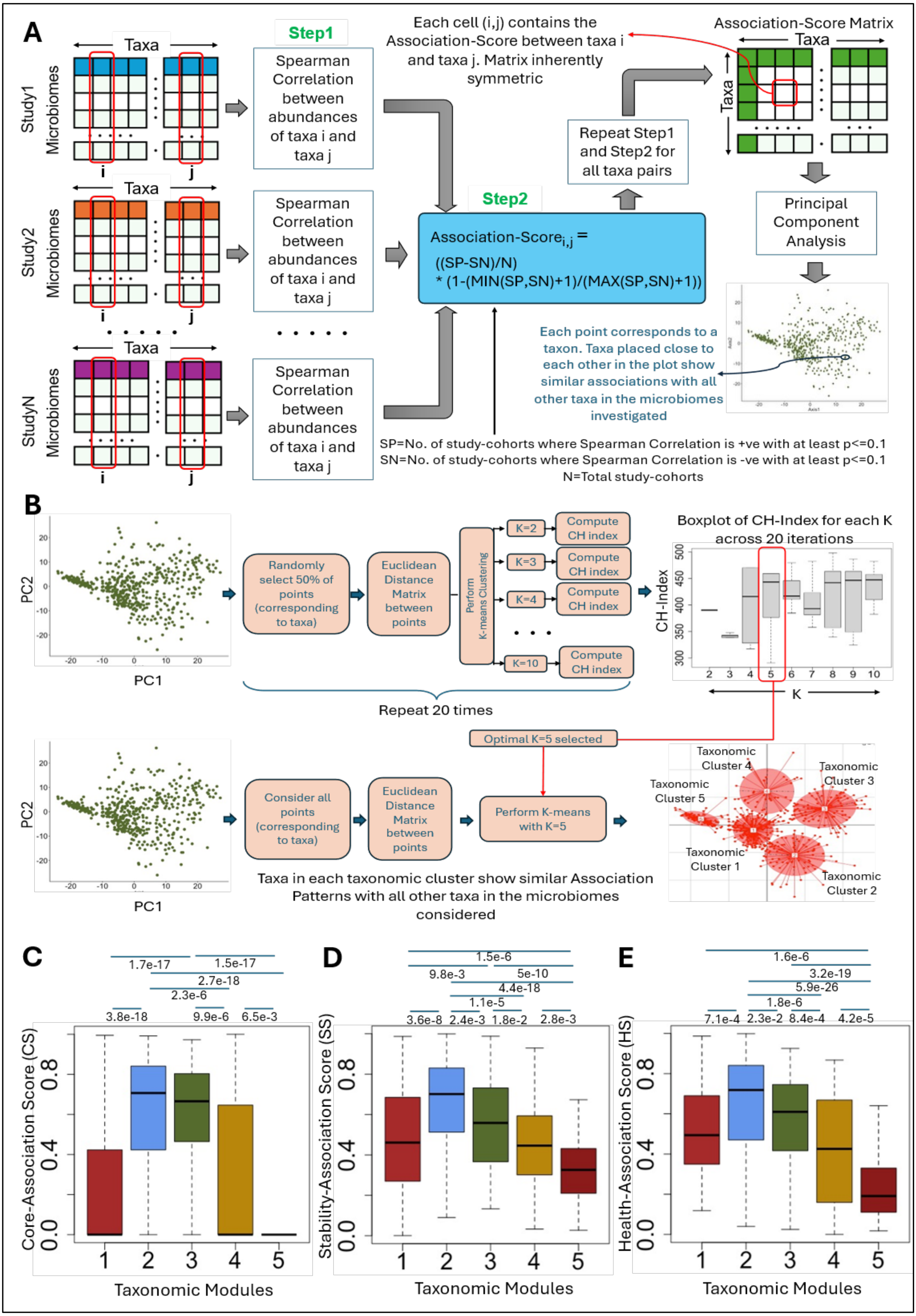
A
Step1
Taxa
Study1 Microbiomes
Study2 Microbiomes
StudyN Microbiomes
i
j
Spearman Correlation between abundances of taxa i and taxa j
Step2
Association-Score$_{i,j}$ = ((SP-SN)/N) * (1-(MIN(SP,SN)+1)/(MAX(SP,SN)+1))
Each cell (i,j) contains the Association-Score between taxa i and taxa j. Matrix inherently symmetric
Repeat Step1 and Step2 for all taxa pairs
Association-Score Matrix
Principal Component Analysis
Each point corresponds to a taxon. Taxa placed close to each other in the plot show similar associations with all other taxa in the microbiomes investigated
SP=No. of study-cohorts where Spearman Correlation is +ve with at least p<=0.1
SN=No. of study-cohorts where Spearman Correlation is -ve with at least p<=0.1
N=Total study-cohorts
B
PC1
PC2
Randomly select 50% of points (corresponding to taxa)
Euclidean Distance Matrix between points
Perform K-means Clustering
K=2
K=3
K=4
K=10
Compute CH index
Repeat 20 times
Boxplot of CH-Index for each K across 20 iterations
CH-Index
K
Optimal K=5 selected
Consider all points (corresponding to taxa)
Perform K-means with K=5
Taxonomic Cluster 1
Taxonomic Cluster 2
Taxonomic Cluster 3
Taxonomic Cluster 4
Taxonomic Cluster 5
Taxa in each taxonomic cluster show similar Association Patterns with all other taxa in the microbiomes considered
C
1.7e-17
1.5e-17
2.7e-18
2.3e-6
3.8e-18
9.9e-6
6.5e-3
Core-Association Score (CS)
Taxonomic Modules
D
1.5e-6
9.8e-3
5e-10
4.4e-18
1.1e-5
3.6e-8
2.4e-3
1.8e-2
2.8e-3
Stability-Association Score (SS)
Taxonomic Modules
E
1.6e-6
3.2e-19
5.9e-26
1.8e-6
7.1e-4
2.3e-2
8.4e-4
4.2e-5
Health-Association Score (HS)
Taxonomic Modules

**Figure S15. Identification and characterization of salivary taxonomic modules based on community-wide association patterns. A.** Schematic overview of the taxon–taxon Association Score framework used to construct salivary taxa association maps. For each pair of taxa, positive and negative association signals were summarized across study-cohorts of non-diseased salivary microbiomes (N = 15,127), yielding an Association-Score based on the consistency of association directionality between the two taxa, penalized for variability in that pattern across cohorts (**Text S3**). These pairwise scores were then used to position all taxa in a two-dimensional PCA space, where taxa located closer together show more similar association patterns with the rest of the salivary community. **B.** Selection of the optimal number of taxonomic modules using k-means clustering. Here, we performed 10 iterations. In each iteration, a random 50% subset of taxa was selected. Euclidean distances between each taxa pair in these selected taxa subset in the PCA space were computed, and k-means clustering was performed selecting different k values ranging from 2 to 10. The efficiency of clustering for each k value across the iterations was evaluated using Calinski–Harabasz (CH) indices. The distribution of CH indices across the 10 iterations were evaluated by examining their variations in boxplot. The minimum value of k that resulted in the highest median CH index with the least variation was selected. Here, k = 5 was selected as the optimal value. A final k-means clustering considering all taxa with k = 5 was then applied to all taxa to define the five discovery-cohort salivary modules. **C-E.** Distribution of Core-Association Score (CS) (**C**), Health-Association Score (HS) (**D**), and Stability-Association Score (SS) (**E**) across the five salivary taxonomic modules. Boxplots show score distributions for taxa within each module, highlighting module-level differences in ecological influence, health-association, and longitudinal stability-association.

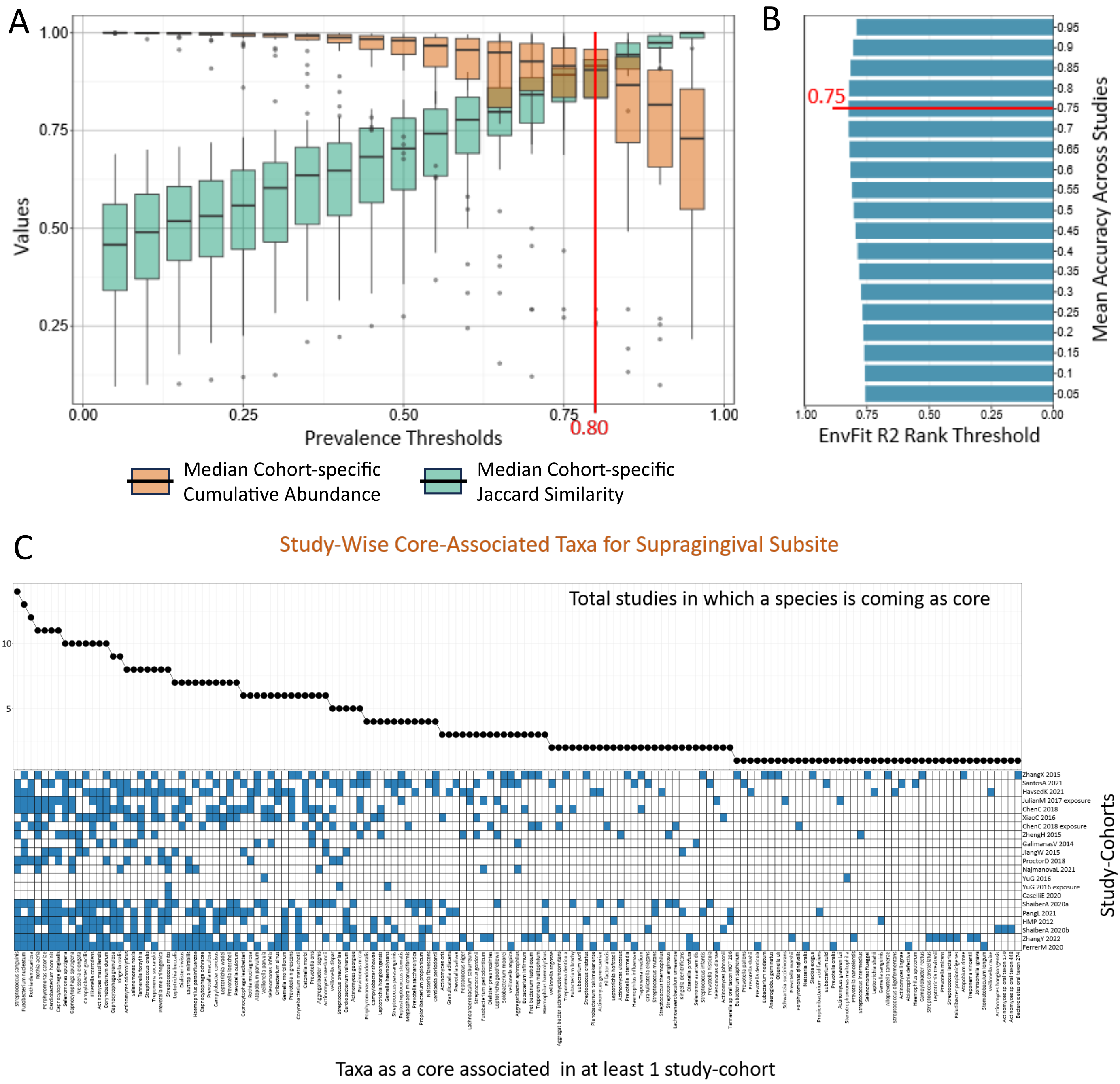


**Figure S16. Threshold optimization of the 3R framework and cohort-specific identification of Core-Associated taxa in the supragingival microbiomes. A. Selection of the prevalence threshold for defining candidate core taxa in supragingival control cohorts.** For each prevalence cutoff along the x-axis, taxa detected above that cutoff within a study-cohort were selected, and two properties were then calculated across that cohort's samples: the median Jaccard similarity across all sample pairs (using this taxa subset) and the median cumulative abundance of this taxa subset across samples. These two properties were computed for every study-cohort, and their distributions across cohorts are shown as paired boxplots at each cutoff: light-brown for median cumulative abundance and green for median

Jaccard similarity. For low prevalence thresholds, while the identified core taxa would always account for a large proportion of the microbiome, there is going to be high variation in their detection rates across microbiomes. For stringent or high prevalence thresholds, the selected small taxa set would show minimal variation across samples but would also capture only a small proportion of the microbiome. An ideal core-taxa set should be both uniformly represented across samples (high Jaccard similarity) and account for a large proportion of overall microbiome composition (high cumulative abundance); the optimal cutoff is one that jointly maximizes both properties. Candidate prevalence thresholds were evaluated to identify the cutoff of >= 0.80, that retained consistently detected taxa while maintaining broad community representation across supragingival control cohorts. **B. Selection of the community-association threshold for identifying taxa strongly linked to supragingival community structure.** Taxa were ranked by $R^2$, and a threshold was chosen to balance two failure modes: a threshold set too high would retain only taxa consistently associated with other members' abundances, but would miss taxa with significant community-composition associations that nonetheless have lower ranked $R^2$ (reduced sensitivity); a threshold set too low would misclassify taxa with non-significant associations as ecologically influential (increased false positives). The ideal threshold therefore maximizes accuracy, the proportion of correctly identified taxa with significant community-composition associations. The optimized $R^2$ rank threshold of 0.75 was selected as, for this threshold, we observed the maximum accuracy for identifying taxa whose variation was significantly associated with variation of other members of microbial communities. Finally, the cohort-specific lists of core-associated taxa were generated using a prevalence cutoff of >= 0.80 and $R^2$ rank threshold >= 0.75 using 3R framework. **C. Heatmap showing the identification of core-associated taxa across supragingival control study-cohorts.** Rows represent supragingival taxa and columns represent individual control cohorts. Blue cells indicate that a taxon was classified as core-associated within that cohort after passing the optimized prevalence threshold of 0.80 and rank-scaled $R^2$ threshold of 0.75. Taxa are ordered according to their Core Association Score, and the top line plot shows their recurrence across cohorts. Only taxa identified as core-associated in at least one study-cohort are shown.

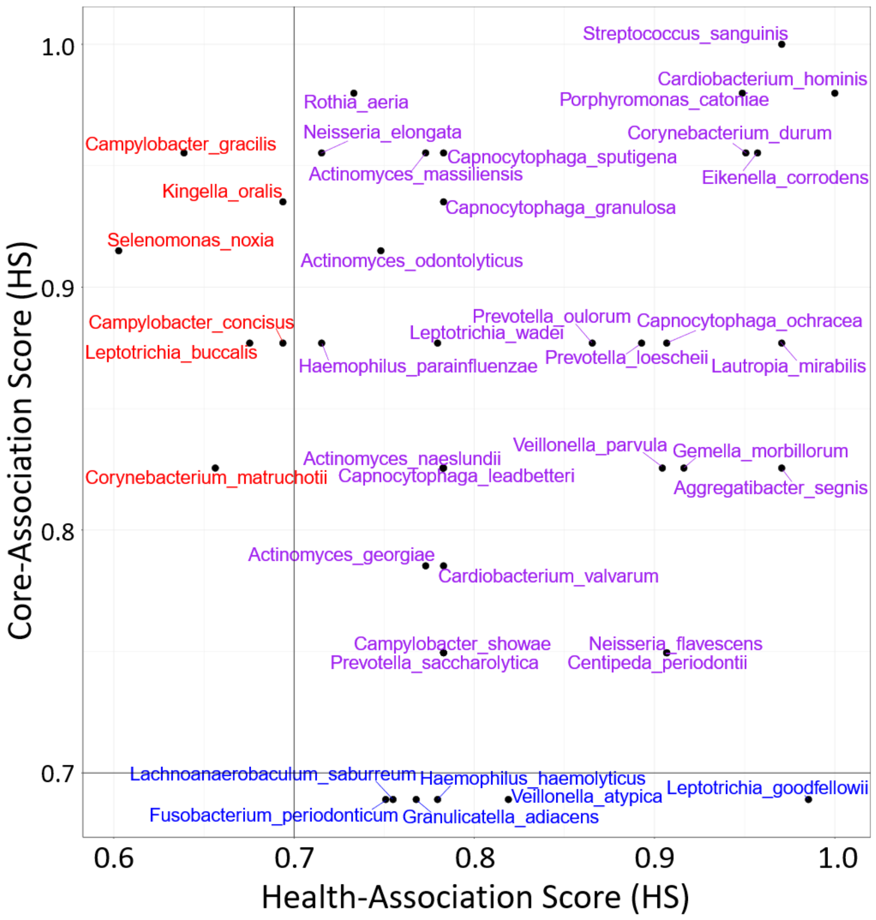


**Figure S17. Scatter plot comparing Core-Association Score (CS) and Health-Association Score (HS) for supragingival taxa**. Only taxa with CS and HS values >=0.60 and are shown. A score threshold of 0.70 for each score was used to stratify taxa into quadrants and identify taxa that were both recurrently core-associated across control cohorts and consistently health-associated across matched case-control cohorts (taxa shown in purple colour).

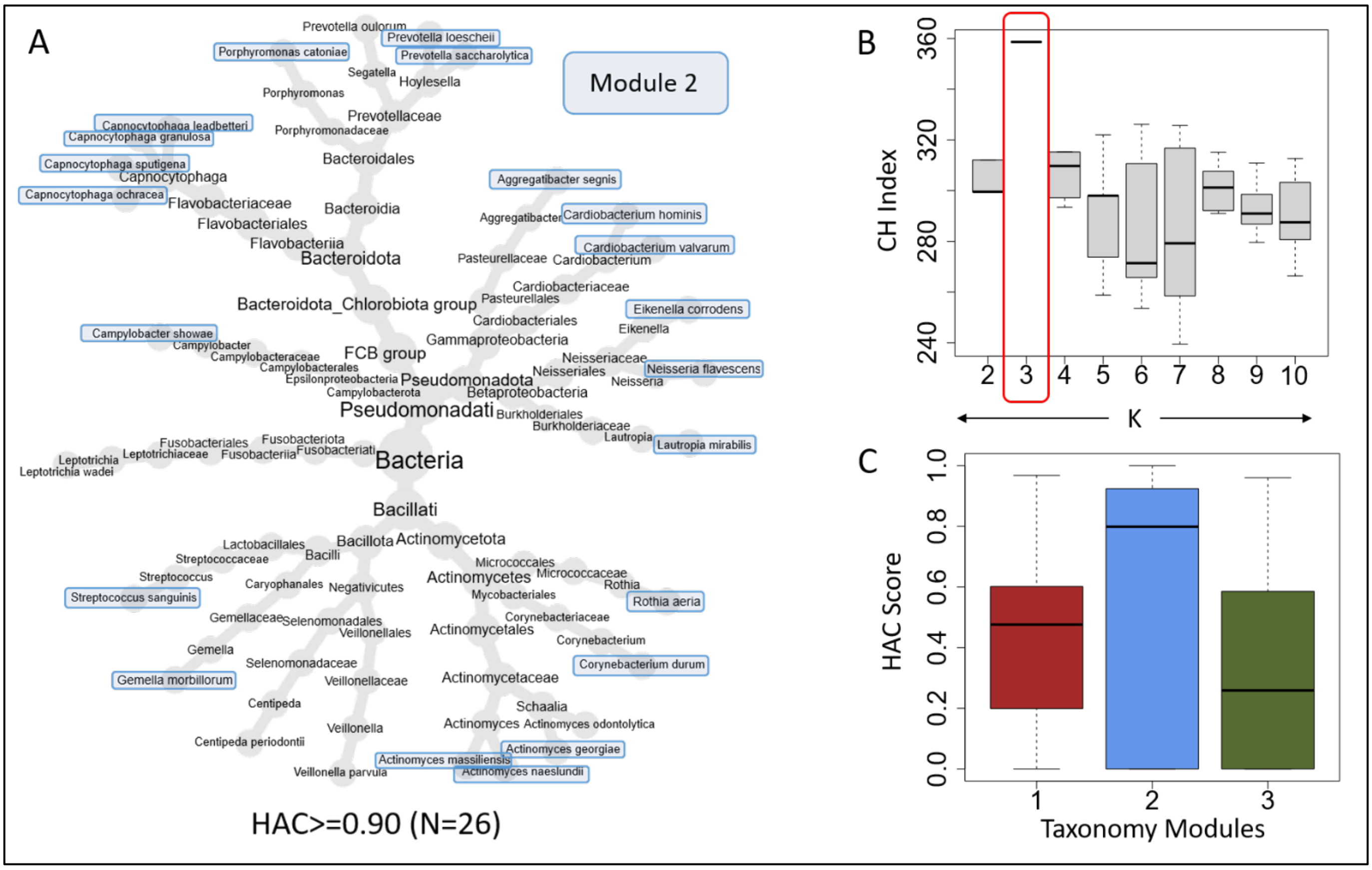


**Figure S18. Identification of the 26 member high-HAC-taxa set (HAC >= 0.90) in the supragingival microbiomes and the high-HAC-enriched microbiome community (supragingival-module-2). A.** Circular phylogenetic tree showing the top 26 high-HAC taxa (HAC >= 0.90) in the supragingival microbiomes, showing their distribution across multiple bacterial lineages. Taxa belonging to taxonomic module 2 (described below, the identification of which is described next) are highlighted in blue boxes. **B.** Selection of the optimal k (=3) used for the identification of supragingival taxonomic modules using iterative k-means clustering analysis. Pairwise association profiles among supragingival consensus taxa were projected into a two-dimensional association space, and a bootstrap based iterative evaluation for various values of k using CH-indices to identify the smallest value of k that gave the best median CH index values across iterations (see **Text S5** and **Figure S15** for methodological details). This analysis selected 3 supragingival taxonomic modules. **C.** Distribution of HAC scores across the three supragingival modules. Module 2 showed noticeably higher HAC values, indicating enrichment of taxa with concordant Health-Association and Core-Association Scores.

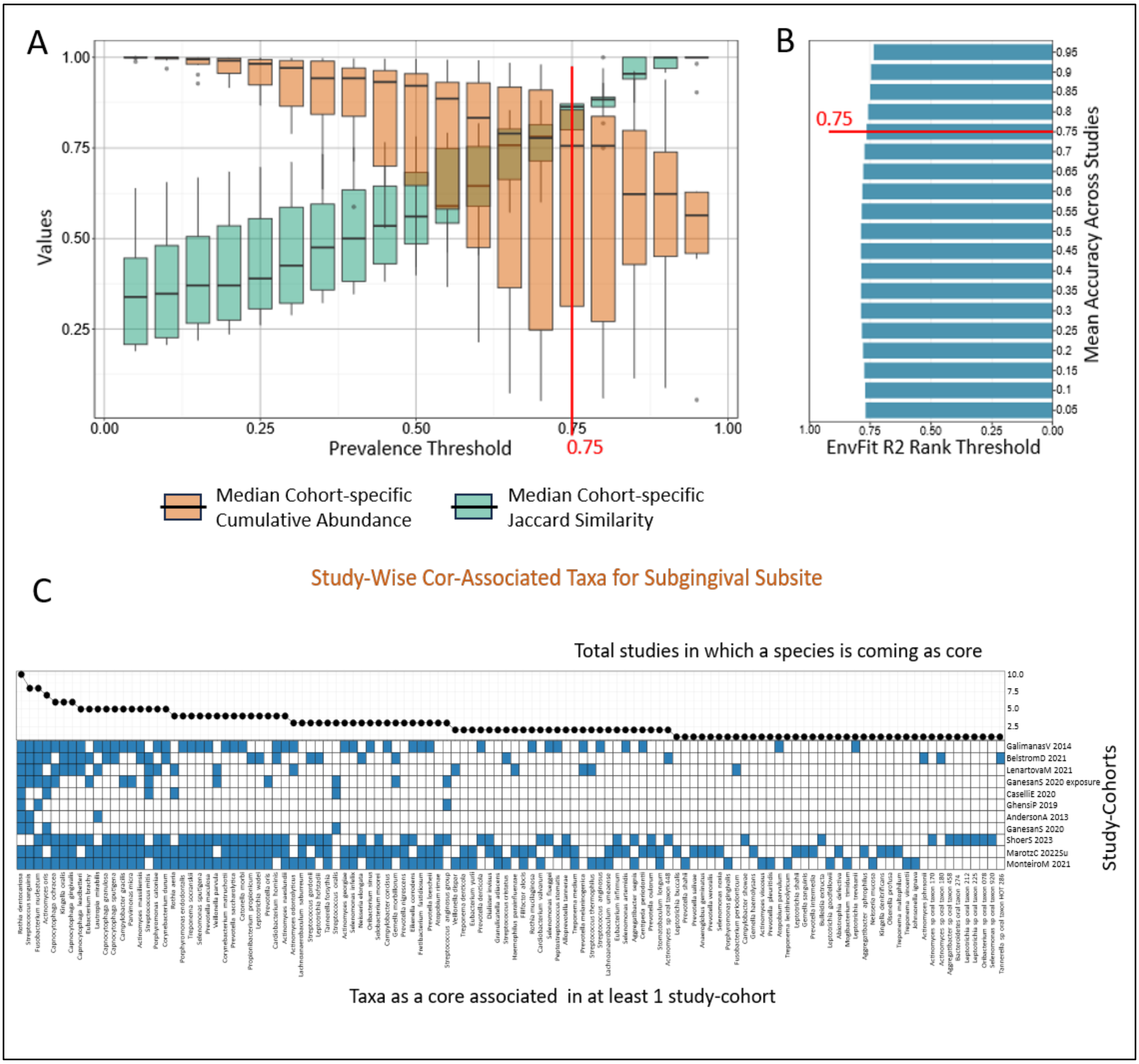


**Figure S19. Threshold optimization of the 3R framework and cohort-specific identification of Core-Associated taxa in the subgingival microbiomes. A**. **Selection of the prevalence threshold for defining candidate core taxa in subgingival control cohorts.** For each prevalence cutoff along the x-axis, taxa detected above that cutoff within a study-cohort were selected, and two properties were then calculated across that cohort's samples: the median Jaccard similarity across all sample pairs (using this taxa subset) and the median cumulative abundance of this taxa subset across samples. These two properties were computed for every study-cohort, and their distributions across cohorts are shown as paired boxplots at each cutoff: light-brown for median cumulative abundance and green for median Jaccard similarity. For low prevalence thresholds, while the identified core taxa would always account for a large proportion of the microbiome, there is going to be high variation in their detection rates across microbiomes. For stringent or high prevalence thresholds, the selected small taxa set would show minimal variation across samples but would also capture only a small proportion of the microbiome. An ideal core-taxa set should be both uniformly represented across samples (high Jaccard similarity) and account for a large proportion of overall microbiome composition (high cumulative abundance); the optimal cutoff is one that jointly maximizes both properties. Candidate prevalence thresholds were evaluated to identify the cutoff of >= 0.75, that retained consistently detected taxa while maintaining broad community representation across supragingival control cohorts. **B**. **Selection of the community-association threshold for identifying taxa strongly linked to subgingival community structure.** Taxa were ranked by $R^2$, and a threshold was chosen to balance two aspects: a threshold set too high would retain only taxa consistently associated with other members' abundances, but would miss taxa with significant community-composition associations that nonetheless have lower ranked $R^2$ (reduced sensitivity); a threshold set too low would misclassify taxa with non-significant associations as ecologically influential (increased false positives). The ideal threshold therefore maximizes accuracy, the proportion of correctly identified taxa with significant community-composition associations. The optimized $R^2$ rank threshold of 0.75 was selected as, for this threshold, we observed the maximum accuracy for identifying taxa whose variation was significantly associated with variation of other members of microbial communities. Finally, the cohort-specific lists of core-associated taxa were generated using a prevalence cutoff of >= 0.75 and $R^2$ rank threshold >= 0.75 using 3R framework. **C. Heatmap showing the identification of core-associated taxa across subgingival control study-cohorts.** Rows represent subgingival taxa and columns represent individual control cohorts. Blue cells indicate that a taxon was classified as core-associated within that cohort after passing the optimized prevalence threshold of 0.75 and rank-scaled $R^2$ threshold of 0.75. Taxa are ordered according to their Core-Association Score, and the top line plot shows their recurrence across cohorts. Only taxa identified as core-associated in at least one study-cohort are shown.

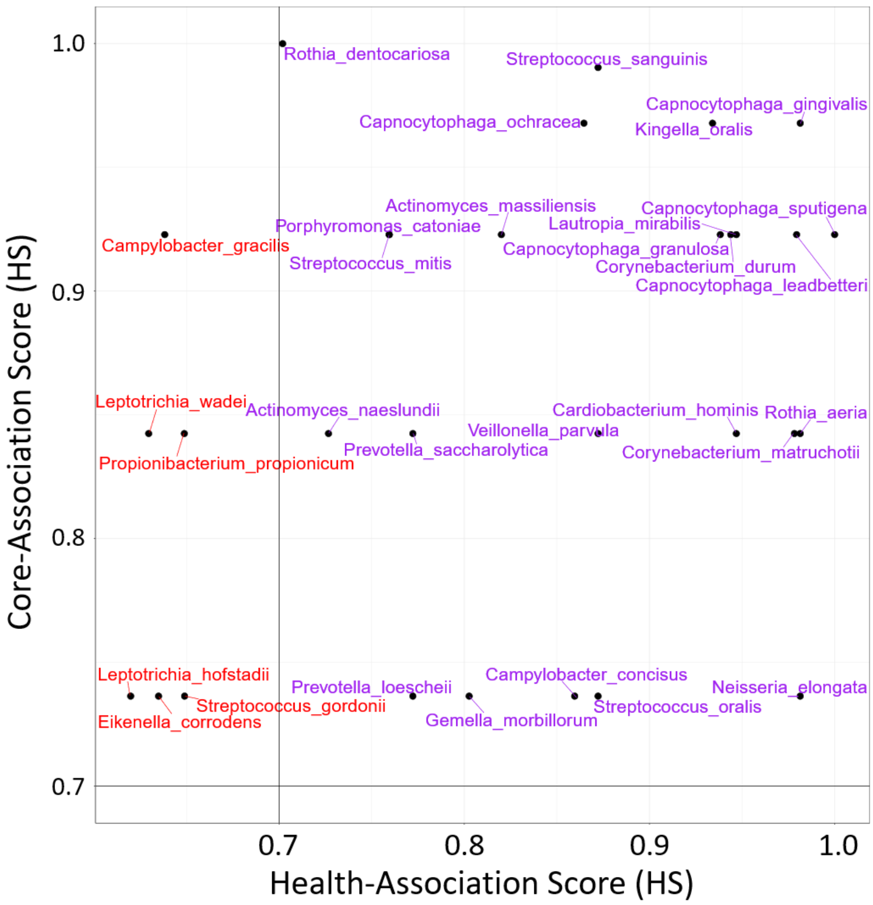


**Figure S20. Scatter plot comparing Core-Association Score (CS) and Health-Association Score (HS) for subgingival taxa**. Only taxa with CS and HS values >=0.60 and are shown. A score threshold of 0.70 was used to stratify taxa into quadrants and identify taxa that were both recurrently core-associated across control cohorts and consistently health-associated across matched case-control cohorts.

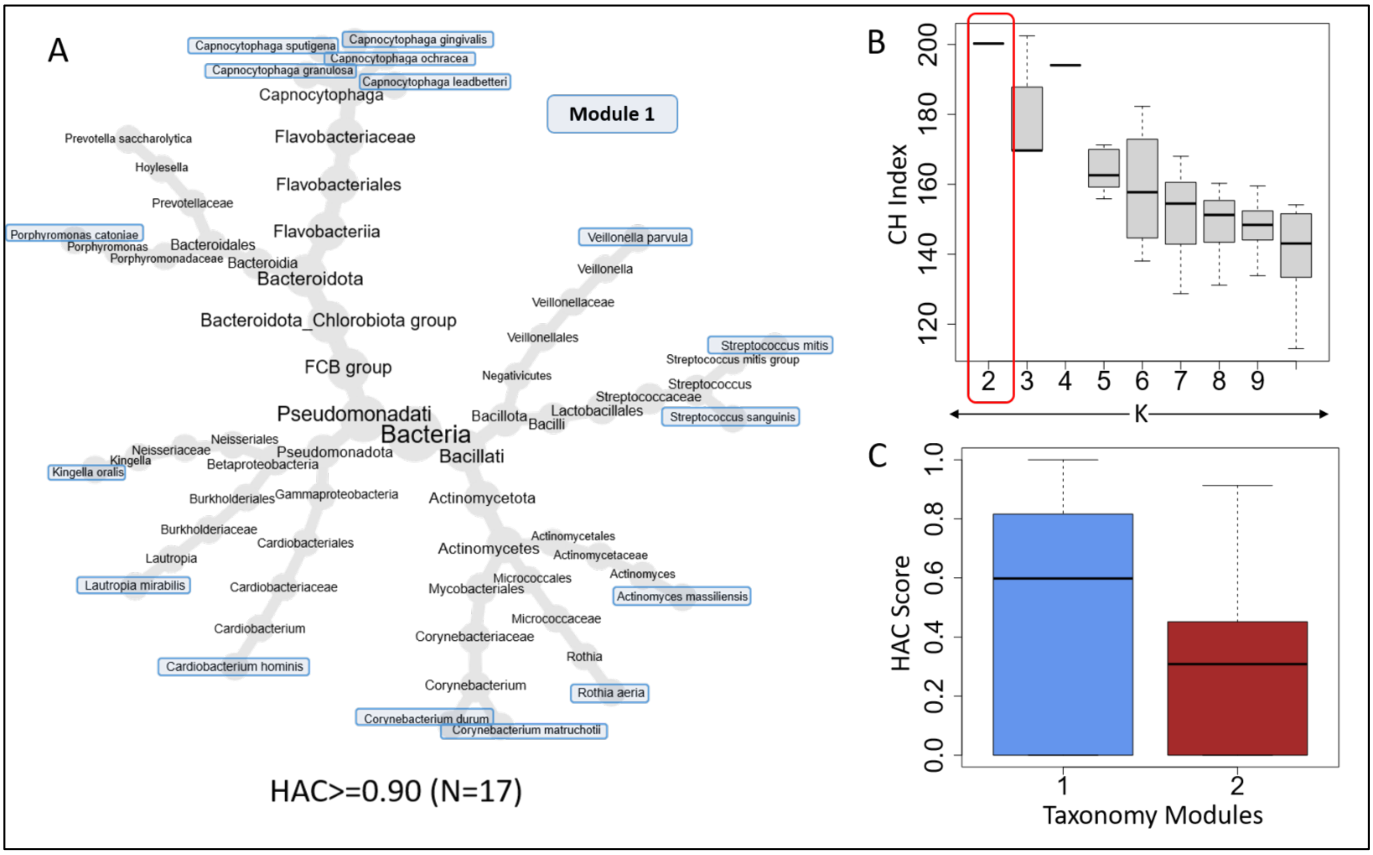


**Figure S21. Identification of the 17 member high-HAC-taxa set (HAC >= 0.90) in the subgingival microbiomes and the high-HAC-enriched microbiome community (subgingival-module-1). A.** Circular phylogenetic tree showing the top 17 high-HAC taxa (HAC >= 0.90) in the supragingival microbiomes, showing their distribution across multiple bacterial lineages. Taxa belonging to taxonomic module 1 (described below, the identification of which is described next) are highlighted in blue boxes. **B.** Selection of the optimal k (=2) used for the identification of subgingival taxonomic modules using iterative k-means clustering analysis. Pairwise association profiles among subgingival consensus taxa were projected into a two-dimensional association space, and a bootstrap based iterative evaluation for various values of k using CH-indices to identify the smallest value of k that gave the best median CH index values across iterations (see **Text S5** and **Figure S15** for methodological details). This analysis selected 2 subgingival taxonomic modules. **C.** Distribution of HAC scores across the three subgingival modules. Module 1 showed noticeably higher HAC values, indicating enrichment of taxa with concordant Health-Association and Core-Association Scores.

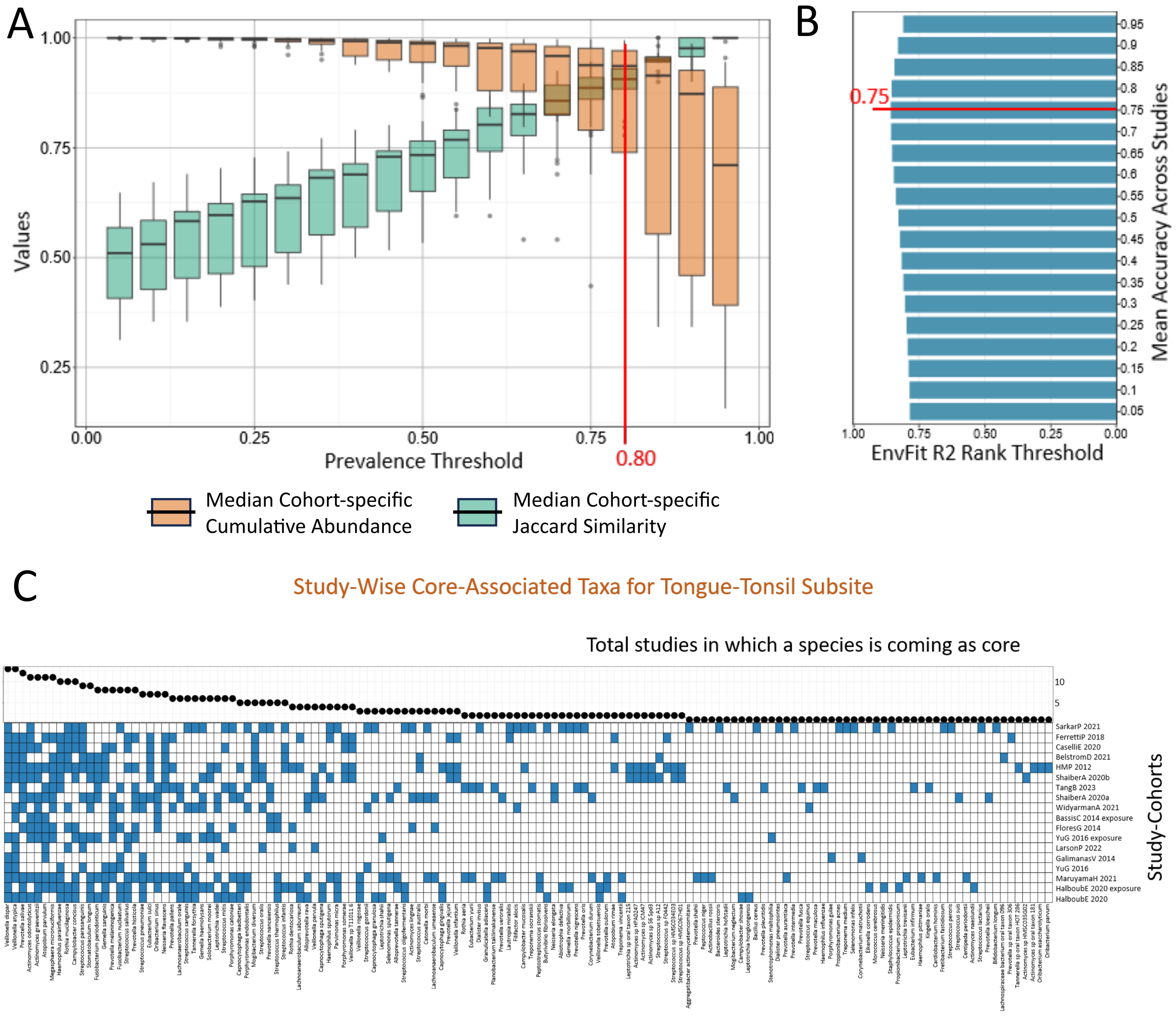


**Figure S22. Threshold optimization of the 3R framework and cohort-specific identification of Core-Associated taxa in the tongue-tonsil microbiomes. A. Selection of the prevalence threshold for defining candidate core taxa in tongue-tonsil control cohorts.** For each prevalence cutoff along the x-axis, taxa detected above that cutoff within a study-cohort were selected, and two properties were then calculated across that cohort's samples: the median Jaccard similarity across all sample pairs (using this taxa subset) and the median cumulative abundance of this taxa subset across samples. These two properties were computed for every study-cohort, and their distributions across cohorts are shown as paired boxplots at each cutoff: light-brown for median cumulative abundance and green for median Jaccard similarity. For low prevalence thresholds, while the identified core taxa would always account for a large proportion of the microbiome, there is going to be high variation in their detection rates across microbiomes. For stringent or high prevalence thresholds, the selected small taxa set would show minimal variation across samples but would also capture only a small proportion of the microbiome. An ideal core-taxa set should be both uniformly represented across samples (high Jaccard similarity) and account for a large proportion of overall microbiome composition (high cumulative abundance); the optimal cutoff is one that jointly maximizes both properties. Candidate prevalence thresholds were evaluated to identify the cutoff of >= 0.80, that retained consistently detected taxa while maintaining broad community representation across tongue-tonsil control cohorts. **B. Selection of the community-association threshold for identifying taxa strongly linked to tongue-tonsil community structure**. Taxa were ranked by $R^2$, and a threshold was chosen to balance two aspects: a threshold set too high would retain only taxa consistently associated with other members' abundances, but would miss taxa with significant community-composition associations that nonetheless have lower ranked $R^2$ (reduced sensitivity); a threshold set too low would misclassify taxa with non-significant associations as ecologically influential (increased false positives). The ideal threshold therefore maximizes accuracy, the proportion of correctly identified taxa with significant community-composition associations. The optimized $R^2$ rank threshold of 0.75 was selected as, for this threshold, we observed the maximum accuracy for identifying taxa whose variation was significantly associated with variation of other members of microbial communities. Finally, the cohort-specific lists of core-associated taxa were generated using a prevalence cutoff of >= 0.80 and $R^2$ rank threshold >= 0.75 using 3R framework. **C. Heatmap showing the identification of core-associated taxa across tongue-tonsil control study-cohorts.** Rows represent tongue-tonsil taxa and columns represent individual control cohorts. Blue cells indicate that a taxon was classified as core-associated within that cohort after passing the optimized prevalence threshold of 0.80 and rank-scaled $R^2$ threshold of 0.75. Taxa are ordered according to their Core-Association Score, and the top line plot shows their recurrence across cohorts. Only taxa identified as core-associated in at least one study-cohort are shown.

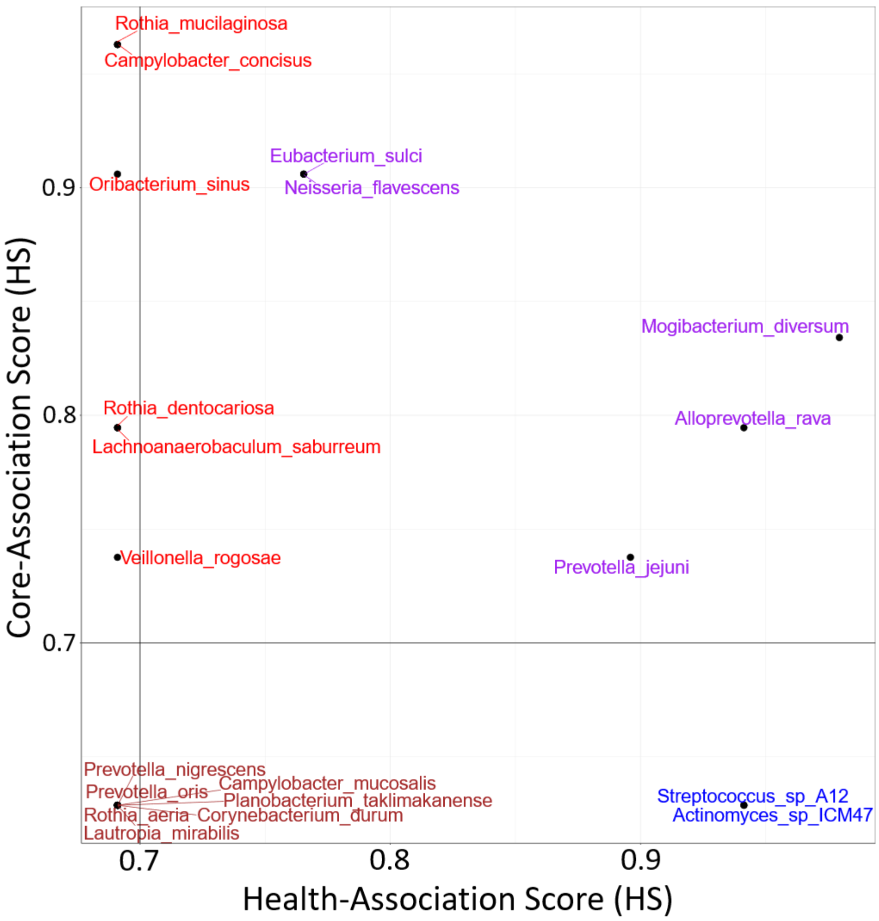


**Figure S23. Scatter plot comparing Core-Association Score (CS) and Health-Association Score (HS) for tongue-tonsil taxa**. Only taxa with CS and HS values >=0.60 and are shown. A score threshold of 0.70 was used to stratify taxa into quadrants and identify taxa that were both recurrently core-associated across control cohorts and consistently health-associated across matched case-control cohorts.

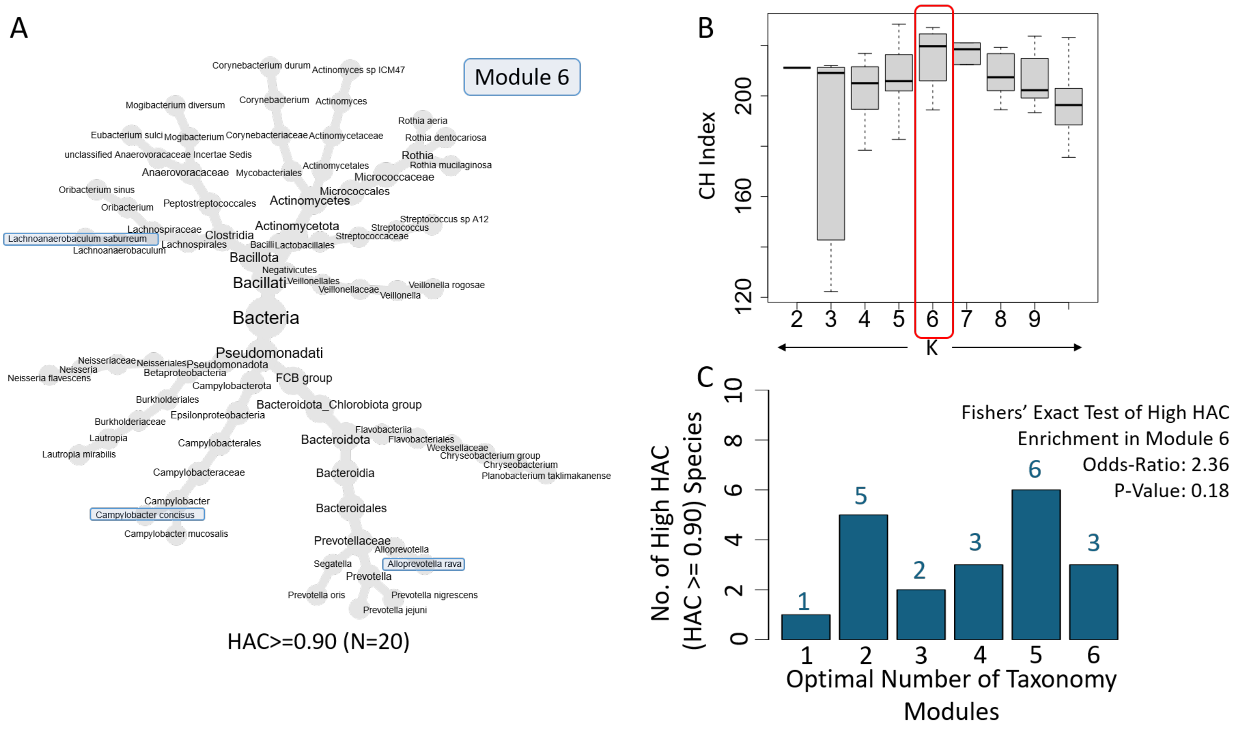


**Figure S24. Identification of the 20 member high-HAC taxa in the tongue-tonsil microenvironment and network-level organization of tongue–tonsil taxonomic modules.** **A.** Circular phylogenetic tree showing the top 20 high-HAC taxa (HAC >= 0.90) in the tongue-tonsil microbiomes, showing their distribution across multiple bacterial lineages. **B.** Selection of the optimal k (=6) used for the identification of tongue-tonsil taxonomic modules using iterative k-means clustering analysis. Pairwise association profiles among subgingival consensus taxa were projected into a two-dimensional association space, and a bootstrap based iterative evaluation for various values of k using CH-indices to identify the smallest value of k that gave the best median CH index values across iterations (see **Text S5** and **Figure S15** for methodological details). This analysis selected 6 tongue-tonsil taxonomic modules. **C.** Distribution of taxa with HAC $\geq$ 0.90 across taxonomic modules in tongue–tonsil subsite. Module 6 showed comparatively higher HAC values in **Figure 6H**, however high-HAC taxa were not significantly enriched in module 6 (odds ratio = 2.36, Fisher's exact test, P= 0.18)

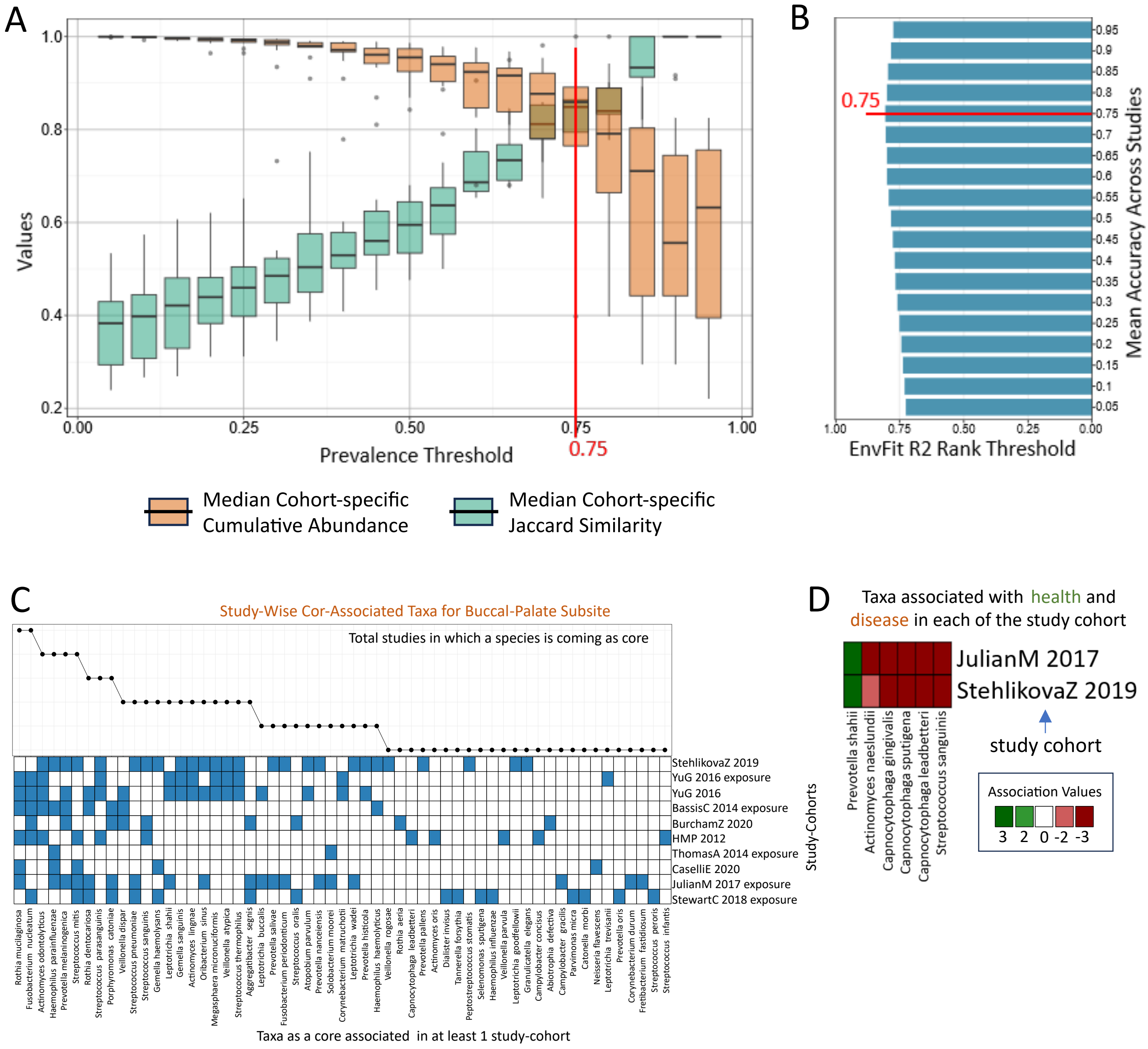


**Figure S25. Core-association and disease-control association patterns in the buccal–palate-other surface subsite. A. Selection of the prevalence threshold for defining candidate core taxa in buccal–palate-other surface control cohorts.** For each prevalence cutoff along the x-axis, taxa detected above that cutoff within a study-cohort were selected, and two properties were then calculated across that cohort's samples: the median Jaccard similarity across all sample pairs (using this taxa subset) and the median cumulative abundance of this taxa subset across samples. These two properties were computed for every study-cohort, and their distributions across cohorts are shown as paired boxplots at each cutoff: light-brown for median cumulative abundance and green for median Jaccard similarity. For low prevalence thresholds, while the identified core taxa would always account for a large proportion of the microbiome, there is going to be high variation in their detection rates across microbiomes. For stringent or high prevalence thresholds, the selected small taxa set would show minimal variation across samples but would also capture only a small proportion of the microbiome. An ideal core-taxa set should be both uniformly represented across samples (high Jaccard similarity) and account for a large proportion of overall microbiome composition (high cumulative abundance); the optimal cutoff is one that jointly maximizes both properties. Based on this investigation, we selected a prevalence threshold cutoff >= 0.75. **B. Selection of the community-association threshold for identifying taxa strongly linked to tongue-tonsil community structure.** Taxa were ranked by $R^2$, and a threshold was chosen to balance two aspects: a threshold set too high would retain only taxa consistently associated with other members' abundances, but would miss taxa with significant community-composition associations that nonetheless have lower ranked $R^2$ (reduced sensitivity); a threshold set too low would misclassify taxa with non-significant associations as ecologically influential (increased false positives). The ideal threshold therefore maximizes accuracy, the proportion of correctly identified taxa with significant community-composition associations. The optimized $R^2$ rank threshold of 0.75 was selected as, for this threshold, we observed the maximum accuracy for identifying taxa whose variation was significantly associated with variation of other members of microbial communities. Finally, the cohort-specific lists of core-associated taxa were generated using a prevalence cutoff of >= 0.75 and $R^2$ rank threshold >= 0.75 using 3R framework. **C. Heatmap showing the identification of core-associated taxa across buccal–palate-other-surface control study-cohorts.** Rows represent taxa and columns represent individual control cohorts. Blue cells indicate that a taxon was classified as core-associated within that cohort after passing the optimized prevalence threshold of 0.75 and rank-scaled $R^2$ threshold of 0.75. Taxa are ordered according to their Core-Association Score, and the left-side line plot shows their recurrence across cohorts. **D. Cohort-wise disease-control association patterns** for buccal–palate-other surface taxa in the two matched disease-control cohorts. Cells indicate taxa classified as control-associated, disease-associated, or non-significant within each cohort. Due to the limited number of matched disease-control cohorts, Health-Association Scores and HAC rankings were not computed for this subsite.

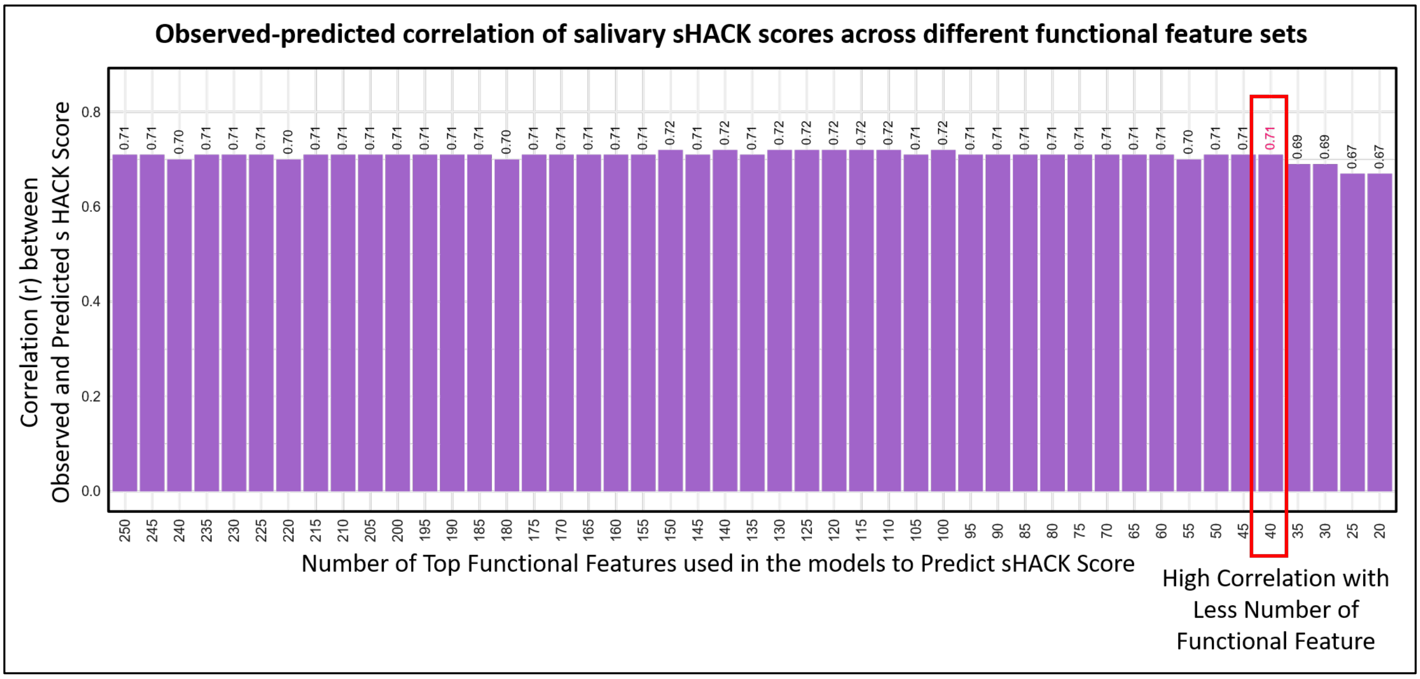


**Figure S26. Identification of the subset of top 40 taxa-level functional features predictive of their salivary-health-associated-core-keystone (sHACK).** Bar plots denoting the out-of-box correlations between the actual sHACK scores (corresponding to 366 taxa) and the sHACK score predicted using Random Forest models using different sized subsets of their top functional features, ranging from 250 to 20 features in decrements of five. The 40-feature model was selected because it retained a comparatively small number of features while achieving a high observed-predicted correlation (Pearson Correlation = 0.71).

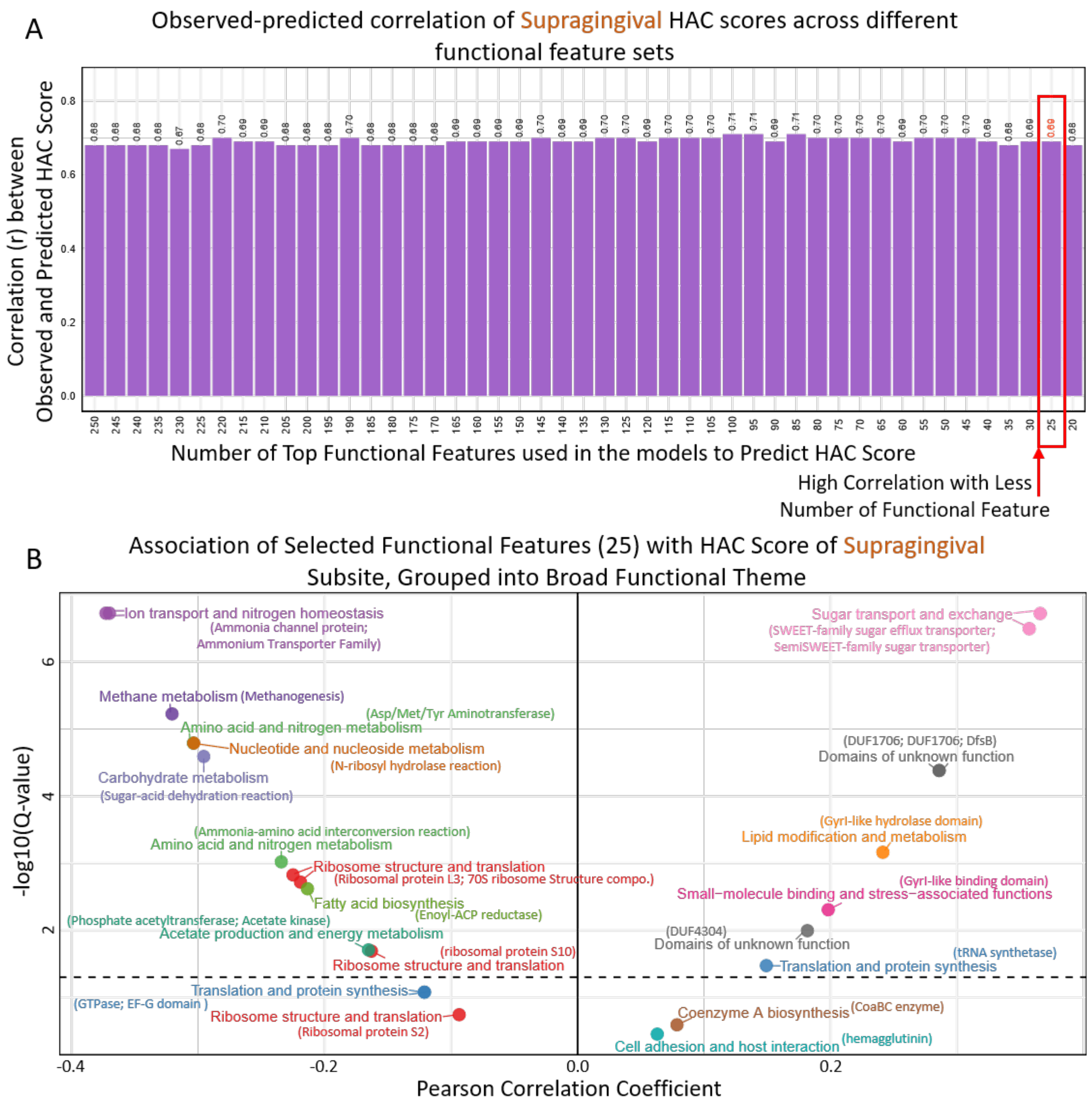


**Figure S27. Identification of top 25 predictive taxa-level functional features and their association patterns with supragingival-HAC scores. A.** Bar plots denoting the out-of-box correlations between the actual supragingival-HAC-scores and HAC-scores predicted using Random Forest models using different sized subsets of their top functional features, ranging from 250 to 20 features in decrements of five. The 25-feature model was selected as that retained a relatively small number of features while maintaining a high observed-predicted correlation (Pearson Correlation = 0.69). **B.** Associations of the top 25 genome-derived functional features with supragingival HAC scores. Volcano-style correlation plot showing the direction and statistical significance of associations between the selected top 25 genome-derived functional features and supragingival HAC scores. The x-axis represents the correlation coefficient, with negative and positive values indicating inverse and direct associations, respectively, and the y-axis represents -log10 of the multiple-testing-adjusted Q-value. The horizontal dashed line indicates the significance threshold at Q= 0.05, and the vertical line indicates a correlation coefficient of zero. Points are coloured and annotated according to their broad functional themes.

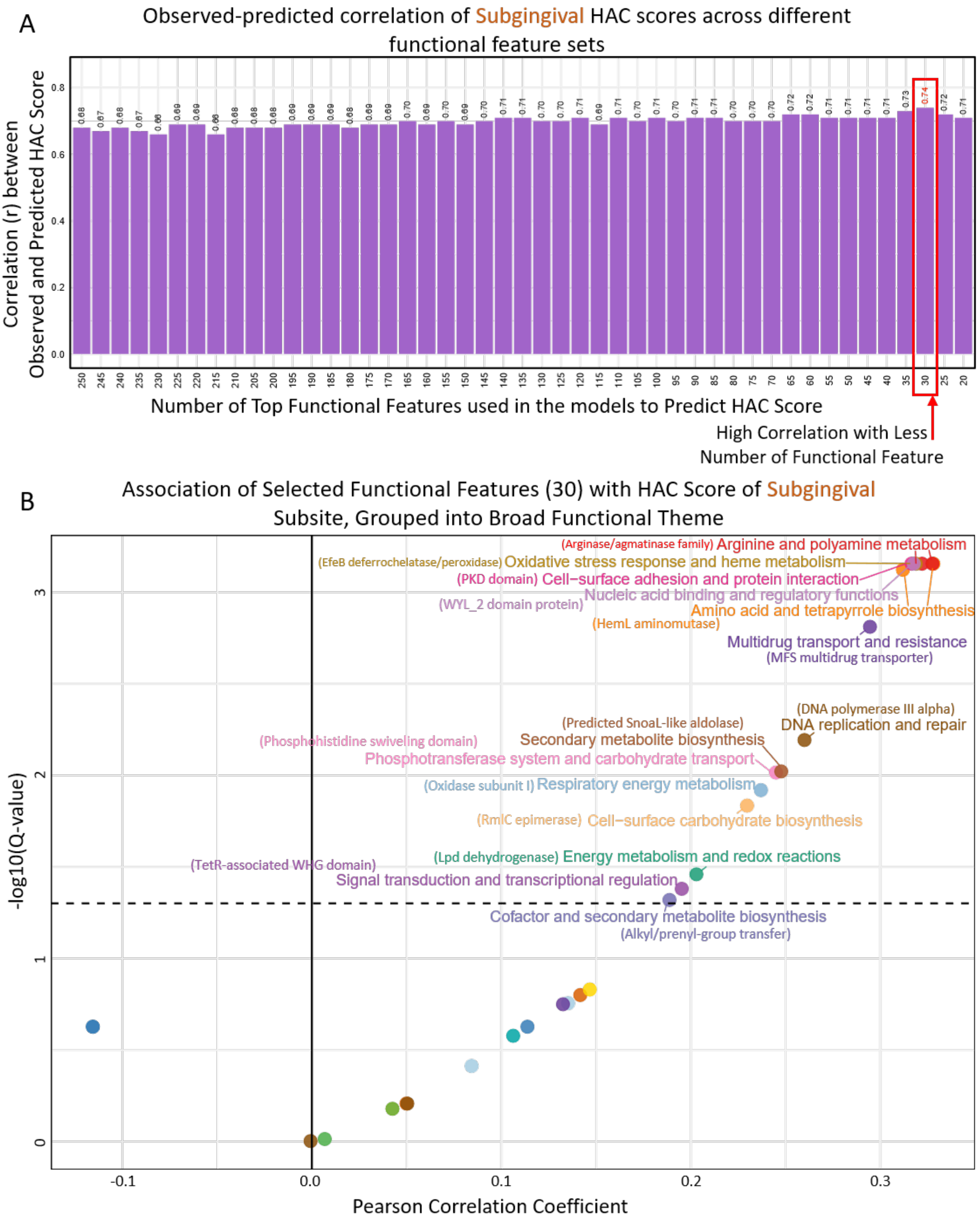


**Figure S28. Identification of top 30 predictive taxa-level functional features and their association patterns with subgingival-HAC scores. A.** Bar plots denoting the out-of-box correlations between the actual subgingival-HAC-scores and HAC-scores predicted using

Random Forest models using different sized subsets of their top functional features, ranging from 250 to 20 features in decrements of five. The 30-feature model was selected as a parsimonious model that retained a relatively small number of features while maintaining a high observed-predicted correlation (Pearson Correlation = 0.74). **B.** Associations of the top 30 genome-derived functional features with supragingival HAC scores. Volcano-style correlation plot showing the direction and statistical significance of associations between the selected top 30 genome-derived functional features and subgingival HAC scores. The x-axis represents the correlation coefficient, with negative and positive values indicating inverse and direct associations, respectively, and the y-axis represents -log10 of the multiple-testing-adjusted Q-value. The horizontal dashed line indicates the significance threshold at Q= 0.05, and the vertical line indicates a correlation coefficient of zero. Points are coloured and annotated according to their broad functional themes.

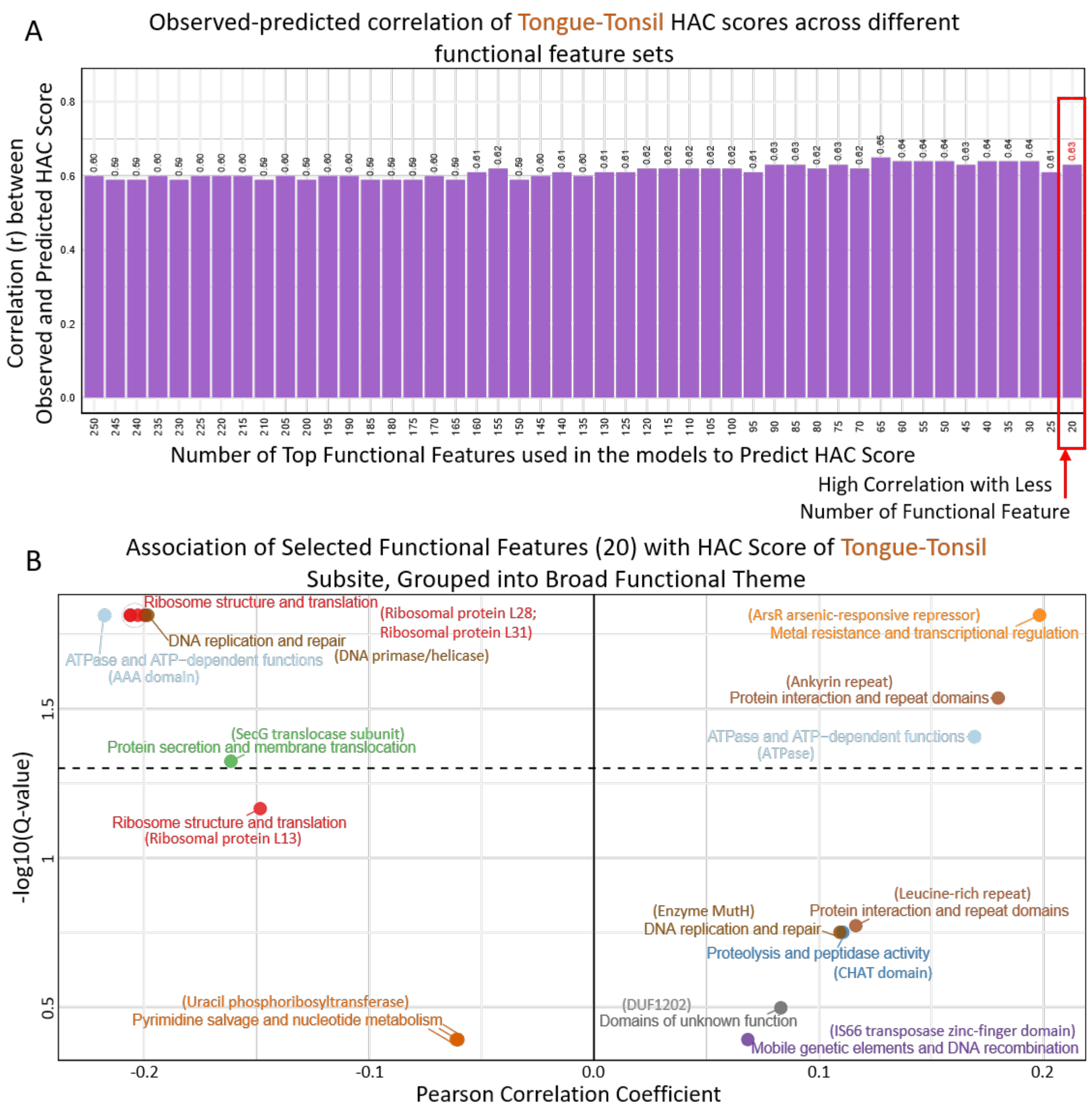


**Figure S29. Identification of top 20 predictive taxa-level functional features and their association patterns with tongue-tonsil HAC scores. A.** Bar plots denoting the out-of-box correlations between the actual tongue-tonsil HAC-scores and HAC-scores predicted using Random Forest models using different sized subsets of their top functional features, ranging from 250 to 20 features in decrements of five. The 20-feature model was selected as a parsimonious model that retained a relatively small number of features while maintaining a high observed-predicted correlation (Pearson Correlation = 0.63). **B.** Associations of the top

20 genome-derived functional features with tongue-tonsil HAC-scores. Volcano-style correlation plot showing the direction and statistical significance of associations between the selected top 20 genome-derived functional features and tongue-tonsil HAC-scores. The x-axis represents the correlation coefficient, with negative and positive values indicating inverse and direct associations, respectively, and the y-axis represents -log10 of the multiple-testing-adjusted Q-value. The horizontal dashed line indicates the significance threshold at Q= 0.05, and the vertical line indicates a correlation coefficient of zero. Points are coloured and annotated according to their broad functional themes.

#### References


1. Goel, A. *et al.* Toward a health-associated core keystone index for the human gut microbiome. *Cell Rep.* **44**, (2025).
